# Fast Bayesian inference of phylogenies from multiple continuous characters

**DOI:** 10.1101/2021.04.21.440863

**Authors:** Rong Zhang, Alexei J. Drummond, Fábio K. Mendes

**Author notes:** Correspondence to be sent to: Department of Biology, Washington University in St. Louis, One Brookings Drive., St. Louis, 63130, MO, United States.

## Abstract

Time-scaled phylogenetic trees are an ultimate goal of evolutionary biology and a necessary ingredient in comparative studies. The accumulation of genomic data has resolved the tree of life to a great extent, yet timing evolutionary events remains challenging if not impossible without external information such as fossil ages and morphological characters. Methods for incorporating morphology in tree estimation have lagged behind their molecular counterparts, especially in the case of continuous characters. Despite recent advances, such tools are still direly needed as we approach the limits of what molecules can teach us. Here, we implement a suite of state-of-the-art methods for leveraging continuous morphology in phylogenetics, and by conducting extensive simulation studies we thoroughly validate and explore our methods’ properties. While retaining model generality and scalability, we make it possible to estimate absolute and relative divergence times from multiple continuous characters while accounting for uncertainty. We compile and analyze one of the most data-type diverse data sets to date, comprised of contemporaneous and ancient molecular sequences, and discrete and continuous characters from living and extinct Carnivora taxa. We conclude by synthesizing lessons about our method’s behavior, and suggest future research venues.

## 1. Introduction

The advent of molecular sequencing has unquestionably revolutionized comparative biology, giving phylogeneticists unprecedented power to recover species relationships and date important evolutionary events (e.g., Zhang et al., 2014; Kawahara et al., 2019; Vanderpool et al., 2020), describe drivers of diversification (Morlon, 2014; Sánchez-Reyes et al., 2017; Condamine et al., 2019), and their relationship with ecologically relevant traits (Goldberg and Igić, 2012; Burin et al., 2016). The accumulation of genomic data further allowed the identification of problems or gaps in molecular evolution models (or their usage; e.g., Sullivan and Swofford, 1997; Kolaczkowski and Thornton, 2004; Mendes and Hahn, 2018), which led to improvements in their realism (Yang, 2006; Rannala and Yang, 2003; Degnan and Salter, 2007), as well as the development of a plethora of computational tools for empiricists wishing to use such models (e.g., Nguyen et al., 2015; Chifman and Kubatko, 2015; Höhna et al., 2016; Zhang et al., 2018; Suchard et al., 2018; Bouckaert et al., 2019).

Despite all progress, abundant genomic sequences and more complex substitution models have not been a panacea for certain challenges in phylogenetics, such as the scaling of phylogenetic trees to calendar years (i.e., “calibrating” trees). Trees measured in absolute time can often be the ultimate goal of evolutionary inference (Philippe et al., 2011), but are also critical ingredients in comparative analyses of morphological and genetic traits, biogeography, and diversification (Felsenstein, 1985; O’Meara, 2012; Uyeda et al., 2018).

There are three main reasons why molecular data and substitution models alone do not provide final solutions to the task of estimating divergence times in absolute scales. First, obtaining non-contemporaneous DNA samples from multiple species for tree calibration is hard at best (Rieux and Balloux, 2016; Cooper and Poinar, 2000; Hagelberg et al., 2015), and impossible in most cases. Second, there is no universal strict molecular clock (Zuckerkandl and Pauling, 1965). Molecular substitution rates have been show to vary among loci, and both among and along lineages (Larracuente et al., 2008; Bromham, 2009; Venditti and Pagel, 2010), and extrapolating them can prove difficult even among closely related species (Besenbacher et al., 2019).

Third, while advances in substitution rate modeling have helped accommodate some substitution rate variation into phylogenetic analyses (Thorne and Kishino, 2005; Drummond et al., 2006; Ho and Duchêne, 2014), popular models do not usually incorporate the correlation of substitution rates with species life-history traits (e.g., organismal size, generation time, fecundity, longevity; Welch et al., 2008; Lanfear et al., 2013) and may support contrasting hypotheses (Duchêne et al., 2014; Beaulieu et al., 2015). More realistic models exist, but their assumptions about how substitution rates evolve (e.g., as a random walk; Lartillot and Delsuc, 2012) might not always hold (Webster and Purvis, 2002; Finarelli and Flynn, 2006), and their implementations can make analyses of large data sets prohibitively slow (Berv and Field, 2018). These complications often lead to phylogenetic trees being reported in lengths of expected substitutions per site – in these “substitution trees”, time and evolutionary rates are conflated.

One way of overcoming some of the aforementioned roadblocks is to include non-molecular heterochronous data in otherwise strictly molecular datasets. The approach known as “tip-dating” does just that, and taxa are placed in the phylogeny according to their heterochronous data (molecular or otherwise). This technique was first employed for divergence time estimation at shorter time scales, in the context of viral phylodynamics (Rambaut, 2000; Drummond et al., 2002), where sample times are usually known with good precision and molecular data can be abundant. When used at macroevolutionary time scales (Pyron, 2011), tip dating has also been dubbed “total-evidence” (“TED”; Ronquist et al., 2012) or “integrative” dating (Ronquist et al., 2016). Placing taxa on trees algorithmically using character data has become a viable alternative to the calibration of node (ages) using fossils (“node dating”; Ho and Phillips, 2009). Earlier studies combined data and analyzed it with methods rooted in parsimony or other heuristics, even if not exclusively (e.g., Grant et al., 2006; Manos et al., 2007; Arango and Wheeler, 2007), while more recent work modeled character evolution probabilistically (e.g., Ronquist et al., 2012; Zhang et al., 2014; Ronquist et al., 2016; Gavryushkina et al., 2017; Ogilvie et al., 2022). For the purposes of the present work, we will explore and discuss the approach taken by the latter subset of studies, and take “total evidence” to mean “probabilistic” total evidence, whereby combined data is analyzed with integrative probabilistic models.

Although TED circumvents some of the shortcomings of node dating, its success also hinges on how often evolutionary model assumptions are met in the wild. Many studies carrying out some form of probabilistic TED (e.g., Pyron, 2017; Gavryushkina et al., 2017; Ogilvie et al., 2022), for example, have relied on the ‘Mk’ and ‘Mkv’ (Lewis, 2001) models (or extensions of them; Pyron, 2017) for discrete morphological characters. This family of models has become popular because, while keeping computational costs under control, it enables the statistical analysis of preexisting (discrete) morphological data sets originally intended for use with parsimony methods. This computational convenience comes at a price, however, and the suitability of these models has received considerable attention (Danforth et al., 2006; Bracken-Grissom et al., 2014; O’Reilly et al., 2016, 2018; Goloboff et al., 2019). Investigations have focused on model assumptions (e.g., discrete characters evolve in uncorrelated fashion), the constraints certain implementations impose (e.g., characters evolve at the same rate), and the appropriateness of the required data preparation protocols (e.g., autapomorphic characters are usually not represented in character matrices).

### Total-evidence with multiple continuous characters

Continuous morphological characters consist of an apropos alternative to their discrete counterpart as a source of phylogenetic information for several reasons. First, statistical models for continuous character evolution are mature and relatively well understood, as a result of a long history of methodological development in this area (e.g., Felsenstein, 1973; Hansen and Martins, 1996; O’Meara et al., 2006; Landis et al., 2013; Bastide et al., 2018; Silvestro et al., 2018; Gaboriau et al., 2020). Second, most phylogenetic methods for studying continuous characters employ models based on the Brownian motion (BM; Felsenstein, 1973), whose formulation (see below) allows them to incorporate among-character evolutionary covariance. Third, continuous characters are usually scored at a resolution that makes them variable across most if not all species, so bias arising from the exclusion of autapomorphic traits is of less concern. Leveraging continuous characters for TED can thus improve inference by injecting phylogenetic information into morphological data sets (see Parins-Fukuchi, 2018b, and Varón-González et al., 2020, for more in-depth discussions), and by providing natural workarounds for issues observed under discrete-character models.

Given the considerations above, it might come as a surprise that continuous-character data sets remain mostly untapped in the context of probabilistic TED. Many computational methods implementing continuous-character models have been developed, but the vast majority of them are geared exclusively toward the study of the evolution of the characters themselves (e.g., Revell, 2012; Pennell et al., 2014; Clavel et al., 2015; Caetano and Harmon, 2017; Mitov et al., 2020), or of the interaction between character evolution and diversification (FitzJohn, 2010). Invariably, these methods require a species tree point estimate or more rarely, a posterior distribution to be available. The purpose of these tools is thus not to infer the phylogeny – phylogenies will have normally been estimated in previous studies using different data sets, often almost strictly molecular (Pyron, 2015). Instead, the phylogeny is treated as observed data, and used in practice to control for common descent (Felsenstein, 1985). This approach can be problematic in the presence of both high levels of phylogenetic uncertainty and of phylogenetic incongruence among molecular and morphological data partitions. In such cases, the morphological data may be analyzed on a phylogenetic “Procrustean bed” (Hahn and Nakhleh, 2016), a species tree that might largely reflect the evolution of the genotype (but ignore that of the phenotype), which can then lead to incorrect evolutionary reconstructions about morphological characters (Mendes and Hahn, 2018).

One alternative is to simultaneously model the evolution of all available data, whereby different data partitions inform on each other’s model parameters and jointly inform the phylogeny itself (Hassler et al., 2023). Efforts in this front are promising, but have so far been shy. Studies of disease spatial phylodynamics (Lemey et al., 2010; Pybus et al., 2012), for example, estimated phylogenetic infection trees together with continuous-character parameters and related quantities. Although implemented in a flexible Bayesian setting, the focus of these methods is phylogeographic and their input was limited to two characters representing latitude and longitude. Furthermore, disease phylodynamic analyses are normally constrained to the short timescales characteristic of viral infections, at which abundant molecular heterochronous data can normally be collected (and then used for node age estimation via the molecular clock; Pybus et al., 2012). Two later, much-needed studies (Lartillot and Delsuc, 2012; Cybis et al., 2015) implemented more general and highly parameterized models capable of leveraging multiple discrete and/or continuous characters, with the important feature of treating character states as covarying quantities. Lartillot and Delsuc (2012) modeled the connection between substitution rates and life-history traits in placental taxa, and reconstructed the ancestral states of those traits jointly with divergence times. Empirical analyses in Cybis et al. (2015), in turn, used the threshold model (Wright, 1934; Felsenstein, 2005) primarily to estimate character correlations. Neither study set out to do TED. In Cybis et al. (2015), many heterochronous molecular samples were available, and in none of their analyses did continuous characters contribute substantially, if at all, to the estimation of relative and absolute node ages; in Lartillot and Delsuc (2012), calibration was attained through node dating (Ho and Phillips, 2009), and the topology was assumed known.

The scope of these previous studies has limited how much headway was made in the context of TED. While illustrating the estimation of phylogenetic trees from continuous characters, they did not attempt, for example, to model within-tip variation nor error in continuous-character measurements (but see Bastide et al., 2020, for a promising recent contribution). They also did not explicitly model the intensity of the process generating heterochronous samples, nor the placement of those samples as ancestor nodes along phylogenetic paths as done elsewhere (Heath et al., 2014; Gavryushkina et al., 2017; Pyron, 2017). One exception has been a recent TED study that analyzed multiple three-dimensional landmarks with a multivariate BM model (Álvarez–Carretero et al., 2019), though the methods therein were limited to node age estimation on a fixed phylogenetic backbone. None of the aforementioned work advancing continuous-character TED methods has extended simple BM to models that can capture evolutionary trajectories more realistically (e.g., Hansen, 1997; Butler and King, 2004; Blomberg et al., 2003). While these methodological efforts have undoubtedly contributed to TED with continuous characters, there is still much to be learned about the factors determining the utility (or lack thereof) of this kind of data, such as fossil abundance, the reliability of the morphological clock, and the impact of model misspecification. Lastly, the actual use and future extensions these tools will see in this context will be largely bounded by the developers’ interests (e.g., disease phylodynamics, trait evolution), as well as by the efficiency, flexibility and user base of both their own and alternative software environments.

Methods for TED from multiple continuous characters have had a slow start likely for several reasons, such as (i) the technical difficulty of implementing and integrating the multiple required model components within the same statistical framework, as well as coupling those with efficient inference machinery, (ii) the prohibitively slow run times due to continuous-character model scalability issues, and (iii) the scarcity of data sets compiling appropriate data (which we expect over time to become less of a hindrance and more of a motivation for method development in this area; Lee and Palci, 2015). To our knowledge there are very few (if any) statistical frameworks with which to readily and efficiently estimate trait-evolution and macroevolutionary parameters jointly with the phylogeny – from both intra- and interspecific data sets comprised of genomic sequences, and multiple discrete and continuous morphological characters from living and fossil taxa. Recent work indicates, however, that there is an increasing demand for (e.g., Silvestro et al., 2018; Cascini et al., 2019; Koch and Thompson, 2020) and ongoing supply of methods capable of integrating such multidimensional data (e.g., Álvarez–Carretero et al., 2019; May and Moore, 2020; Gaboriau et al., 2020; Ogilvie et al., 2022).

Here, we introduce a new method for total-evidence phylogenetic inference from multiple continuous characters. We implement state-of-the-art algorithms that use the most general model formulation currently available (Mitov et al., 2020) – as well as other useful methods (Álvarez–Carretero et al., 2019) – into the flexible hierarchical Bayesian frame-work of the BEAST 2 platform (Bouckaert et al., 2019). As a result, our method is general, extendable, and fast, scaling well with both the number of characters and of taxa. A birth-death model so far absent from the BEAST 2 environment (Stadler and Yang, 2013) is also implemented, to be used as a process underlying the species tree node ages. We then carefully and thoroughly validate all these model components by integrating them into a large hierarchical model, and then use this model in extensive simulations and inferential analyses. In order to better characterize the behavior and utility of our method for TED – and more broadly, the utility of continuous characters – we further explore our integrative model through simulations in a series of different scenarios. Lastly, we compile one of the most diverse total-evidence data sets to date, in terms of data types: molecular sequences, multiple discrete and continuous characters from both living and (sub)fossil Carnivora taxa, and fossil ages. We then carry out phylogenetic inference using this data set and our new method, and analyze, summarize and interpret the resulting reconstructions, while proposing ways of moving forward.

## 2. Materials and M ethods

### 2.1. Modeling continuous characters

The key model we develop in the present work is popularly referred to in comparative biology as the (phylogenetic) Brownian motion model (BM; Felsenstein, 1973). BM and models derived from it have become the main family of models used in phylogenetics for the analysis of continuous-character evolution. These models and their assumptions and properties have been extensively described in the literature (we point the interested reader to Harmon, 2019, and Garamszegi and Møller, 2010, for good introductions and further discussion). Briefly, in its basic phylogenetic formulation, BM treats species continuous characters (each character being represented by a single value per species) as continuous random variables changing over time, along the branches of a phylogenetic tree Φ. More formally, a phylogenetic BM model is a continuous-time stochastic process that describes a collection of successive (random) character-value “dislocations” for all characters and species, according to Φ. In an infinitesimal window of time, these dislocations are drawn from a normal distribution with mean zero and variance that scales with the evolutionary rate *r*.

Phylogenetic BM is a convenient model for the comparative study of continuous characters because its density function is the same as that of the well known multivariate normal distribution:

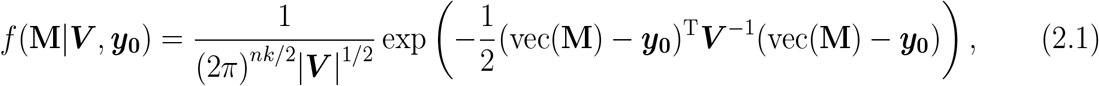

where **M** is an *n × k* matrix of observed continuous characters (*n* species, *k* characters, vec(**M**) denoting the vectorization of **M**), ***V*** is the phylogenetic variance-covariance matrix, and ***y***_**0**_ represents the character values from all characters at the root of Φ. Matrix ***V*** in equation (2.1) is given by the Kronecker product between two matrices: (i) the character variance-covariance matrix **Σ**, and (ii) the ***T*** matrix; that is, ***V*** = **Σ** ⊗ ***T***. Below, we first define **Σ** and then ***T***.

Matrix **Σ** is a *k × k* matrix, defined as 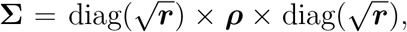, where ***r*** = {*r*_*i*_ : 1 ⩽*i ⩽k*} is a vector containing all relative character-specific evolutionary rates, and:

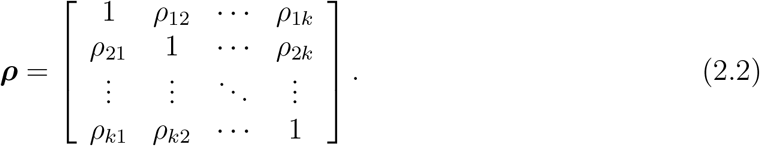

Note that if all character-specific relative evolutionary rates are assumed equal to 1 (i.e.,

***r*** = **1**; Supplementary Fig. 18), then **Σ** = ***ρ***.

Matrix ***T*** is a symmetric *n × n* matrix obtained deterministically from phylogenetic tree Φ, relative branch rates ***b***_*m*_ and global evolutionary rate *c*_*m*_ (subscript *m* indicates these are rates from the morphological model, as opposed to rates from molecular substitution models):

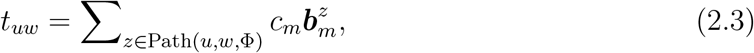

where Path(*u, w*, Φ) returns the set of branch lengths on the phylogenetic path shared by any two species *u* and *w* in Φ (from the root to the most recent common ancestor of *u* and *w*), and 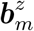 is the relative rate of the branch whose length *z* is being returned by Path(·).

Now that **Σ** and ***T*** are both defined and their dimensions made explicit, one can see that matrix ***V*** (where ***v***_*uw*_ = *t*_*uw*_**Σ**) has dimensions *nk × nk* as a result of the Kronecker product between an *n*^2^ matrix and a *k × k* matrix. Equations (2.2) to (2.3) show that ***V*** can be defined in terms of Φ, ***b***_*m*_, ***r, ρ***, *c*_*m*_. It thus follows that the left side of equation (2.1) can be written as *f* (**M**|Φ, ***b***_*m*_, ***r, ρ***, *c*_*m*_, ***y***_**0**_), a form we will make use of later.

### Intraspecific. variation in continuous characters

As mentioned above, comparative methods traditionally employ a single measurement expected to represent each species in a phylogeny, such as averages from a group of individuals of a species. Individuals within a species, however, will invariably exhibit different phenotypes for a myriad of reasons, such as genetic variability (Lynch and Walsh, 1998), direct effects of environmental factors that differ among populations, variation related to age and sex, seasonal fluctuations (Ives et al., 2007), to name a few. In addition, each data point can be further biased by measurement error due to nonrandom sampling of individuals and instrumental error (Garamszegi and Møller, 2010; Hansen and Bartoszek, 2012). Failing to address intraspecific phenotypic variance can mislead comparative analyses in multiple ways (Kostikova et al., 2016). For example, different modes of evolution can be inferred (e.g., rapid vs. gradual body size changes in vertebrates; Landis and Schraiber, 2017), and both evolutionary rates (Clavel and Morlon, 2017) and divergence times (Álvarez–Carretero et al., 2019) can be overestimated.

One way to account for phenotypic variation among conspecifics is to increment character variances and among-character covariances by some constant **v**_err_ (Ives et al., 2007). This amounts to using phylogenetic variance-covariance matrix 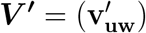, which is updated by **v**_err_ from ***V*** = (**v**_**uw**_) (in Eq. 2.1). We have:

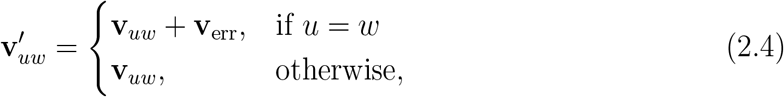

where **v**_err_ is a *k ×k* matrix, defined as **v**_err_ = diag(***σ***)*×****ρ****×*diag(***σ***), and ***σ*** = {*σ*_*i*_ : 1 ⩽*i* ⩽*k*} is a vector containing the intraspecific standard deviation (assumed to be the same for all species) for each character.

If an unbiased estimator of intraspecific character variation, e.g., 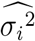 (for variance 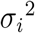), is not available or cannot be computed – in the absence of measurements from multiple individuals from a species – **v**_err_ can be inferred at the cost of longer MCMC chains. When measurements from multiple individuals from a species are available, however, then 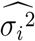 can be straightforwardly calculated for each of the continuous characters, which gives us 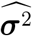.

An additional useful step consists of normalizing each *i*-th observed character in **M** by their corresponding 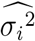, i.e., 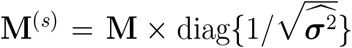, where 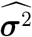 holds the unbiased estimators of intraspecific variance (of all *k* characters). Using **M**^(*s*)^ simplifies calculations because this normalization leads to unit variances and then **v**_err_ = ***ρ***. (***ρ*** can be replaced by its linear shrinkage estimate, ***ρ***^∗^; see the next section and Eq. (2.5) below.)

### Calculating the probability of the data under the model: scalability with respect to the number of characters and taxa

We move now to the task of calculating the probability of the data given the model, *f* (**M**|Φ, ***b***_*m*_, ***r***, *c*_*m*_, ***y***_**0**_). It is clear from equation (2.1) and the definition of ***V*** that computing the probability density of **M** under the BM model for large *n* and *k* will be computationally demanding. Not only is a Kronecker product (***V*** = **Σ** ⊗ ***T***) required, causing the evaluation of equation (2.1) to slow down proportionally to *k*^2^ and *n*^2^, but also ***V*** must be inverted, an operation the lower bound of which is (*nk*)^2^ (Freckleton, 2012).

Fortunately, in the same work proposing BM as an evolutionary model for continuous characters, Felsenstein (1973) also introduced the pruning algorithm as the basis for addressing both problems mentioned above. In a nutshell, the original pruning algorithm amounts to computing three quantities, for each of the (2*n* − 1) nodes in the tree: the variance, the variance-weighted expectation, and the probability density of a multivariate normal distribution given the first two quantities. This algorithm precludes the computation of **Σ** ⊗ ***T*** and the inversion of ***V***, although it is still necessary to invert and calculate the determinant of ***ρ***. This remaining requirement can be ensured possible by either (i) rejecting MCMC proposals causing ***ρ*** to have a determinant of zero, or (ii) reparameterizing **Σ** as **Σ** = ***LL***^*T*^ (an “upper-triangular” decomposition, a factorization similar to Cholesky decomposition; Mitov and Stadler, 2019). We note that in the latter case, the elements of ***L*** (which we would then assign priors to and estimate) are neither character-specific rates (***r***) nor between-character correlation values (***ρ***) anymore, though such parameters can still be easily recovered by post-processing the MCMC chain. Alternatively, inverting ***ρ*** can be avoided altogether by transforming the data, **M** (Eq. 6 in Álvarez–Carretero et al., 2019). Both these options are possible with our implementation, and elaborated on in the supplement. For the sake of brevity, and because this algorithm has been described and generalized in many subsequent studies (e.g., Felsenstein 1973; Pybus et al. 2012; Freckleton 2012; Caetano and Harmon 2017; Álvarez–Carretero et al. 2019; Mitov et al. 2020), we point the interested reader to those references and to our supplementary material for more detailed descriptions of the algorithm and a worked example.

More recently, Mitov et al. (2020) proposed a very general pruning-based solution for calculating equation (2.1), as well as the probability density function of more general models, such as BM with early bursts (Harmon et al., 2008) and accelerating or decelerating rates (Blomberg et al., 2003), BM with trends (Hansen and Martins, 1996) and the Ornstein-Uhlenbeck process (Hansen, 1997; Butler and King, 2004). Unlike the pruning algorithm by Felsenstein (1973), the algorithm in Mitov et al. (2020) does not compute the maximum-likelihood estimate of character values at internal nodes of the tree, but instead calculates a series of intermediate values (which gives this algorithm its flexibility; see Eq. 2 in Mitov et al. 2020). These intermediate values are then combined at the root node in the calculation of an integral, which then gives the final probability density (Eq. 6 in Mitov et al. 2020). Readers can find a detailed description of this algorithm in Mitov et al. (2020), with it being put to use in Mitov and Stadler (2019). We provide a worked example in the supplementary material.

The second obstacle to carrying out inference for multiple characters under phylogenetic BM is posed by the curse of dimensionality. As *k* increases, the number of character correlation parameters (the off-diagonal elements of ***ρ***) we must estimate increases quadratically; for Bayesian inference, this means long MCMC chains must be employed in order to achieve convergence. Furthermore, unless the number of taxa *n* also increases in a similar fashion, there will be more parameters to estimate than data points (i.e., *k > n*). The larger *k* is relative to *n*, the less informative will each observation be, generally, to the extent that the posterior distribution about one or more parameters may become largely or effectively indistinguishable from the prior. Pathologically, as *k >> n*, the determinant of ***ρ*** approaches zero and this matrix becomes singular and non-invertible. As a result, **Σ** cannot be inverted and then the likelihood of the phylogenetic BM model cannot be evaluated (Eq. (2.1)). (We point the interested reader to the supplement, where we show this issue manifests precisely when (*n* − 1) *< k*.)

Clavel et al. (2019) recently addressed the *k > n* issue by borrowing a technique from the regularization/shrinkage statistical literature and using it in the context of penalized maximum-likelihood estimation. One of the penalties these authors investigated, the linear ridge penalty, allows one to arrive at the “archetypal ridge estimator”:

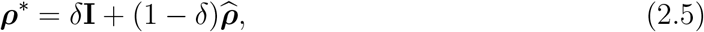

which consists of the average between the *k* × *k* identity matrix (though other matrices can be used) and 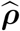 (an unbiased estimator of ***ρ***; see below), weighted by the shrinkage (a.k.a., tuning) parameter *δ*. This parameter can be optimized as described in Schäfer and Strimmer (2005) (more details can be found in the supplementary material). Later, Álvarez–Carretero et al. (2019) used this same linear shrinkage estimator in a Bayesian context.

As summarized above, there are obvious challenges involved in learning about the evolution of morphological character correlations (e.g., Goswami et al., 2014; Caetano and Harmon, 2017), but recent work suggests they are not insurmountable (Clavel et al., 2019). Additionally, although the non-independence among characters should always be accounted for in some manner, it might be of secondary interest relative to the estimation of a time-scaled phylogenetic tree. Different approaches have been explored or suggested for such cases (e.g., Adams, 2014; Goolsby, 2016; Adams and Collyer, 2018), but one simple solution is to employ an unbiased estimator of ***ρ***, 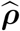, obtained from multiple characters observed across multiple individuals within a species in the phylogeny. Note that here this estimator is unbiased with respect to the population of a single species, and by using 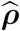 we are assuming character correlations are the same across species and over time.

### 2.2. Tree models for heterochronous samples

As opposed to ad hoc techniques like node dating, whereby experts deliberately attach heterochronous samples to nodes in the phylogeny, in TED we must instead allow the data to guide the placement of those samples. In the case of fossils, morphological characters and the underlying morphological models play key roles in informing fossil affinity. Moreover, we would like to consider the statistical uncertainty around the phylogenetic placement of samples, which requires calculating the probability of the data given different node configurations. Hence, we need a stochastic model capable of generating phylogenetic trees with different nodes and times. The fossilized birth-death process (FBD; Stadler, 2010) is one such model that has received considerable attention in TED due to its ability to account for fossilization, speciation, extinction and taxon sampling (Heath et al., 2014; Gavryushkina et al., 2014a; Zhang et al., 2016). The birth-death-sequential-sampling model (BDSS; Stadler and Yang, 2013) is yet another option; unlike its statistical cousin in the FBD process, the BDSS model conditions on the left-to-right orientation of tip times.

### 2.3. Putting it all together: the integrative model

Carrying out probabilistic TED requires integrating the models mentioned above, as well as other model components, under a coherent statistical framework. This can be done by building a hierarchical integrative model in a Bayesian setting, which can be expressed as the product of the probability density and mass functions of its several component sampling distributions:

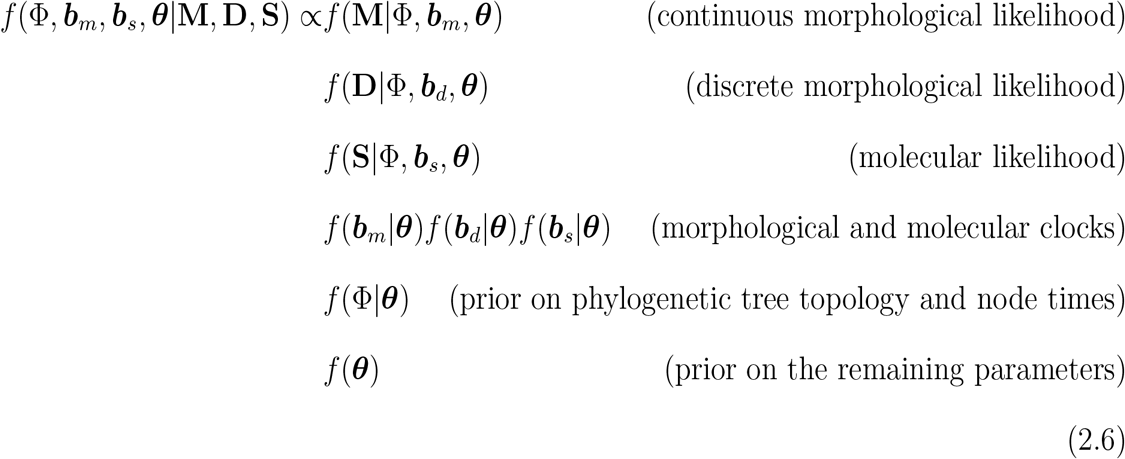

This joint posterior probability density gives the posterior distribution of the time-scaled phylogenetic tree (Φ), morphological and molecular relative branch rates (***b***_*m*_, ***b***_*d*_, ***b***_*s*_) and all remaining parameters (***θ***) – given continuous and discrete morphology data matrices, **M** and **D**, respectively, and molecular sequence alignment **S**.

The continuous morphological likelihood corresponds to the probability of observing **M** under a phylogenetic BM model (Felsenstein, 1973), as described above. The discrete morphological likelihood gives the probability of observing **D** given Lewis’ Mkv model (Lewis, 2001), and the molecular likelihood the probability of observing **S** under a molecular substitution model (Felsenstein, 1981). Finally, the tree prior *f* (Φ|***θ***) gives the probability of a specific topology and node times in phylogenetic tree Φ, with *f* (***θ***) corresponding to the prior distribution on all remaining parameters (see Supplementary Table S14 and Supplementary. Fig. 18 for a description of all parameters).

The posterior distribution *f* (Φ, ***b***_*m*_, ***b***_*d*_, ***b***_*s*_, ***θ***|**M, S**) under our integrative model is approximated by Markov Chain Monte Carlo (MCMC) sampling in BEAST 2 (Bouckaert et al., 2019).

### 2.4. Implemented software

Here, we implement a general, state-of-the-art pruning-based algorithm (Mitov et al., 2020) for phylogenetic Brownian models under a Bayesian framework, as part of BEAST 2’s (Bouckaert et al., 2019) `contraband` package (https://github.com/fkmendes/contraband). We integrate this phylogenetic BM implementation with other methods (Álvarez–Carretero et al., 2019) implemented here, and with several pre-existing model units and features of BEAST 2, such as discrete morphological and molecular models, clock models, parametric distributions, the FBD model, efficient MCMC moves, to name a few.

This integration involved extensive benchmarking and validation work, attained through large simulation experiments (see below and the supplementary material). Under our implementation, the among-character covariance can either be sampled (as in Caetano and Harmon 2017) or its unbiased estimator can be used (as in Álvarez–Carretero et al. 2019). For cases where the number of characters is near to or larger than the number of species, computing the linear shrinkage estimate of the character variance-covariance matrix is also available as an option.

Lastly, we implemented and validated the BDSS model for its utility both as an alternative to the FBD tree prior in TED, and as a necessary component for validation. Validating our BDSS implementation required us to derive equations for, and implement a direct BDSS simulator (to our knowledge, the first of its kind). Details on the BDSS model and the simulator can be found in the supplementary material.

### 2.5. Simulation for assessing method correctness, accuracy, precision and speed

We performed four large simulation studies that together cover three tasks: (a) verify that our newly implemented model is correct and that parameters can be learned from synthetic data, (b) ascertain under what conditions our method is useful to TED, and (c) measure its computational efficiency against implementations elsewhere. We leave fully detailed accounts on the technical settings of all studies to the supplementary material. The fourth study, concerned exclusively with goal (c), is also delegated to the supplementary material.

#### Simulation study I

The overall objective of this experiment was to show that our integrative model is correct and that parameters can be estimated with good accuracy. This experiment can be seen as a large integration test, and it proceeds in three stages. First, we simulate synthetic data from an integrative model. Second, we perform inference under the very same model, using the data simulated in the first step. In our case, inference amounts to executing MCMC analyses, and then logging and summarizing the resulting posterior distributions over the model parameters. Third, for each parameter we calculate the coverage of a given highest-posterior-density (HPD) interval. A model that is correct, or “well-calibrated”, is characterized by appropriate coverage; e.g., if 95%-HPD intervals are used, coverage should be around 95%, that is, the true (simulated) parameter value is expected to be contained in the interval close to 95% of the time. In addition to coverage, the posterior mean of each parameter can be plotted against the corresponding true value; if these two quantities are highly correlated for a given parameter, one can conclude that such parameter is identifiable, and that the method has good accuracy (at least for the size of the simulated data set).

The integrative model we used in this simulation experiment is similar to that defined in equation (2.6), but without discrete morphological and molecular data. Briefly, we used an FBD tree prior, a phylogenetic BM model for four correlated continuous characters, and a strict morphological clock. In total, we simulated 100 data sets each unique in their combination of parameters and data, and then followed the four steps described above. Details about MCMC analyses are presented in the supplementary material.

#### Simulation study II

The second simulation study was designed with the general purpose of characterizing some of the conditions that make continuous characters useful to TED. Our goal was to assess the accuracy and precision of divergence time estimation as a function of the number of continuous characters and fossils, as well as of the interaction between character number and (i) character correlation, (ii) fossil phylogenetic location, and (iii) fossil age. We paid attention to the accuracy of root and average node divergence time estimates, as reflected in the absolute difference between the true (simulated) and inferred (posterior mean) node times. Precision was measured as the width of 95%-HPD intervals.

For five different numbers of continuous characters (1, 2, 5, 10, and 50) we inferred node divergence times from uncorrelated characters and highly correlated characters (simulated with *ρ* = 0.0 and *ρ* = 0.9 for all pairs of characters, respectively), but always assuming character independence. For this analysis, we used the tree shown in supplementary figure 9a; average divergence time estimates here were calculated over 18 internal nodes excluding the root, which was analyzed separately. We also investigated the effect of fossil phylogenetic location on divergence time estimates. We first compared root age estimates when leveraging 18 randomly chosen fossils (9 on each side of the root) to estimates using only fossils (18 of them) from one side of the tree shown in supplementary figure 9b. Second, using the same aforementioned tree, we compared inferred node ages (averaged over 48 nodes) from one side of the tree where all fossils were removed, with node ages on the opposing side (again averaged over 48 nodes) keeping all fossils. Third, we examined how fossil age affects the inference of divergence times. Here, we compared divergence time estimates using the oldest half of all fossils as opposed to the youngest half, again in the tree shown in supplementary figure 9a. Unless described otherwise, all above experiments assumed a known *ρ* of 0.9.

In the above, the evolutionary rate was co-estimated with the other parameters and assumed to be log-normally distributed. Each simulated evolutionary rate was shared by all branches in the phylogeny (i.e., a strict morphological clock). The two phylogenies used in the analyses described above (Supplementary Fig. 9a,b) are arbitrary samples from an FBD process and were assumed partially known: their topologies (but not branch lengths) descending directly from extant taxa were fixed to the simulated truth. Extinct lineage and direct-ancestor fossil placements were estimated, but those nodes and branches were pruned off before divergence time estimates were parsed.

As done elsewhere (Revell et al., 2015; Parins-Fukuchi, 2018a), we assumed the extant tree topology was known because thousands of loci are increasingly available for multiple, mostly extant taxa, and are often strongly informative of how these taxa are related. This strong assumption also greatly improved our ability to interpret model behavior as a function of data set features, while still allowing us to estimate many divergence times (and other parameters), repeatedly, within reasonable CPU hours. More importantly, keeping the topology constant (and pruning fossil tips and lineages and direct ancestors) meant that we had a common set of internal nodes among the trees we used for simulation and the phylogenetic posterior samples. These are the nodes whose estimated ages we averaged over. We must note, however, that while for our purposes holding part of the phylogeny constant felt justified, relaxing such assumption whenever possible is of course recommended in empirical analyses, and could also be done in future simulation work.

We did not jointly estimate all parameters for several reasons. First, simulation study I (described above) already conducted inference integrating over the uncertainty about all parameters, confirming model correctness. Second, for simulation study II, our focus was to distill the effect that specific data features can have on the task of dating phylogenies. Examining multiple data features and parameters at once can quickly obscure otherwise clear connections between data and model. Third, the number of character correlation parameters increases quadratically with the number of scored characters – meaning increasingly larger phylogenies are required for accurate inference. Not only do large trees lead to prohibitively slow analyses, but it is also unclear how to choose those phylogenies. So given the large number of replicated experiments we wished to carry out, we reasoned that constraining parameter space was a logical, if not unavoidable, step in our protocols.

All the above analyses contrasted two conditions (e.g., the off-diagonal elements of ***ρ*** fixed to 0.0 or 0.9; “Young” vs. “Old” half of the fossils, etc.), both replicated 20 times (i.e., a total of 40 replicates) for each number of characters. (Note that a total of only 20 replicates were generated for simulations with a single character.) This amounted to a grand total of 580 simulation and MCMC procedures. Details about MCMC analyses are presented in the supplementary material.

#### Simulation study III

The third simulation study was concerned with a different set of morphological data features with the potential to improve the accuracy or precision of divergence time inference. Namely (i) the number of fossils in a data set (with fossils being pruned from the tree in Supplementary Fig. 9c, and being scored for ten characters), and (ii) adherence of the data to phylogenetic BM model assumptions. We also compared different methods for accounting for character correlation. As in simulation study II, accuracy was measured as the absolute difference between true and inferred (posterior mean) node ages, and precision as the width of 95%-HPD intervals. Again, we focused on the root age and on the average node age.

Our simulation protocol was similar to that in Münkemüller et al. (2012) for obtaining continuous characters with increasingly better fit to phylogenetic BM. Multiple characters (either two or ten; Fig. 3d-f) were simulated as:

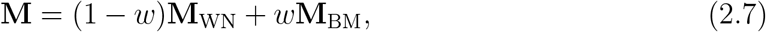

where **M**_WN_ and **M**_WN_ are matrices of character values simulated on the same phylogeny (Supplementary Fig. 9b), under a white noise (WN) and phylogenetic BM models. (Note that under the WN model, the tree topology is irrelevant.) The same evolutionary rate sampling distribution (a log-normal with mean 0.0 and standard deviation 0.1, both in logspace) was used in simulation and as the prior in inference, under a strict morphological clock. Character correlation was kept constant and assumed known (*ρ* = 0.9 for all pairs of characters). We inferred node ages under phylogenetic BM for different values of *w*, the weighing factor dictating the fit between the simulated data and the character model. A *w* of 1.0 indicates a data set simulated exclusively under BM, and thus a perfect model fit.

We compared the performance of alternative methods for accounting for character correlation under trees of different size.While holding the tree topology constant, we pruned an increasingly larger number of extant taxa from a 50-leaf tree (Supplementary Fig. 9d), starting from 35 (15 tips remaining) and stopping at five (45 tips remaining). So as to ensure valid parameter values for simulation and inference, we simulated ten characters using the upper-triangular decomposition mentioned above. The diagonal elements of ***L*** were drawn from a log-normal with mean 0.0 and standard deviation 0.1 in log-space, and the off-diagonal elements from a normal distribution with mean 0.5 and standard deviation 0.5, truncated at -1.0 and 1.0. (***ρ*** was obtained deterministically from **Σ** and ***L***, and used for simulating characters varying within-species; see (iii) below.) These two distributions were correctly specified in inference, during which we either (i) assumed characters to be independent (i.e., the off-diagonal elements of ***ρ*** were set to 0.0; we referred to this analysis as “Independent”), (ii) estimated ***L*** (the “MCMC” analysis), or (iii) approximated ***ρ*** using its linear shrinkage estimate (the “***ρ***^**∗**^” analysis).

As in simulation study II, we replicated data set simulations multiple times for each value of an investigated data feature. When estimating divergence times as a function of the number of fossils or extant taxa in the tree, each data feature value was replicated 25 times, amounting to 200 replicates (100 for panels (a-c) and 100 for panels (g-i) in Fig. 3). For each level of fit to phylogenetic BM (*w*) considered, we generated 20 replicates, to a total of 200. Simulation study III thus yielded 400 different replicates overall. We note that the parameter space investigated here was again partially constrained, for the same reasons outlined above. Details about MCMC analyses are presented in the supplementary material.

### 2.6. Case study: Carnivora phylogeny

#### Molecular and morphological Carnivora data

Our Carnivora data set includes molecular and morphological data from 27 species (see Supplementary Table S11), with 16 of those taxa being extant and 11 extinct. We collected the NCBI sequences of 12 mitochondrial genes from all extant species and the extinct *Smilodon populator* (see Supplementary Table S12). Sequences were concatenated, aligned using MAFFT (Katoh and Standley, 2013), and then partitioned into first-and-second codon positions versus third codon positions (7,320 and 3,660 sites, respectively).

Discrete morphological data from ten taxa (three of which are extinct) was retrieved from Barrett et al. (2021). These consisted of 227 characters partitioned according to their number of states (two, three or four). This partitioning scheme was determined to be the best by Barrett et al. (2021). Continuous morphological data was obtained from Álvarez– Carretero et al. (2019), and consisted of 29 cranium landmarks scored for 19 taxa, each landmark comprising three (continuous) coordinates. Our continuous data set thus contained 87 characters.

Continuous characters like those in our data set have to be aligned prior to analysis. In addition to size and shape, raw landmarks carry nuisance information about position and orientation, which preclude their statistical analysis (Mitteroecker et al., 2013). Distilling shape and size from raw landmarks can be done with Procrustes superimposition, commonly used in biological shape analysis to “align” (superimpose) landmarks (Mitteroecker et al., 2013). Procrustes superimposition consists of rotating, translating and scaling landmark configurations relative to their centroid (i.e., their average position) and its size, so as to minimize the Procrustes distance – a measure of how different in shape two landmark configurations are (Gower, 1975; Rohlf and Slice, 1990). The Procrustes distance is given by the summed squared distance over landmarks and their sample average position; if zero, then two landmarks have the same shape.

Lastly, in order to control for character variation within species, we normalized each landmark in all species by the square root of their corresponding unbiased estimate of intraspecific trait variance, i.e., 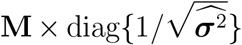. Variance estimates for all landmarks, 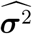, were calculated from landmark data from 21 individuals of *V. vulpes*. Both landmark “alignment” and normalization protocols are further described and used in Álvarez–Carretero et al. (2019).

#### Comparison of cranium landmarks using multidimensional scaling

In addition to using continuous characters in phylogenetic inference, we quantitatively explored cranium landmark data with multidimensional scaling (MDS). MDS is a data exploration technique that projects some measure of similarity (also referred to as “proximity”) among high-dimensional data points onto a low-dimensional spatial map (Mugavin, 2008; Jaworska and Chupetlovska-Anastasova, 2009). This projection is done through a transformation function that preserves essential information (Kruskal and Wish, 1978; Borg and Groenen, 2005). By looking at the spatial map, one then hopes to better grasp how data points compare to each other; if points cluster together, for example, that indicates they share some characteristic.

With result visualization in mind, we carried out MDS by choosing the transformation function that allowed for the easiest compression of the high-dimensional data onto a 2-D spatial map (i.e., the function that minimized the so-called “stress” statistic; Kruskal and Wish 1978). MDS was conducted on the Euclidean distances among the 19 Carnivora species scored for cranium landmarks, across all superimposed landmarks. We used the mds() subroutine from the `smacof` R package (Mair et al., 2019).

#### Integrative probabilistic model for TED

The general mathematical form of the integrative model used in the Carnivora data analysis is shown in equation (2.6), and a model graphical representation is presented in the supplement (Supplementary Fig. 18). We describe all prior distributions used in our analyses in supplementary table S14.

We modeled speciation, extinction and fossilization with either the fossilized birth-death (FBD; Gavryushkina et al., 2014b) or birth-death-sequential-sampling (BDSS; Stadler and Yang, 2013) processes. Extant tip sampling was assumed random, i.e., the probability of sampling extant taxa was a free parameter in our model. Phylogenetic inference of divergence times was carried out while either (1) co-estimating the topology (i.e., no topological constraints), or (2) constraining the monophyly of Carnivora, the *Smilodon* genus, Hyaenidae (hyaenas) and Erinaceidae (hedgehogs and moonrats) families, the latter being used as the outgroup, or (3) constraining Carnivora, the *Caniformia* and *Feliformia* suborders and the Erinaceidae family. We conducted nested sampling (Table 1) to obtain and then compare the fit of different model permutations to Carnivora data.

**Table 1.** Six different integrative model configurations used to estimate the Carnivora phylogeny. The name of the analysis indicates which tree prior was used: “FBD” denotes the fossilized birth-death model, and “BDSS” the birth-death sequential sampling model. “N/A” denotes not applicable.

| Analysis | Model | Clade constraints |
| --- | --- | --- |
| 1 | FBD-1 | N/A |
| 2 | FBD-2 | (*S. fatalis*, *S. populator*), (*H. hyena*, *H. wongii*), Carnivora, Erinaceidae |
| 3 | FBD-3 | Feliformia, Caniformia, Carnivora, Erinaceidae |
| 4 | BDSS-1 | N/A |
| 5 | BDSS-2 | (*S. fatalis*, *S. populator*), (*H. hyena*, *H. wongii*), Carnivora, Erinaceidae |
| 6 | BDSS-3 | Feliformia, Caniformia, Carnivora, Erinaceidae |

Models HKY + Γ (Hasegawa et al., 1985; Yang, 1994) and Lewis’ Mkv + Γ (Lewis, 2001) were used for the molecular and discrete morphological characters, respectively. In the molecular substitution model, equilibrium frequencies were set to their empirical values, while in the discrete-morphological model, state frequencies were assumed to be equal (Lewis, 2001). Discrete Γ models were used to account for among-character rate variation; five categories were used for the molecular data, and four were used for the morphological data. Each discrete morphological data partition (see section above) was characterized by its own set of parameters.

Continuous characters were modeled using the phylogenetic BM model described above. For reasons related to analysis running times and parameter identifiability, (i) all continuous characters were assumed to share the same evolutionary rate, and (ii) among-character correlation parameters were fixed to their shrinkage estimates, and assumed constant along the phylogeny. Landmark root values (***y***_**0**_) were estimated with maximum-likelihood 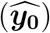 by taking the byproduct of pruning as done in Á lvarez–Carretero et al., 2019. We did so first because this approach allows for direct comparison with results from that study, and second because it is not immediately obvious how to choose a prior for ***y***_**0**_ (but see discussion). Finally, our model included three uncorrelated relaxed clock models, one for each data type, with log-normally distributed rates (Drummond et al., 2006) across branches in the phylogeny.

#### Phylogenetic inference and model comparison with Bayesian Markov Chain Monte Carlo

Phylogenetic inference and model comparison was carried out with the BEAST 2 platform, which uses MCMC to approximate the posterior distribution over model parameters. For each of the six model configurations listed in table 1, we ran three independent 1 billion-state MCMC chains starting from parameter values randomly from the prior distributions (Supplementary Table S13). Chains were sampled every 2 thousand generations, and the first 10% of samples were discarded as burn-in. All independent chains attained effective sample sizes ⩾ 200, which was our criterion for chain convergence. We then combined the chains (from each model configuration) before further analysis.

We further compared the fit of the six model configurations to the Carnivora empirical data. Models were compared in a pairwise fashion by calculating their posterior odds using the nested sampling technique (Russel et al., 2019). Details about the nested sampling analyses can be found in the supplementary material.

## 3. Results

### 3.1. Simulation study I

#### Method validation

Appropriate coverage was observed for all investigated character-evolution and phylogenetic parameters, as indicated by approximately 95 out of 100 vertical bars (representing 95%-HPD intervals) containing the true simulated parameter values (Fig. 1; Supplementary Figs. 6, 7 and 8 show the results for other parameters). These results indicate our implementation is correct.

**Fig. 1.**
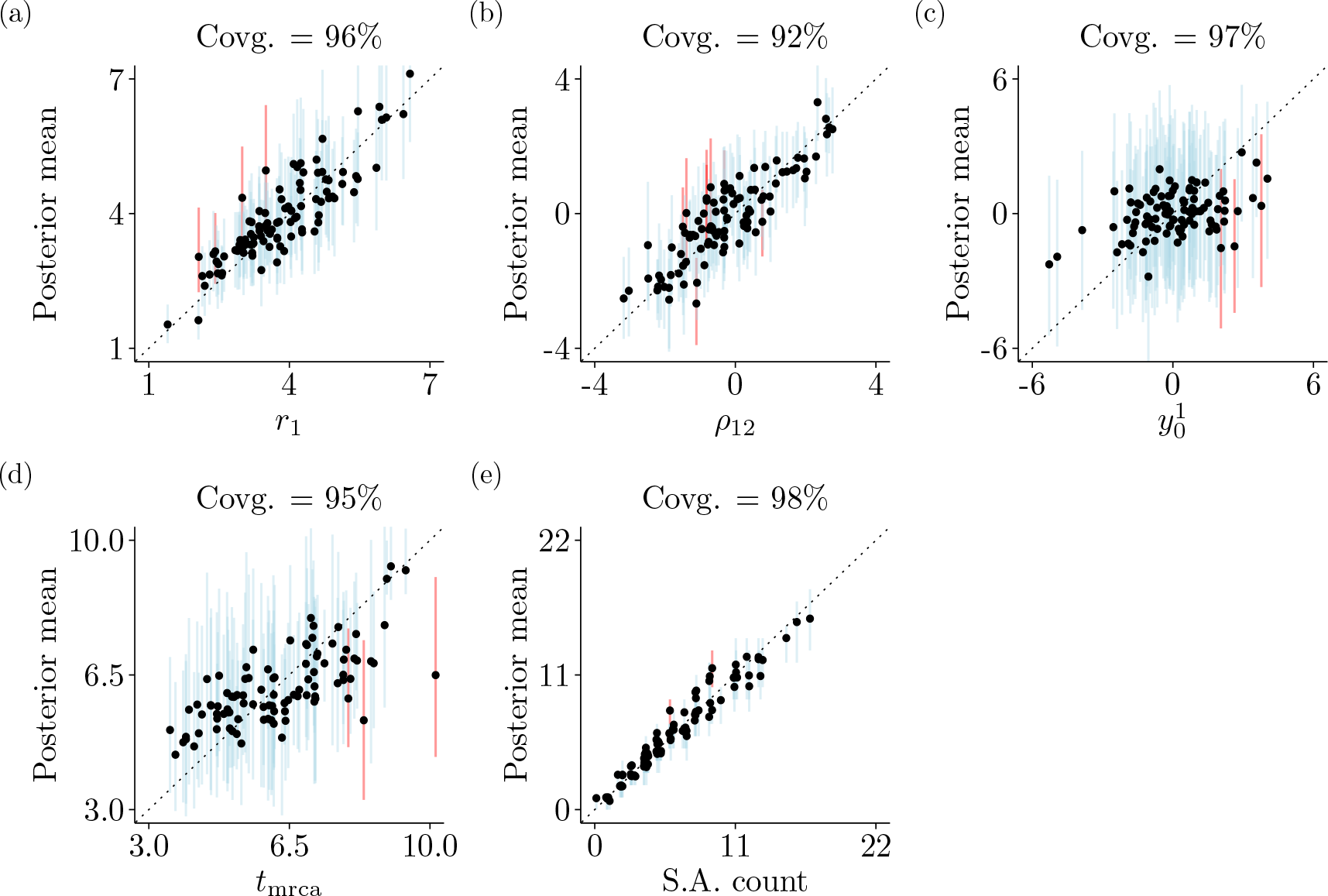
Results from simulation study I. Each panel plots mean posterior values (resulting from summarizing converged MCMC runs) for a different parameter, replicated 100 times, as a function of that parameter’s true (simulated) value. Every dot corresponds to one of the 100 replicates, with vertical bars representing the 95%-HPD intervals about the posterior means. Blue and red bars indicate 95%-HPD intervals that do and do not contain the true value, respectively. The focal parameters are (a) the evolutionary rate of character 1 (*r*_1_), (b) the covariance between characters 1 and 2 (*ρ*_12_), (c) the ancestral value of character 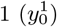, (d) the root age (*t*_mrca_), and (e) the number of fossils estimated as direct ancestors of extinct and living lineages (“S.A. count”). The identity line is shown as a dashed line in all panels.

#### Co-estimation of fossil placement and divergence times

Under correctly specified models, the age of the phylogeny and the number of direct fossil ancestors were accurately estimated from four correlated continuous characters (Fig. 1d,e). These results were observed while co-estimating the morphological evolutionary rate, the tree topology and the branch lengths. Character-related parameters were also all accurately inferred (Fig. 1a,b and Supplementary Fig. 8), with the exception of the ancestral character state at the root, for which we could mostly only recover values around the prior mean of zero (Fig. 1c; Supplementary Table S2) – accuracy for this parameter was then unsurprisingly lower.

### 3.2. Simulation study II

#### Number of characters and TED performance

The number of continuous characters increased the accuracy and precision of divergence time estimates, but the effect tapered off in the examined range (gray curve; Fig. 2a-c). Under a misspecified model (black curve; Fig. 2a-c), however, the observed trend for accuracy was reversed: increasing the number of characters beyond approximately five lowered TED accuracy.

**Fig. 2.**
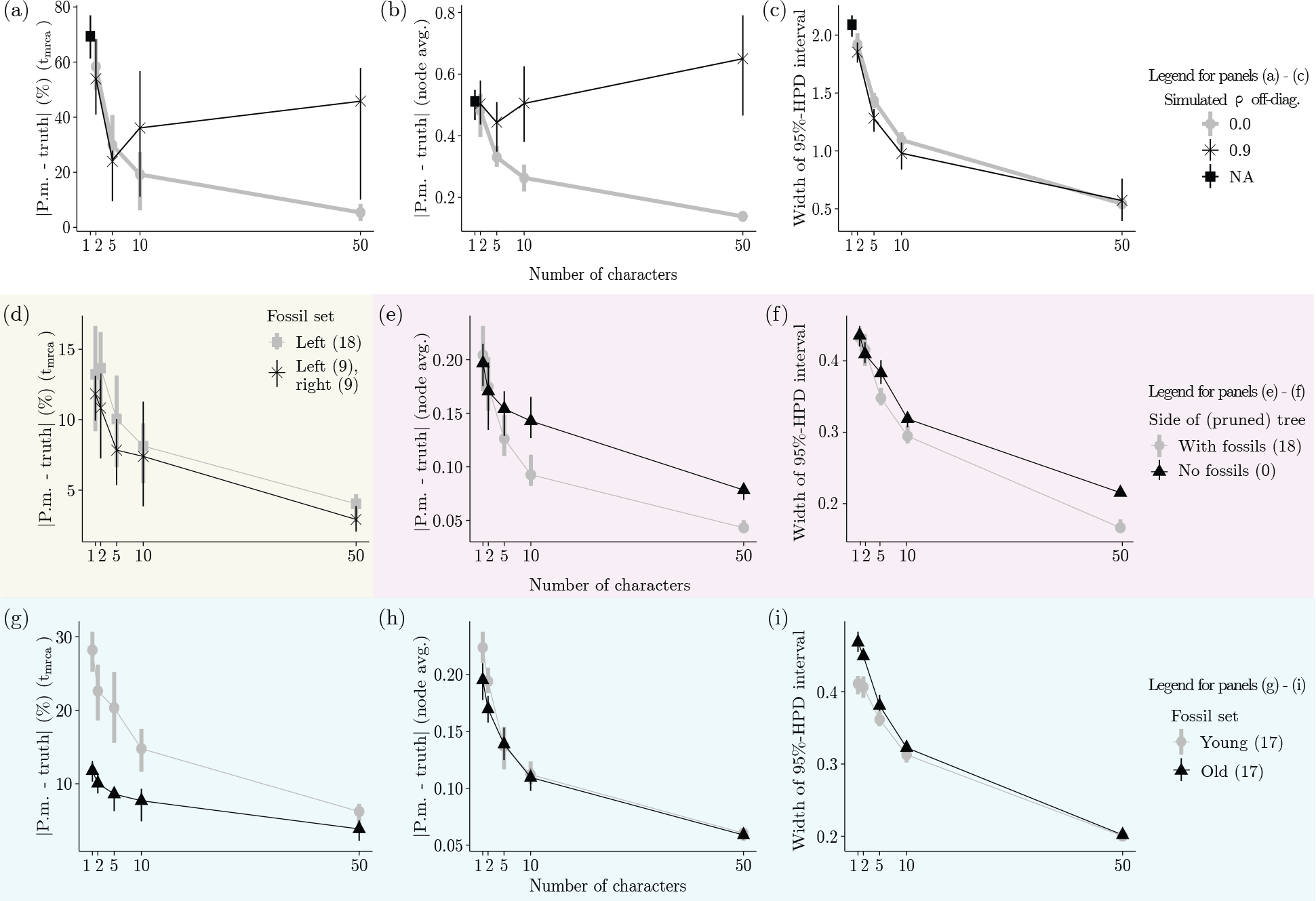
Results from simulation study II. All panels display the effect of the number of continuous characters (x-axis) on accuracy (panels a, b, d, e, g, h) and precision (panels c, f, i) of node age estimates. For both black and gray curves, each x-axis bin comprises 20 simulated replicates (i.e., a total of 40 replicates) over which the parameter tracked by the y-axis is being averaged. The whole figure thus comprises 580 replicated simulations (note that only 20 replicates were generated for 1 character in panels (a-c), under “NA”). Accuracy (y-axis of corresponding panels) is measured as the absolute difference between a parameter’s true (simulated) value and its estimated posterior mean (we show this difference as a percentage of the true value for the root age; panels a, d and g). Precision (y-axis of corresponding panels) is measured as the width of 95%-HPD intervals. Each row of panels contrasts accuracy and precision of parameter estimates with respect to a different aspect of the data, shown as curves of different colors (see main text for details). Panels (a), (d), and (g) focus on the root age (*t*_mrca_). Panels (b), (c), (e), (f), (h) and (i) focus on average internal node ages (except for the root).

#### Fossil phylogenetic relatedness and TED performance

First, we found that given an internal node of interest (e.g., the root) and a fixed number of fossils, it is mildly beneficial to have fossils on both sides of the node when estimating its age, as opposed to having all fossils on one side (Fig. 2d). Second, nodes with neighboring fossils on the same side of the tree tend to have their divergence times more accurately estimated (Fig. 2e), especially with five or more characters. Fossil proximity also improves statistical precision, but to a lesser degree (Fig. 2f). Third, older fossils contain more useful information for TED than younger fossils, as indicated by both more accurate and precise average node ages (Fig. 2h); this was found to be true for the root, the oldest node in the tree (Fig. 2g). But as with fossil phylogenetic location, this advantage decreases as more continuous characters are scored (Fig. 2g).

### 3.3. Simulation study III

#### Number of fossils and TED performance

Introducing fossils in a data set increased the accuracy and precision of node age estimation (Fig. 3a-c). However, our results suggest that it is not the case that estimates from a large number of fossils are necessarily more accurate than those from fewer fossils. When looking at the root age, TED performed worse on average for a simulated tree with 30 fossils relative to the same tree with 10 fossils (Fig. 3b). But while the effect of fossil count on statistical accuracy tapered off (from about 10-20 fossils across the parameter space we examined), precision still monotonically increased with more fossils, as reflected by increasingly narrower 95%-HPD intervals (Fig. 3c).

**Fig. 3.**
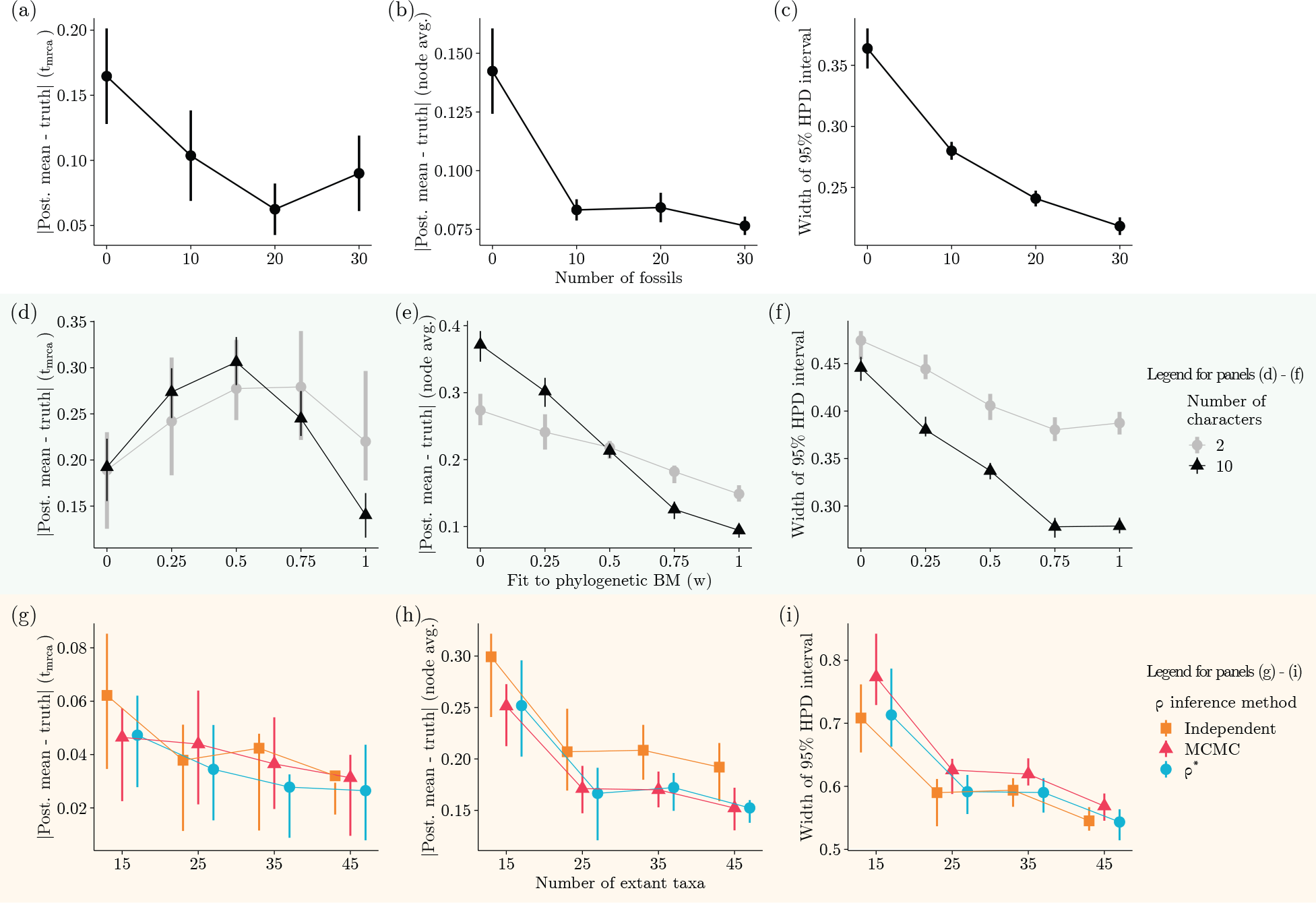
Results from simulation study III. All panels display the effect of different data features (x-axis) on accuracy (panels a, b, d, e, g, h) and precision (panels c, f, i) of node age estimates. Each x-axis bin in panels (d-f) comprises 20 simulated replicates, with bins in the remaining panels comprising 25 replicates. Parameter tracked by the different y-axes are averaged over these replicates. Ten characters were simulated for panels (a-c) and (g-i). Accuracy (y-axis of corresponding panels) is measured as the absolute difference between a parameter’s true (simulated) value and its estimated posterior mean (we show this difference as a percentage of the true value for the root age; panels a, d and g). Precision (y-axis of corresponding panels) is measured as the width of 95%-HPD intervals. Panels (d-i) further contrast accuracy and precision of parameter estimates with respect to a different aspect of the data or inference method, shown as curves of different colors (see main text for details). Panels (a), (d), and (g) focus on the root age (*t*_mrca_). Panels (b), (c), (e), (f), (h) and (i) focus on average internal node ages (except for the root).

#### Model misspecification and TED performance

When the character evolution model was perfectly specified, root age estimates were the most accurate and very precise (Fig. 3d,f). The same was true for node ages averaged over the phylogeny, although low to moderate levels of model misspecification (*w* = 0.5 and *w* = 0.75) also yielded accurate estimates (Fig. 3e). While TED under a completely misspecified model (*w* = 0.0) was the least accurate and precise when averaging over internal nodes (Fig. 3e,f), this was not true for the root node – the accuracy of root age estimates was maximally decreased by intermediate misspecification levels. The qualitative trends described above were observed for data sets with both two and ten characters; but the degree to which TED statistical accuracy and precision varied with respect to model misspecification was higher with more characters.

#### TED performance when accounting for character correlation

The three methods we examined for addressing correlation among ten characters overlapped in terms of their TED performance, as a function of tree size (Fig. 3g-i). Overall, estimated internal node ages averaged over larger phylogenies tended to be more accurate and precise (Fig. 3h,i).

### 3.4. Simulation study IV

#### Method benchmarking

We verified that our algorithms are competitively fast, improving on a previous similar method (see “Simulation study IV” section in the supplementary material). More specifically, our method always outperformed an alternative tool (Álvarez–Carretero et al., 2019) in terms of running times, with the advantage becoming greater as both the number of taxa and characters increased (top panels in Supplementary Figs. 10 and 11). In one particular setting, our method was approximately six times faster (see 50 taxa in Supplementary Fig. 10). Our tool was also superior with respect to the amount of observed autocorrelation among MCMC samples, as measured by the higher effective sample sizes (ESS) of its output relative to the alternative tool (bottom panels in Supplementary Figs. 10 and 11). Given the same MCMC chain lengths and the running-time results mentioned above, higher ESSs indicate a closer approximation of the target posterior distribution per unit time, though we must note that a better performance in this case is likely due to multiple factors beyond the model’s likelihood, such as the components of the MCMC machinery.

### 3.5. Carnivora phylogeny

All pairs of model configurations with which we analysed Carnivora (Table 1) data were substantially different in terms of model fit. The BDSS (Stadler and Yang, 2013) and FBD (Gavryushkina et al., 2014a) tree priors, both in the absence of monophyletic constraints (BDSS-1 and FBD-1; Table 2), exhibited the best and second best fits to the data, respectively. Outside of these two model settings, constraining larger clades led to a better model fit for the FBD model (FBD-3 fit the data better than FBD-2). BDSS and FBD tree priors generally led to very similar divergence time reconstructions with small topological differences, given the same monophyletic constraints (Supplementary Figs. 19b-e). Below we summarize results from analyses using the model with the highest support (BDSS-1; Fig. 4).

**Table 2.** Each cell shows a log-Bayes factor (the difference between two marginal likelihoods, *Ƶ* _0_ − *Ƶ* _1_; see supplementary material for more details) for any two models *ℳ* _0_ and *ℳ* _1_. Positive values indicate *ℳ* _0_ fits the data better than *ℳ*_1_, and negative values indicate the opposite.

| $\mathcal{M}_0 \backslash \mathcal{M}_1$ | FBD-1 | FBD-2 | FBD-3 | BDSS-1 | BDSS-2 | BDSS-3 |
| --- | --- | --- | --- | --- | --- | --- |
| FBD-1 | - | 178.48 | 169.66 | -41.31 | 150.34 | 172.57 |
| FBD-2 |  | - | -8.82 | -219.79 | -28.14 | -5.91 |
| FBD-3 |  |  | - | -210.97 | -19.32 | 2.91 |
| BDSS-1 |  |  |  | - | 191.65 | 213.88 |
| BDSS-2 |  |  |  |  | - | 22.23 |
| BDSS-3 |  |  |  |  |  | - |

**Fig. 4.**
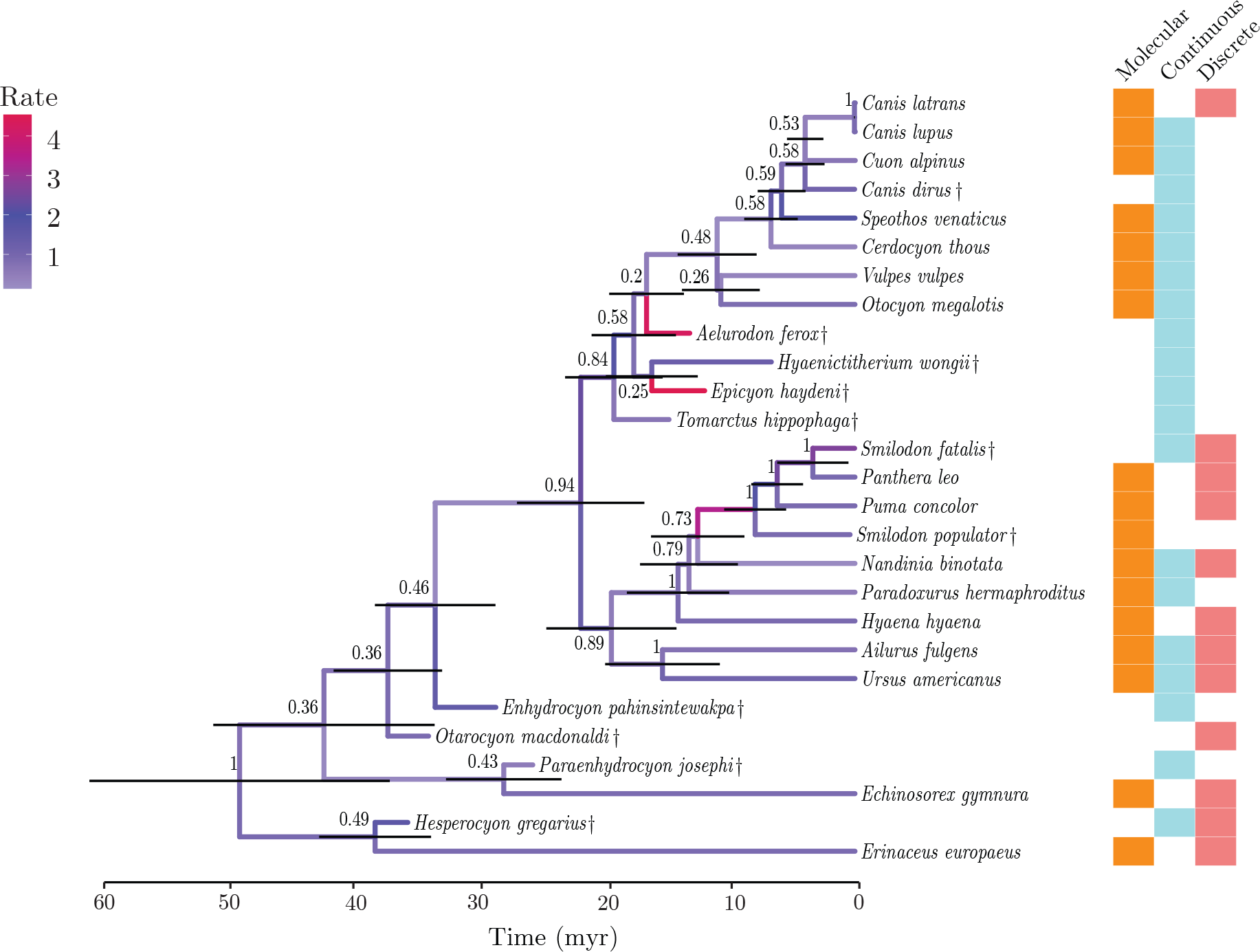
Maximum-clade-credibility summary tree of Carnivora and two outgroup taxa (extinct taxa are indicated with †), using the model configuration with the best fit to the data (“BDSS-1”, *Ƶ* = −84736.92; Table 2). Horizontal bars at internal nodes represent the 95% highest-posterior-density intervals about node ages. Numbers next to internal nodes indicate each clade’s posterior probability. Branches are colored based on the posterior mean of their (continuous) morphological evolutionary rates. The different types of data each species was scored for are summarized by colored boxes on the right. Blank squares indicate a data type that was not available for a given species.

Most caniforms grouped together in our maximum-clade-credibility (MCC) tree, with moderately high node support (posterior probability, PP, of 0.84; Fig. 4). The exceptions were the two Arctoidea species, *A. fulgens* and *U. americanus*, and the three oldest caniform fossils – *H. gregarius, P. josephi* and *E. pahinsintewakpa*. These fossils were estimated to be near the root, albeit with uncertainty. We observed a pattern of high clade support (*>* 0.5 PP) when most taxa were scored for molecular characters, and low support (*<* 0.5 PP) when taxa on one side of a clade’s most recent common ancestor (MRCA) were only scored for continuous characters. For example, internal nodes within Caniformia deeper than the MRCA of *O. megalotis* and *C. latrans* showed low support; all had one or more fossils (no molecular data) as the only descendant(s) on one of their sides. Despite uncertain evidence, however, species of Borophaginae (a subfamily of Canidae) scored for continuous characters only were inferred to be closely related, outside of the Caninae subfamily. The extinct dire wolf *Canis dirus* was also recovered as a member of the *Canis* genus based solely on continuous characters.

Species in Feliformia largely grouped together, except for the fossil hyaenid *H. wongii*, which was inferred as more closely related to dogs. The only other fossil cat, *S. fatalis* (the *S. populator* specimen is a subfossil), was emphatically inferred to be sister to modern lions. Internal nodes within Feliformia all had very high support, and so did this clade’s sister taxa, the two representatives of Arctoidea.

Imposing monophyletic constraint priors during inference had a series of consequences both in terms of topology and divergence times. When constraining Carnivora, the *Smilodon* genus, hyaenids and the outgroup (BDSS-2 and FBD-2; Table 1; Supplementary Fig. 19c,d), *H. wongii* was inferred to be outside of Caniformia, but so were *T. hippophaga* and *E. haydeni* (BDSS-2), both fossil canids of the Borophaginae subfamily (Wang et al., 1999). These two fossils were estimated to be relatively closely related under the best model configuration (Fig. 4). These constraints also reduced the support for certain nodes within Feliformia, and slightly increased certain estimated node ages. For example, the posterior mean age of the MRCA of *N. binotata* and the Felidae species (*P. leo, P. concolor*, and the *Smilodon* specimens) went from 12.51 (posterior median of 12.33) (BDSS-1; Fig. 4) to 23.77 (posterior median of 23.59) (BDSS-2; Supplementary Fig. 19c).

Constraining the monophyly of larger groups (BDSS-3 and FBD-3; Table 1) markedly increased divergence-time estimates relative to those obtained under the best model configuration. The MRCA of *N. binotata* and Felidae was inferred to be 26.91 (posterior median of 26.74) million years old and the root to be 63.91 (posterior median of 64.64) million years old (posterior mean values, in BDSS-3; Supplementary Figs. 19e), as compared to 12.51 (posterior median of 12.33) and 48.93 (posterior median of 48.03) million years old (BDSS-1), respectively, when the species tree topology was unconstrained (Fig. 4). As with the unconstrained topology analyses, the older caniform (*H. gregarius, O. macdonaldi, E. pahinsintewakpa*, and *P. josephi*) and feliform (*H. wongii*) fossils generally clustered deeper in the tree, closer to the MRCA of these clades. Of these older fossils, *H. gregarius* was further inferred to have a very high morphological rate when forced to group within Caniformia. One striking difference, however, was the topological placement of *A. ferox* well within Caninae, as opposed to outside of this clade and as sister to either or both of the Borophaginae species (*T. hippophaga* and *E. haydeni* ; e.g., Fig. 4 and Supplementary Fig. 19d,e).

### 3.6. Further comparison of cranium landmarks

Extant caniforms and feliforms formed two separate clusters on the 2-D spatial map generated by MDS of cranium landmark Euclidean distances. MDS captured a greater cranium-landmark similarity between extant ursids (*A. fulgens* and *U. americanus*) and feliforms than between ursids and caniforms. Of the fossil specimens, three had outlier cranium landmarks as compared to extant species, as revealed by MDS: *E. pahinsintewakpa, E. haydeni* and *S. fatalis* (a saber-toothed tiger specimen), the latter being the most extreme of them. Fossil hyaenid *H. wongii*, a feliform, was found to have cranium landmarks resembling those of caniform species substantially more than those of other feliforms (*P. hermaphroditus* and *N. binotata*).

## 4. Discussion

While the principle behind the “total evidence” approach – simultaneously leveraging multiple sources of data in phylogenetic reconstruction – is over 30 years old (Kluge, 1989), it was only given a statistically principled treatment in the last decade (Nylander et al., 2004; Pyron, 2011; Ronquist et al., 2012). Even the more recent examples of total evidence analyses, however, do not always model trait evolution statistically (e.g., Arrigo et al., 2013; Polotow et al., 2015), or limit themselves to discrete or discretized morphological characters (Lee and Palci, 2015). This observation is not meant as a criticism of studies modeling morphology of the discrete kind exclusively. As illustrated by the present study, discrete characters can certainly be useful. But perhaps more importantly, such data is often the only available kind in many cases, as a result of (and later maybe also as a motivator for) the historical shortage of methods for phylogenetic inference from continuous characters.

It seems unlikely that the use asymmetry between discrete and continuous characters has been driven by a perceived superiority of the former over the latter type of character, in terms of their usefulness to phylogenetic inference. In fact, discrete character models are known to face challenges with accounting for among-character correlation, and with the stationary distribution over character states (Lewis, 2001; Klopfstein et al., 2015). Discrete character data sets also suffer from subjectivity in the inclusion and scoring of characters, and from loss of information caused by the discretization of continuous characters (Goloboff et al., 2006).

The common use of maximum-parsimony in total-evidence inference is also not due to a consensus on the superiority of this criterion; rather, tools for the joint modeling of continuous morphological and molecular evolution are still lacking (but see Lartillot and Delsuc, 2012; Álvarez–Carretero et al., 2019; May and Moore, 2020; Gaboriau et al., 2020).

It was not until recently that careful simulation studies investigated the use of phylogenetic BM (Parins-Fukuchi, 2018b; Varón-González et al., 2020) and implemented statistical tools with the purpose of placing fossils and inferring phylogenies (Parins-Fukuchi, 2018a; Álvarez– Carretero et al., 2019; May and Moore, 2020).

In order to fill the methodological gap described above, we introduced a new probabilistic total-evidence method for the analysis of multiple continuous characters in addition to other types of data, e.g., molecular sequences and discrete morphological characters. Our study is a methodological contribution, and the main goal was to introduce a tool (i) that is efficient and general (both assessed in comparison to other tools whenever those exist), (ii) that implements models correctly, and (iii) that can be used with real data and produce interpretable results. With respect to (i) and (ii), our method implemented in BEAST 2 is correct and works as intended, making TED possible from continuous characters scored in living and extinct taxa. Our implementation is also more efficient than existing software (see section 6 in the supplementary material), and fast – all else being equal, it scales linearly with the number of species.

Our method’s efficiency does not come at the cost of generality. First, our tool supports both the inference of between-character correlations with MCMC, as well as their approximation by means of the linear shrinkage method. Second, because it follows a general mathematical framework (Mitov and Stadler, 2019), our method can be readily extended to include models such as BM with trends (Hansen and Martins, 1996), Ornstein-Uhlenbeck (OU; Hansen and Martins, 1996; O’Meara et al., 2006), early burst (EB; Harmon et al., 2010), accelerated or decelerated-rate models (ACDC; Blomberg et al., 2003), to name a few. In the face of all these potential extensions, we must clarify that it was not an objective of our study to investigate the many ways in which models might be misspecified (or better specified) and the downstream consequences of that. While certainly methodologically relevant, this is a task that deserves special attention in future contributions; this kind of work must also be guided by future empirical studies.

We have nonetheless gone beyond implementing, validating and benchmarking our new tool. By means of extensive exploratory simulations, we characterized the behavior of our method as a function of different data features, model assumptions, and model fit to data. To our knowledge, this is the first attempt at quantifying and synthesizing the effect the above can have on probabilistic TED; more specifically, in a situation where extant and fossil taxa are scored for multiple continuous characters, with fossil placement being estimated (rather than predetermined ad hoc). We highlight the many patterns that emerged below. These will help guide users when specifying models and choosing the most appropriate data sets, although we note that any conclusions we drew here are contingent upon the simulation settings we investigated.

### 4.7. Lessons on model behavior during TED from simulation experiments

As a rule of thumb for choosing and preparing data sets, our method will produce more accurate and precise divergence times estimates in data sets with more fossils and more scored continuous characters (Figs. 1 and 2). This is unsurprising, though the effect of fossil count tapers off quickly for node age estimate accuracy (i.e., the relationship between accuracy and number of fossils is not linear). If for some reason users must choose among a number of fossils, older fossils should be preferred over younger ones, particularly if the root node holds special meaning. This is in line with previous work investigating the impact of fossil calibration choice on divergence time estimates (Duchêne et al., 2014). Leveraging (more) young-fossil continuous characters still improves divergence time estimation (as expected from a correctly specified model), but not at the same rate as if using older fossils. Moreover, given a focal node, it seems generally beneficial to have fossils on both its descendant sides, and ideally on the same side of the tree relative to the root (both accuracy and precision are improved by the latter).

In our search for trends when model assumptions held partially, we observed that when model misspecification was not severe (when *w >* 0.5 in our experiment), scoring more characters led, again (see above), to more accurate node age estimates (Fig. 3d,e). On the other hand, when model misspecification was moderate to maximal (when *w <* 0.5), scoring a larger number of characters instead decreased the accuracy of node age estimates (Fig. 3d,e). Furthermore, our results suggest that having more characters increases estimate precision (Fig. 3f). Together, our observations should be understood by empiricists as a message of caution: if scoring a larger number of continuous characters may improve inference under a correctly specified model, it may just as well boost one’s confidence on an incorrect answer should the model be misspecified.

Another important decision to be made when using our method is what to do with among-character covariance parameters. One option is to assume characters evolve independently, for the sake of analysis running times. Doing so can hinder the estimation of divergence times if characters are all highly correlated, a problem that gets worse with more characters (Fig. 2a-c). Future analyses may suggest ways to refine this assumption, by examining its consequences when correlation values are highly variable, or variable in specific ways, e.g., if independent evolution is mostly observed among modules of characters otherwise tightly correlated. What we can say for now is that assuming among-character correlations are zero – when they in fact follow a standard normal distribution, for example – can lead to divergence time estimates that are worse than (yet still overlapping with) those obtained when inferring those parameters (see below).

When estimating covariance parameters, we showed that successful estimation of among-character covariances may depend on (i) the ratio of number of taxa to number of scored continuous characters, and (ii) what one assumes about character correlation before carrying out the analysis (Fig. 3g-i). As for the first point, phylogenetic estimates using data sets with more taxa can at times be more accurate and precise at least up to a certain taxon count. More scored taxa means more internal-node ages to estimate, but it also means more data points per free covariance parameter – whose estimation will then improve or hinder the inference of all node ages. Characterizing this trade-off should be a goal of future research.

Our analyses further showed that the different methods for accounting for character evolutionary correlation can largely overlap in terms of how accurate and precise node age estimates were. The most conservative and time-consuming consisted of estimating all covariance parameters as free parameters with MCMC. Assuming characters evolved independently (despite their non-zero simulated covariance), on the other hand, produced less accurate estimates of divergence times. These results of course reflect our simulation conditions. Assuming that the trait covariance structure is shared by all species over time, or that traits are independent, will lead to varying levels of model misspecification given different empirical data sets.

### 4.8. Lessons on model behavior during TED from empirical analyses

In addition to the large number of simulated scenarios under which we validated and characterized our method’s behavior, we conducted a TED analysis of a Carnivora data set we compiled from two different studies (Álvarez–Carretero et al., 2019; Barrett et al., 2021). We note that we did not particularly aim at contributing to a greater understanding of Carnivora evolution, e.g., by producing the most correct Carnivora phylogeny to date. This would entail collecting larger molecular and morphological data sets, scoring more fossils, and doing extensive model comparison and sensitivity analyses. Such thorough data curation and model analysis make fundamental contributions to organismal biology. Here, the goal of method development is to test a method so as to examine its features and behavior, often under potentially unfavorable conditions.

As we illustrate here, the value of empirical analyses is not limited to what definitive answers it may bring to questions about classification and dating. Although we do not know the truth when dealing with empirical data, the same data can be analysed under distinct models and results can be then compared among themselves and with those from other studies. We demonstrate how, by doing so, one can (a) uncover general method behavior patterns (in the face of real, noisy data) that users might expect, guiding future tool use, (b) determine where the method development effort should focus next by gauging the magnitude of methodological shortcomings, and (c) shed a different light on conclusions drawn from other empirical studies, provided similar data sets have been employed.

By comparing different models when analyzing Carnivora data, we observed that model fit changed significantly when monophyletic constraints were added to a model, with the best fit being observed when no constraints were imposed on the phylogeny, followed by models constraining clades defined by deep nodes, and then by those constraining smaller, generally younger clades. This result may be unsurprising if one expects topological constraints and the signal in the data to be in disagreement, but is an observation that has nonetheless remained relatively underappreciated as a result of inference being often done on a fixed phylogeny (e.g., Álvarez–Carretero et al., 2019). Generally speaking, the consequence of monophyletic constraints to a model’s fit to data is likely to be contingent upon the contrast between a researcher’s evolutionary hypothesis and the signal present in the data. But more analyses like ours are needed if one wishes to uncover the connections (or lack thereof) between clade characteristics, such as size or age, and their effect on model fit in the presence of monophyletic constraints.

We also observed that tree priors resulting in largely overlapping topologies and divergence times can still substantially differ in their fit to data. So it would seem that monophyletic constraints are not the only way to obtain a sharp contrast between a model’s assumptions and the evidence in the data. In any case, it is difficult to know the extent of this contrast ahead of phylogenetic inference; this is a place where domain knowledge, familiarity with the data, and techniques like MDS (e.g., the present study) or principal component analysis (PCA; Álvarez–Carretero et al., 2019) can prove critical. For example, our MDS analysis of cranium landmarks revealed that both feliform fossils involved in smaller clade constraints, *H. wongii* and *S. fatalis*, have extreme morphology in relation to the other two extant feliform species (*N. binotata* and *P. hermaphroditus*). In particular, *H. wongii* groups much more closely with caniforms in the two-dimensional space produced by MDS, which suggests a strong signal in the data against the grouping of this fossil within Feliformia.

Other than the obvious topological outcomes expected under monophyletic priors, we observed a few different trends emerging from constraining species relationships. For example, when constraining hyaenids and genus *Smilodon* under the BDSS model, internal nodes neighboring and upstream of these clades were estimated to be older (Supplementary Fig. 19c) than their counterparts in the unconstrained model (Fig. 4). Among the affected nodes, close ancestors of the constrained lineages were also characterized by lower node support. We hypothesize that these results are being driven by a difficulty in accommodating the divergent morphology of *H. wongii* and *S. populator* on the postulated topology, as a result of (i) the greater similarity of these fossils to more distantly related specimens, (ii) the continuous-morphology clock model, and (iii) the challenge of learning how correlated character evolution is. As far as some of our results suggest, managing those three items is harder the larger the number of taxa being forcefully placed in the phylogeny despite their morphology.

As for (i), when lineages with extreme character values are forced to group within a clade with closer yet more morphologically dissimilar relatives, these morphological outlier taxa may attract lineages that are foreign to the constrained clade. This seems to have been the case for fossils *T. hippophaga* and *E. haydeni* in our analysis of Carnivora, who got pulled by *H. wongii* into Feliformia despite being canids. Unless otherwise constrained, during Bayesian inference these “hitchhiker” taxa will be moved around the focal constrained lineage(s), causing neighboring and upstream node support to decline. In the case of (ii), if morphological clock rates are constrained by their prior distributions, homoplasy-like patterns in the data are expected to lead to inflated divergence times (Ogilvie et al., 2022) – though under sufficiently diffuse rate priors (or, conversely, strongly informed tree priors), morphological rates may instead be inflated (Mendes et al., 2018).

The rationale behind (iii) involves realizing that highly correlated character evolution may allow distantly related taxa to evolve similar complex morphologies that would be otherwise unlikely. Failing to account for character correlation could thus put an erroneously low ceiling on the likelihood of true homoplasy, causing such trait patterns to be accommodated by (artefactually) older node ages (what Ronquist et al., 2016, dubbed “deep root attraction”, when carrying out TED with discrete morphological characters). If our modeling of character correlation is inadequate (see more below), it is a possibility that our monophyletic priors are having an effect analogous to the effect that can be induced by vague tree priors (e.g., those employed in Ronquist et al., 2016), leading to a phenomenon akin to deep root attraction. Moreover, deep root attraction seems to manifest in complex conditions that include not just the misspecification of character evolution models, but also the violation of taxon sampling assumptions – an issue that can be curbed by the use of diversified sampling approaches and tree priors that penalize ghost lineages (Höhna, 2014; Ronquist et al., 2016). This is a topic worthy of investigation that deserves careful treatment and that we leave for future research.

Constraining larger clades (Supplementary Fig. 19d,e) – Erinaceidea (the outgroup), Carnivora (the ingroup), Caniformia and Feliformia – led to divergence time estimates in large agreement with previous work (Álvarez–Carretero et al., 2019; Barrett et al., 2021).

Mean divergence times in Caniformia and Feliformia were generally older in these analyses when compared to results from our best-fit (unconstrained) model. The only exceptions were the internal nodes that were direct ancestors or MRCAs of the older caniform fossils (*H. gregarius, P. josephi* and *E. pahinsintewakpa*), whose overlapping 95%-HPDs remained mostly unchanged. Under the unconstrained model, these fossils were inferred to be close descendants of the root and other deep nodes, whose node-age 95%-HPDs ranged between approximately 30 to 50 million years old (Fig. 4). When the Caniformia monophyletic constraint was applied (as in Álvarez–Carretero et al., 2019), these fossils were placed as close descendants of the MRCA of this group instead, with HPDs ranging again between approximately 30 and 50 million years old (Supplementary Fig. 19d,e); under this latter model, other internal nodes within Caniformia were pulled towards the past as a side-effect.

When carrying out TED, constraining living and extinct taxa within a clade can thus induce a lower bound on the clade’s age. This outcome might be intended by the empiricist, and it echoes the use of fossils for node calibration (Ho and Phillips, 2009), which involve specifying explicit bounds for clade ages. In comparison to the results from our best-fit model, it is unclear what aspect of this and other observed effects during TED (e.g., the inflation of node ages within a constrained clade) reflects the truth, and what should be treated as bias. Methods capable of co-estimating tree topologies and divergence times from continuous characters, such as the one we introduce here, will prove critical in future simulation work investigating the interaction between different modeling choices and fossil data. Comparative studies examining model (mis)specification outcomes across a range of clades (e.g., Drummond et al., 2006; Mendes et al., 2019) will further illuminate the degree to which our observations are to be expected with other data sets.

We are only beginning to understand the power and utility of leveraging discrete and continuous morphology in addition to molecules within a robust statistical framework. Even if many clades are not prone to fossilization, the vast majority of species to ever roam the planet have gone extinct (Lee and Palci, 2015), and obtaining their DNA (but not measuring their morphology) is challenging at best (Cooper and Poinar, 2000; Hagelberg et al., 2015), and impossible in most cases (Austin et al., 1997). Here, we observed continuous characters having strong influence over the likelihood of certain phylogenetic reconstructions, while also being uninformative with respect to the resolution of certain nodes.

For example, the sabertooth cat, *S. fatalis*, was placed among dogs (Supplementary Fig. 20) with strong support when only its outlier continuous morphology was used (Fig. 5). This suggests the phylogenetic BM or morphological clock models used here (and elsewhere; Álvarez–Carretero et al., 2019) were misspecified in some manner. Assuming that all continuous characters evolve at the same rate along a branch, like we did, is not a requirement of our implementation. Although it is unlikely to hold in most empirical contexts, making this assumption improves analysis running times. It remains to be determined how serious this type of model violation can be, and how often. But if history has taught us anything, such issues will be alleviated by the development and use of methods for accommodating among-character rate variation (e.g., Yang, 1996; Sullivan and Swofford, 1997). Our implementation should be immediately compatible with such methods, as it is already possible to relax the assumption of equal among-character rates.

**Fig. 5.**
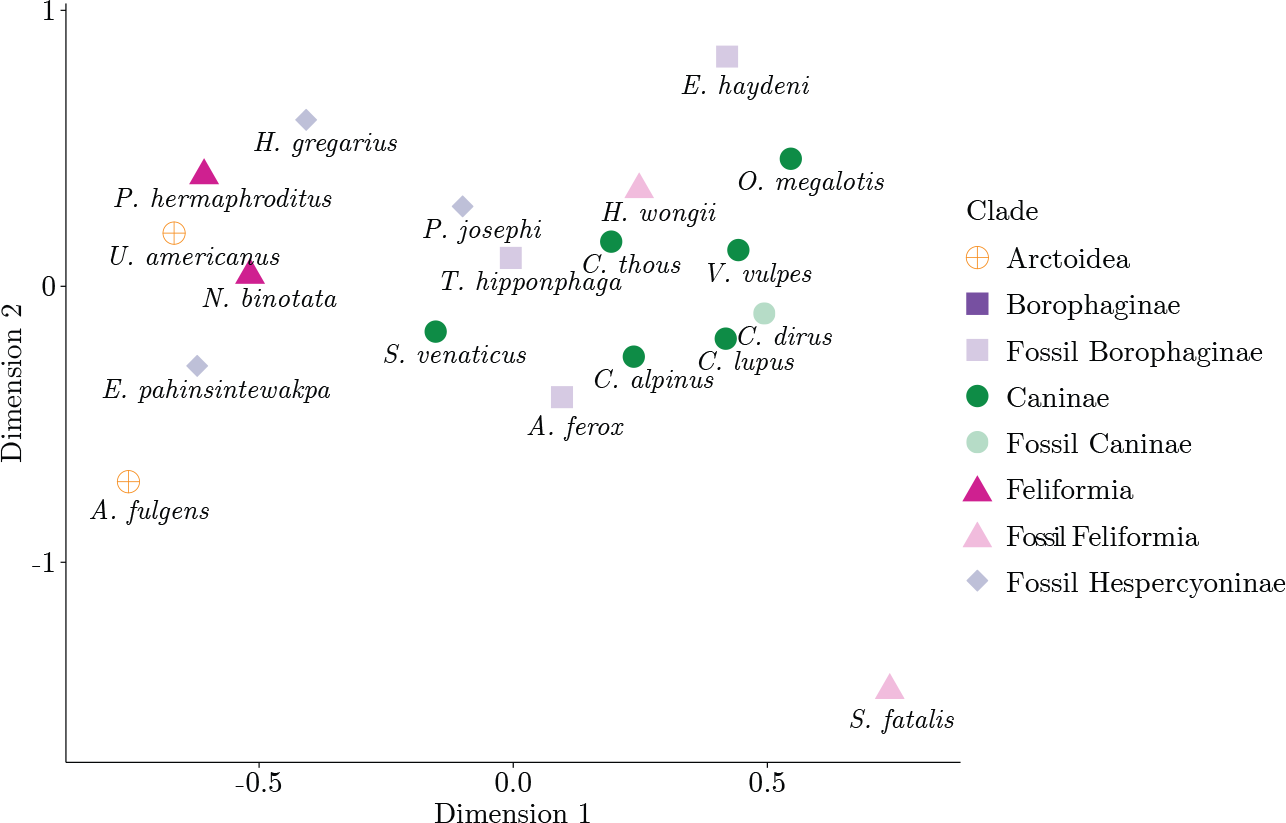
Multidimensional scaling (MDS) of Euclidean distances between Carnivora species cranium landmarks. Shapes with increased transparency represent fossil taxa, while the remaining shapes represent extant taxa.

Despite its outlier continuous morphology, the sabertooth cat still grouped within Felidae (Fig. 4) when all data are used, due to this specimen’s discrete morphology. The extreme continuous character values of *S. fatalis* were as a result accommodated through elevated evolutionary rates along this lineage’s history. This outcome captures the main idea behind TED: allowing different data types to collectively inform inference and to compete should their phylogenetic signals disagree for whatever reason, e.g., model misspecification underlying one or more of the data types. When species are scored for only one type of data (e.g., *C. dirus* and *H. wongii*), however, there is less of a buffer for things like model misspecification. The extinct dire wolf *C. dirus* grouped with its congeners in *Canis* in agreement with the literature (Prevosti, 2010; but see Perri et al., 2021), and is a fortunate example in which model misspecification was not excessive. Fossil *H. wongii*, on the other hand, was inferred to be in Caniformia (rather than Feliformia; Ö zkurt et al., 2015), within which its placement was uncertain. In the case of this fossil hyaenid, the scoring of discrete characters by an expert could potentially improve its inferred phylogenetic placement.

We believe there is promise in the use of continuous characters in TED, as this data type can clearly inform the phylogenetic placement of both extant and extinct taxa. Although part of our reconstructions of Carnivora are likely incorrect, such as the placement of Arctoidea species within Feliformia, we remain optimistic because we have used relatively simple evolutionary models for continuous characters and their rates of change.

### 4.9. Moving forward

Many modeling approaches remain open, some of which might help remedy the issues we and others have observed. Local morphological clock models applied to evolutionary rates and adaptive optima regimes (in the case of OU models; Bastide et al., 2020), for example, could further accommodate ecologically relevant traits evolving under selection (Eastman et al., 2011; Uyeda and Harmon, 2014; Gaboriau et al., 2020). Moreover, such characters can display evolutionary trends or evolve at high rates and as a result their ancestral states can be hard to estimate (Oakley and Cunningham, 2000; Webster and Purvis, 2002). The success of TED using continuous characters may thus depend on which strategy for inferring or integrating out ancestral states is used (e.g., conjugate priors; Pybus et al., 2012; Solís– Lemus et al., 2014). All of the topics above deserve future studies of their own.

In the present work we used one of the most efficient shrinkage estimators for the character variance-covariance matrix, the archetypal ridge estimator, which employs a linear shrinkage penalty. Other estimators have been proposed for phylogenetic inference, however, and could be investigated in the context of TED, such as the LASSO and quadratic ridge estimators (Clavel et al., 2019). Importantly, these different estimators have different computational efficiencies, and not all are rotation-invariant. In the latter case, those estimators are not robust to the reorientation of continuous characters, typical of geometric morphometric data (Clavel et al., 2019). Data set sizes and the nature of the continuous characters will thus determine the appropriateness of each of these different methodological alternatives.

Critically, strategies like leave-one-out-cross-validation (Clavel et al., 2019) for estimating the variance-covariance matrix (using the options mentioned above) are difficult to conceive and implement in a Bayesian context. As in previous work (Álvarez–Carretero et al., 2019), we instead assumed character correlation is the same within and between species, for all species. This is a strong assumption that is known to not hold for different types of continuous characters (e.g., lifespan and body size; Garamszegi, 2014). Future simulation and empirical studies may further illuminate the importance of modeling the evolution of character correlation itself. These investigations, particularly if large data sets are involved, will benefit from the use of convenient prior distributions (e.g., the Wishart distribution, Cybis et al., 2015; shrinkage priors, van Erp et al., 2019), and from both more efficient Bayesian proposals (e.g., that decouple character variance and covariance parameters; Caetano and Harmon, 2017), and sampling mechanisms (Bastide et al., 2020).

Accounting for gene tree discordance among the loci underlying continuous characters may also prove necessary (Mendes et al., 2018), especially if it is shown that population-level processes cannot be merely buffered out through an additional variance term. Progress made on this front (Bastide et al., 2018) and on the phylogenetic modeling of intraspecific trait variance (Gaboriau et al., 2020) might hold the key to capturing additional dimensions of phenotypic evolution.

We are confident that methods such as the one described here will motivate the curation, expansion and publication of rich morphological data sets, which will in turn create a demand for novel statistical models and methods. We trust that this feedback loop will fuel the integrative modeling paradigm, of which TED is an example, and push the frontiers of comparative biology and phylogenetics.

## Supplementary Material

Data available from the Dryad Digital Repository: https://doi.org/10.5061/dryad.YYYY

## Acknowledgments

We thank Julien Clavel and Venelin Mitov for useful discussions on the implementation of continuous trait evolutionary models, and Sandra Álvarez-Carretero for assistance with MCMCTree analyses. We thank Cécile Ané and Michael Landis for helpful discussions that improved this manuscript. This work was supported by the China Scholarship Council (No. 201706990021; RZ), by the Royal Society of New Zealand (16-UOA-277; FKM and AJD), and by the National Science Foundation (grant DEB-2040347; FKM).

