## Supplementary material for "Fast Bayesian inference of phylogenies from multiple continuous characters"

### 1. Phylogenetic Brownian motion model for continuous characters

#### 1.1 Calculating the probability of the data with phylogenetically independent contrasts (PIC)

As explained in the main text, calculating the phylogenetic Brownian motion (BM) model likelihood in its GLS form (sometimes referred to as the "matrix inverse" method; solving Eq. (1) in main text "as is") can be computationally onerous if the numbers of species $n$ and/or characters (traits) $k$ are large. The burden comes from the computation of $\boldsymbol{V} = \boldsymbol{\Sigma} \otimes \boldsymbol{T}$ and its inversion.

Felsenstein (1973) proposed a pruning solution that scales linearly with $n$ (though assumed independence among characters), later further formalized by Freckleton (2012). This method traverses the tree making use of character-value contrasts (between children of a node) at every internal node, from tips to root. If we assume characters can be correlated, at every internal node $g$ with children nodes $u$ and $w$, we compute its log-likelihood as:

$$\ln f_g(\boldsymbol{x}_g|\Phi, \boldsymbol{b}_m, c_m, \boldsymbol{\Sigma}, \boldsymbol{y_0}) = -\frac{k}{2}\ln(2\pi) - \frac{1}{2}\ln|(t_u + t_w)\boldsymbol{\Sigma}| - \frac{1}{2}\boldsymbol{x}_g{}^{\mathrm{T}}[(t_u + t_w)\boldsymbol{\Sigma}]^{-1}\boldsymbol{x}_g, \quad (1)$$

where $\boldsymbol{x}_g$ is the vector of contrasts between character values of $u$ and $w$, i.e., $\boldsymbol{x_g} = \boldsymbol{m_u} - \boldsymbol{m_w}$, and where $t_u$ (and $t_w$, replacing the subscript) is defined after the main text notation (Eq. (2)):

$$t_u = c_m b_m^u \ell(u). \quad (2)$$

We remind the reader that $c_m$ is global morphological clock rate, $b_m^u$ is $u$'s relative branch morphological clock rate, and $\ell(u)$ is the length of branch $u$ in absolute time units. Note that we abuse notation in Eq. (1) in that $t_u$ (and $t_w$) is a function of other parameters (Eq. (2)), and $\boldsymbol{y_0}$ is used only after pruning step at the root has been done (see below).

Recall that (i) $\mathbf{M} = [m_{ij}]$ is an $n \times k$ matrix, with $m_{ij}$ corresponding to the $j$-th character from the $i$-th species; $\boldsymbol{m_i}$ denotes the vector of all character values from species $i$, and (ii) $\boldsymbol{\Sigma}$ is a $k \times k$ matrix, given by $\boldsymbol{\Sigma} = \mathrm{diag}(\sqrt{\boldsymbol{r}}) \times \boldsymbol{\rho} \times \mathrm{diag}(\sqrt{\boldsymbol{r}})$; $\boldsymbol{r} = \{r_i : 1 \leq i \leq k\}$ is a vector containing all relative character-specific evolutionary rates, and $\boldsymbol{\rho}$ contains pairwise character correlation values in its off-diagonal elements. Note that Eq. (1) is the same as Eq. (10) in Freckleton (2012), but in a more general form.

The character-value contrasts of an internal node $g$ is in turn given by:

$$\boldsymbol{x}_g = \boldsymbol{m}'_u - \boldsymbol{m}'_w, \quad (3a)$$

or after the root node $g = (2n - 1)$ has been visited (and now $g = 2n$; see Eq. (4)):

$$\boldsymbol{x}_g = \boldsymbol{m}'_g - \boldsymbol{y_0}. \quad (3b)$$

Eq. (3b) accounts for our Bayesian estimation of the root character values, $\boldsymbol{y_0}$, which in our framework will have a prior distribution and be sampled with MCMC. When we calculate the log-likelihood of the parent of $g$, then:

$$\boldsymbol{m}'_g = \begin{cases} \boldsymbol{m}_u, & g \text{ is a tip} \\ \frac{t_w \boldsymbol{m}_u + t_u \boldsymbol{m}_w}{t_u + t_w}, & g \text{ is an internal node} \end{cases} \quad (3c)$$

$$t'_g = \begin{cases} t_g, & g \text{ is a tip} \\ t_g + \frac{t_u t_w}{t_u + t_w}, & g \text{ is an internal node} \end{cases} \quad (3d)$$

Finally, the probability density of $\mathbf{M}$ under phylogenetic BM over the entire tree $\Phi$ is then given by:

$$\begin{aligned}\ln f(\mathbf{M}|\Phi, \boldsymbol{b}_m, c_m, \boldsymbol{\Sigma}, \boldsymbol{y_0}) &= \sum_{g=n+1;u,w\in\mathrm{Desc}(g)}^{2n} \ln f_g(\boldsymbol{x}_g|t_u, t_w, \boldsymbol{\Sigma}, \boldsymbol{y_0}) \\ &= \sum_{g=n+1;u,w\in\mathrm{Desc}(g)}^{2n} -\frac{k}{2}\ln[2\pi(t'_u + t'_w)] - \frac{1}{2}\ln|\boldsymbol{\Sigma}| - \frac{1}{2(t'_u + t'_w)}\boldsymbol{x}_g^{\mathrm{T}}\boldsymbol{\Sigma}^{-1}\boldsymbol{x}_g \end{aligned} \tag{4}$$

Again, we abuse notation in the same way as Eq. (1).

##### 1.2 Restricted maximum-likelihood (REML) implementations

Phylogenetic BM model implementations can sometimes consider the root character value ($y_0$) as a nuisance parameter by running the summation in Eq. (4) over $(2n-1)$ instead of $2n$; in Felsenstein's (1973) words, this has the effect of "dropping the estimation of $[y_0]$" (our notation within squared brackets). This implementation is referred to as the "restricted maximum-likelihood" method (REML). Occasionally, one might find descriptions that equate the PIC methods to REML* , though the modification to the density function mentioned above is not mandatory in PIC methods.

As formally described for one character (Felsenstein, 1973), the PIC implementation does not *by default* consider $y_0$ as a nuisance parameter, and yields a probability density identical to our Eq. (1) in the main text (Eq. (1) and (5) in Freckleton, 2012), which is sometimes referred to as the "full likelihood". This is reflected in the summation going from 0 to $(n-1)$ in Eq. (5) in Freckleton (2012), which matches Eq. (4) here**. Freckleton (2012) further specifies that REML requires modifying his Eq. (5) to sum from 1 (rather than 0) to $(n-1)$.

Despite not being described in Álvarez–Carretero et al. (2019)'s text, their implementation (against which we benchmark and compare ours) uses restricted likelihood. This can be seen from Álvarez–Carretero et al. (2019)'s Eq. (4), where the product is from $(s+1)$ to $(2s-1)$ (rather than to $2s$, where $s$ is their number of species). We also observed that probability densities computed with `MCMCTree` and our implementation under restricted likelihood were the same.

#### 2. BEAST 2's implementation of Mitov and Stadler (2019)'s method

The method of Mitov and Stadler (2019) we implement in BEAST 2 is also a dynamic programming algorithm, but it formulates the probability density of a very general class of Gaussian continuous-character phylogenetic models, of which the phylogenetic BM is a specific case. Rather than calculating the probability density of character value sister-node contrasts, Mitov and Stadler (2019)'s builds on top of:

$$\ln f(\boldsymbol{m}_u|\boldsymbol{m}_g, \boldsymbol{\Sigma}) = -\frac{k}{2}\ln(2\pi t_u) - \frac{1}{2}\ln|\boldsymbol{\Sigma}| - \frac{1}{2t_u}(\boldsymbol{m}_u - \boldsymbol{m}_g)^{\mathrm{T}}\boldsymbol{\Sigma}^{-1}(\boldsymbol{m}_u - \boldsymbol{m}_g), \tag{5}$$

* "PICs are a formulation of a REML model", https://lukejharmon.github.io/pcm/chapter4_fitbm/#footnote4.3
**It is also reflected in method 'pic' from function `mvBM()` from the `mvMORPH` R package (Clavel et al., 2015)

which represents the probability density of the character values at node $u$, $\boldsymbol{m}_u$, given the character values at its parent node $g$, $\boldsymbol{m}_g$. The contrasts in Eq. (5) are between parent and child nodes. We note that Eq. (5) simplifies Eq. (3) in Mitov and Stadler (2019) with their $\boldsymbol{\omega}_u = \mathbf{0}$ ($k$-sized vector) and $\boldsymbol{\Phi}_u = \mathbf{I}$ ($k \times k$) (replacing the subscripts). This probability density can be presented as a quadratic polynomial (Mitov and Stadler, 2019):

$$\ln f(\boldsymbol{m}_u|\boldsymbol{m}_g, \boldsymbol{\Sigma}) = \boldsymbol{m}_u^{\mathrm{T}}\mathbf{A}_u\boldsymbol{m}_u + \boldsymbol{m}_g^{\mathrm{T}}\mathbf{C}_u\boldsymbol{m}_g + \boldsymbol{m}_g^{\mathrm{T}}\mathbf{E}_u\boldsymbol{m}_u + f_u, \tag{6}$$

where:

$$\begin{aligned}
\boldsymbol{A_u} &= -\frac{1}{2t_u}\boldsymbol{\Sigma}^{-1} \\
\boldsymbol{C_u} &= -\frac{1}{2t_u}\boldsymbol{\Sigma}^{-1} \\
\boldsymbol{E_u} &= \frac{1}{t_u}\boldsymbol{\Sigma}^{-1} \\
f_u &= -\frac{k}{2}\ln(2\pi t_u) - \frac{1}{2}\ln|\boldsymbol{\Sigma}|\,.
\end{aligned} \tag{7}$$

Again, note that we Eq. (7) is simplified from Mitov and Stadler (2019)'s Eq. (1), as their $\boldsymbol{b}_u$ and $\boldsymbol{d}_u$ (replacing the subscripts) reduce to $\mathbf{0}$ ($k$-sized vectors).

If we then write the probability density of $\mathbf{M}_i$ at any internal node $i$ that roots a (sub-)tree as:

$$\ln f(\mathbf{M}_i|\Phi, \boldsymbol{b}_m, c_m, \boldsymbol{\Sigma}, \boldsymbol{y}_i) = \underbrace{\left[\sum_{u=1}^{n} \ln f(\boldsymbol{m}_u|\boldsymbol{y}_i)\right]}_{\text{tips}} + \underbrace{\left[\sum_{u=n+1}^{2n-1} \ln[\int f(\boldsymbol{m}_u|\boldsymbol{y}_i) f(\mathbf{M}_u|\boldsymbol{m}_u)\mathrm{d}_{\boldsymbol{m}_u}]\right]}_{\text{non-tips}}, \tag{8}$$

where node $i$ is the parent of node $u$, and set:

$$\begin{aligned}
\boldsymbol{\alpha}_i{}^{\text{tips}} &= \sum_{u \in \text{Desc}(i)} \boldsymbol{C_u} \\
\boldsymbol{\beta}_i{}^{\text{tips}} &= \sum_{u \in \text{Desc}(i)} \boldsymbol{E_u}\boldsymbol{m}_u \\
\gamma_i{}^{\text{tips}} &= \sum_{u \in \text{Desc}(i)} \boldsymbol{x}_u^{\mathrm{T}}\boldsymbol{A_u}\boldsymbol{m}_u + f_u
\end{aligned} \tag{9}$$

$$
\begin{aligned}
\boldsymbol{\alpha_i}^{\text{non-tips}} &= \sum_{u \in \text{Desc}(i)} \boldsymbol{C_u} - 0.25 \times \boldsymbol{E_u}(\boldsymbol{A_u} + \boldsymbol{\alpha}_u)^{-1} \boldsymbol{E_u}^{\text{T}} \\
\boldsymbol{\beta_i}^{\text{non-tips}} &= \sum_{u \in \text{Desc}(i)} -0.5 \times \boldsymbol{E_u}(\boldsymbol{A_u} + \boldsymbol{\alpha}_u)^{-1} \boldsymbol{\beta}_u \\
\gamma_i^{\text{non-tips}} &= \sum_{u \in \text{Desc}(i)} f_u + \gamma_u + \frac{k}{2}\ln(2\pi) - 0.5\ln(|-2(\boldsymbol{A_u} + \boldsymbol{\alpha}_u)|) \\
&\qquad -0.25 \times \boldsymbol{\beta}_u^{\text{T}} (\boldsymbol{A_u} + \boldsymbol{\alpha}_u)^{-1} \boldsymbol{\beta}_u,
\end{aligned}
\tag{10}
$$

then Eq. (8) (Eq. (6) in Mitov and Stadler, 2019) can be simplified to:

$$
\begin{aligned}
\ln f(\mathbf{M}_i | \Phi, \boldsymbol{b}_m, c_m, \boldsymbol{\Sigma}, \boldsymbol{y}_i) &= \boldsymbol{y_i}^{\text{T}} \boldsymbol{\alpha}_i \boldsymbol{y}_i + \boldsymbol{y}_i^{\text{T}} \boldsymbol{\beta}_i + \gamma_i \\
&= \boldsymbol{y_i}^{\text{T}} (\boldsymbol{\alpha}_i^{\text{tips}} + \boldsymbol{\alpha}_i^{\text{non-tips}}) \boldsymbol{y_i} + \boldsymbol{y}_i^{\text{T}} (\boldsymbol{\beta}_i^{\text{tips}} + \boldsymbol{\beta}_i^{\text{non-tips}}) \\
&+ (\gamma_i^{\text{tips}} + \gamma_i^{\text{non-tips}}).
\end{aligned}
\tag{11}
$$

The probability density of the full data set becomes (Eq. (S1) from Appendix A in Mitov and Stadler, 2019):

$$
\ln f(\mathbf{M} | \Phi, \boldsymbol{b}_m, c_m, \boldsymbol{\Sigma}, \boldsymbol{y_0}) = \boldsymbol{y_0}^{\text{T}} \boldsymbol{\alpha}_0 \boldsymbol{y_0} + \boldsymbol{y_0}^{\text{T}} \boldsymbol{\beta}_0 + \gamma_0. \tag{12}
$$

#### 3. Alternative parameterization: $\boldsymbol{\Sigma} = \boldsymbol{L}\boldsymbol{L}^{\text{T}}$

No matter which algorithm one chooses to evaluate the likelihood of phylogenetic BM, being able to invert $\boldsymbol{\Sigma}$ is a requirement (see Eq. (4) and Eq. (10)). One naïve strategy to meet this requirement during Bayesian inference is to reject $\boldsymbol{\Sigma}$ when it cannot be inverted (i.e., when $\boldsymbol{\Sigma}$ has a determinant of zero). There are at least two reasons why this strategy is sub-optimal. First, especially when the number of characters is large, too many different $\boldsymbol{\Sigma}$ instances end up being rejected during MCMC, which makes the procedure inefficient. (In particularly bad cases, it is very hard to even initialize a very large $\boldsymbol{\Sigma}$ matrix before the chain is started.) Second, rejecting MCMC proposals in such way will lead to a mismatch between the prior distribution specified for the analysis and that that is "manifested" during inference. In other words, censoring certain parameter values causes model misspecification.

Another strategy is to use a procedure similar to the Cholesky decomposition of $\boldsymbol{\Sigma}$ (referred to here as "upper-triangular" decomposition; Eq. (S4) in Mitov and Stadler, 2019), $\boldsymbol{\Sigma} = \boldsymbol{L}\boldsymbol{L}^{\text{T}}$. We start by composing a $k \times k$ upper-triangular matrix $\boldsymbol{L}$ (where $k$ is the number of characters) whose diagonal elements $\boldsymbol{p} = \{p_1, p_2, ..., p_k\}$ are all non-zero, e.g., $P \sim \text{Lognormal}(\cdot)$. The $(k \times (k-1)/2)$ off-diagonal elements can be populated by first drawing values $\boldsymbol{o}$ from a uniform distribution bounded by -1 and 1, i.e., $O \sim \text{U}(-1, 1)$, and then:

$$
\boldsymbol{L} = \begin{bmatrix} p_1 & \sqrt{p_1 p_2} o_{1,2} & \cdots & \sqrt{p_1 p_k} o_{1,k} \\ 0 & p_2 & \cdots & \sqrt{p_2 p_k} o_{2,k} \\ \vdots & \vdots & \ddots & \vdots \\ 0 & 0 & \cdots & p_k \end{bmatrix}. \tag{13}
$$

Because $\boldsymbol{L}$ is a triangular matrix with non-zero diagonal elements, it is invertible and its determinant $\det(\boldsymbol{L})$ is non-zero. By noting that $\det(\boldsymbol{\Sigma}) = \det(\boldsymbol{L}\boldsymbol{L}^{\mathrm{T}}) = \det(\boldsymbol{L}) \times \det(\boldsymbol{L}^{\mathrm{T}}) = [\det(\boldsymbol{L})]^2$, we see that the upper-triangular decomposition of $\boldsymbol{\Sigma}$ gives us a $\boldsymbol{\Sigma}$ that has a non-zero determinant and is therefore invertible.

Finally, it is possible to recover the values of parameters in the canonical expression of $\boldsymbol{\Sigma}$, $\boldsymbol{\Sigma} = \mathrm{diag}(\sqrt{\boldsymbol{r}}) \times \boldsymbol{\rho} \times \mathrm{diag}(\sqrt{\boldsymbol{r}})$. We first realize that:

$$\boldsymbol{\Sigma}_{ij} = \sum_{x=1}^{k} \boldsymbol{L}_{ix}(\boldsymbol{L}^{\mathrm{T}})_{xj}, \tag{14}$$

from which we can obtain the $i$-th character's evolutionary rates, $r_i$, and the correlation of that character with any other character $j$, $\rho_{ij}$:

$$r_i = \sqrt{\boldsymbol{\Sigma}_{ii}}, \tag{15}$$

and

$$\rho_{ij} = \frac{\boldsymbol{\Sigma}_{ij}}{\sqrt{\boldsymbol{\Sigma}_{ii}\boldsymbol{\Sigma}_{jj}}}. \tag{16}$$

##### 3.3 Can $\boldsymbol{\Sigma}$ be inverted?

In this section, we will briefly expand on a claim made in the main text, namely, that if characters outnumber species significantly, i.e., $k >> n$, the character variance-covariance matrix $\boldsymbol{\Sigma}$ cannot be inverted. As discussed in the previous section, we must invert $\boldsymbol{\Sigma}$ to evaluate the likelihood of the phylogenetic BM model. Our treatment will be short and informal.

Let us assume the parameterization described in the main text, which makes use of between-character correlation values in $\boldsymbol{\rho}$ and character-specific evolutionary rates in $\boldsymbol{r}$. We start with a $\boldsymbol{\Sigma}$ matrix put together from user-specified $\boldsymbol{\rho}$ and $\boldsymbol{r}$. This matrix is referred generally as the "sample covariance matrix", and the reader might recognize it from its canonical formulation:

$$\boldsymbol{\Sigma} = \frac{1}{(n-1)}(\mathbf{M} - \mathbf{1}\overline{\boldsymbol{m}})^{\mathrm{T}}(\mathbf{M} - \mathbf{1}\overline{\boldsymbol{m}}), \tag{17}$$

where superscript "T" denotes vector or matrix transpose, $\mathbf{M}$ is the $n \times k$ data matrix in Eq. 4, $\mathbf{1}$ is an $1 \times k$ row vector of 1's, and $\overline{\boldsymbol{m}} = \{m_j : 1 \leq j \leq k\})$ is the sample mean $k \times 1$ (column) vector, with:

$$m_j = \frac{1}{n}\sum_{i=1}^{n} m_{ij} \tag{18}$$

Being able to invert $\boldsymbol{\Sigma}$ is contingent upon this matrix being full rank, but before we examine its rank, we will conveniently rewrite Eq. 17 as:

$$\boldsymbol{\Sigma} = \frac{1}{(n-1)}\mathbf{Z}\mathbf{Z}^{\mathrm{T}}, \tag{19}$$

with $\mathbf{Z}$ denoting the data matrix after subtracting $\overline{\boldsymbol{m}}$ from it column-wise, that is, after centering the observed values for each of the $k$ random variables (i.e., each character in the phylogenetic BM model). In other words, we will make it so for each character $k$, $\sum_{i=1}^{n} z_{ik} =$

0. This constraint means that once $(n-1)$ rows of $\mathbf{Z}$ are known, we can deduce what the last row is.

One can now remember that, given an $n \times k$ matrix $\mathbf{A}$, $\text{rank}(\mathbf{A}) = \min(n, k)$, and that $\text{rank}(\mathbf{A}) = \text{rank}(\mathbf{A}\mathbf{A}^{\text{T}})$. From these two properties, the constraint mentioned above, and Eq. (19), we can see that $\text{rank}(\mathbf{Z}) = \min(n-1, k)$. Now if $k > (n-1)$, then $\text{rank}(\mathbf{\Sigma}) = \min(n-1, k) = (n-1)$, and $\mathbf{\Sigma}$ becomes rank-deficient and non-invertible – we cannot evaluate the likelihood under the phylogenetic BM model.

#### 4. Data transformation

Given a constant $\mathbf{\Sigma}$ – e.g., characters are assumed to be independent, or an estimate of $\boldsymbol{\rho}$ is used – one way to save time in the calculation of the phylogenetic BM likelihood is to decompose $\mathbf{\Sigma}$ so that $(\boldsymbol{x}_g^{\text{T}}\mathbf{\Sigma}^{-1}\boldsymbol{x}_g)$ inside the summation in Eq. (4) becomes $(\boldsymbol{x}_g^{\text{T}}\boldsymbol{H}^{\text{T}}\boldsymbol{H}\boldsymbol{x}_g)$. If we let $\boldsymbol{H}\boldsymbol{x}_g = \boldsymbol{x}'_g$, one can interpret this decomposition as a data transformation:

$$\mathbf{Z} = \mathbf{M}\boldsymbol{H}^{\text{T}}, \tag{20}$$

where the characters in $\mathbf{Z}$ are uncorrelated. We can then do a Cholesky decomposition of $\mathbf{\Sigma}$ (Álvarez–Carretero et al., 2019), $\mathbf{\Sigma} = \mathcal{L}\mathcal{L}^{\text{T}}$, for example, such that $\boldsymbol{H} = \mathcal{L}^{-1}$. Because $\mathbf{\Sigma}$ is constant, this procedure is done only once at the start of MCMC. (Note that we distinguish the Cholesky factor of $\mathbf{\Sigma}$, $\mathcal{L}$, from the upper-triangular factor $\boldsymbol{L}$ mentioned in section 3. The former, $\mathcal{L}$, is never exposed to the user in BEAST 2 and is not a parameter to be sampled.)

If we now look at the PIC formulation of the model, the summation in Eq. (4) becomes:

$$\sum_{g=n+1;u,w\in\text{Desc}(g)}^{2n} -\frac{k}{2}\ln[2\pi(t'_u + t'_w)] - \frac{1}{2}\ln|\mathbf{\Sigma}| - \frac{1}{2(t'_u + t'_w)}\boldsymbol{x}'^{\text{T}}_g\boldsymbol{x}'_g, \tag{21}$$

where $\boldsymbol{x}'_g$ is the contrast between the transformed values of $u$ and $w$, i.e., $\boldsymbol{x}'_g = \boldsymbol{z}_u - \boldsymbol{z}_w$. Now instead of carrying out a vector-matrix-vector product for each internal node $g$, we have a vector-vector product.

Math operations in Mitov and Stadler's method can also be made simpler by transforming the original data according to Eq. (20). Eq. (5) becomes:

$$\ln f(\boldsymbol{z}_u|\boldsymbol{z}_g, \mathbf{\Sigma}) = -\frac{k}{2}\ln(2\pi t_u) - \frac{1}{2}\ln|\mathbf{\Sigma}| - \frac{1}{2t_u}(\boldsymbol{z}_u - \boldsymbol{z}_g)^{\text{T}}(\boldsymbol{z}_u - \boldsymbol{z}_g), \tag{22}$$

with constants $\boldsymbol{A}$, $\boldsymbol{C}$, $\boldsymbol{E}$, $f$ and parameters $\boldsymbol{\alpha}$, $\boldsymbol{\beta}$, $\gamma$ being rewritten, correspondingly:

$$\begin{aligned} A_u &= -\frac{1}{2t_u} \\ C_u &= -\frac{1}{2t_u} \\ E_u &= \frac{1}{t_u} \\ f_u &= -\frac{k}{2}\ln(2\pi t_u) - \frac{1}{2}\ln|\mathbf{\Sigma}| \end{aligned} \tag{23}$$

$$
\begin{aligned}
{\alpha_i}^{\text{tips}} &= \sum_{u \in \text{Desc}(i)} C_u \\
{\boldsymbol{\beta}_i}^{\text{tips}} &= \sum_{u \in \text{Desc}(i)} E_u \boldsymbol{z}_u \\
{\gamma_i}^{\text{tips}} &= \sum_{u \in \text{Desc}(i)} A_u \boldsymbol{z}_u^{\text{T}} \boldsymbol{z}_u + f_u
\end{aligned} \tag{24}
$$

$$
\begin{aligned}
{\alpha_i}^{\text{non-tips}} &= \sum_{u \in \text{Desc}(i)} C_u - \frac{1}{4(A_u + \alpha_u)} \times {E_u}^2 \\
{\boldsymbol{\beta}_i}^{\text{non-tips}} &= \sum_{u \in \text{Desc}(i)} -\frac{1}{2(A_u + \alpha_u)} \times E_u \boldsymbol{\beta}_u \\
{\gamma_i}^{\text{non-tips}} &= \sum_{u \in \text{Desc}(i)} f_u + \gamma_u + \frac{k}{2} \ln(2\pi) - \frac{1}{2} \ln[(-2)(A_u + \alpha_u)^k \left|\boldsymbol{\Sigma}^{-1}\right|] \\
&\qquad - \frac{1}{4(A_u + \alpha_u)} \boldsymbol{\beta}_u^{\text{T}} \boldsymbol{\beta}_u.
\end{aligned} \tag{25}
$$

Note how terms $A_u$, $C_u$, $E_u$ in Eq. (23) are now scalars (cf. Eq. 7), and how many operations in Eq. (25) are also simpler. The final probability density of the whole normalized observed data set $\mathbf{Z}$ becomes:

$$
\ln f(\mathbf{Z}|\Phi, \boldsymbol{b}_m, c_m, \boldsymbol{\Sigma}, \boldsymbol{z}_0) = \alpha_0 {\boldsymbol{z}_0}^{\text{T}} \boldsymbol{z}_0 + {\boldsymbol{z}_0}^{\text{T}} \boldsymbol{\beta}_0 + \gamma_0, \tag{26}
$$

where $\boldsymbol{z}_0$ denotes the transformed ancestral state.

Lastly, if we want to calculate the restricted likelihood at the root, as explained in the previous sections, we must subtract the variance associated with estimating $\boldsymbol{y_0}$, the root character values:

$$
\ln f(\boldsymbol{z}_0|\boldsymbol{z}'_R) = -\frac{k}{2} \ln(2\pi t'_R) - \frac{1}{2} \ln|\boldsymbol{\Sigma}| \tag{27}
$$

where

$$
t'_R = \begin{cases} t_R + \frac{t_u t_w}{t_u + t_w}, & \text{if there is a root branch } t_R \text{ connecting it to an origin node} \\ \frac{t_u t_w}{t_u + t_w}, & \text{otherwise.} \end{cases} \tag{28}
$$

#### 5. A worked example

We took inspiration from Freckleton (2012) and included a worked example in which we compute the probability density of the phylogenetic BM model under both PIC (Felsenstein, 1973) and the general method of Mitov and Stadler (2019), for both untransformed ($\mathbf{M}$) and transformed data ($\mathbf{Z}$). Our example (Boxes 1.1 to 3.3) illustrates the calculations on a conveniently small data set (three species and two characters, i.e., $n = 3$, $k = 2$).

#### 5.1 Simulating and transforming a data set of continuous characters

We first simulate the data set we will compute the probability density of, as shown in Supplementary Fig. 1. Parameters were arbitrarily chosen ($r = 0.3$, $\rho_{12} = \rho_{21} = 0.25$) and used as input in function `mvSIM()` from the `mvMORPH` R package (Clavel et al., 2015). Note that $r$ is assumed constant between traits, and $b^z_m$ constant among each $z$ tree branch.

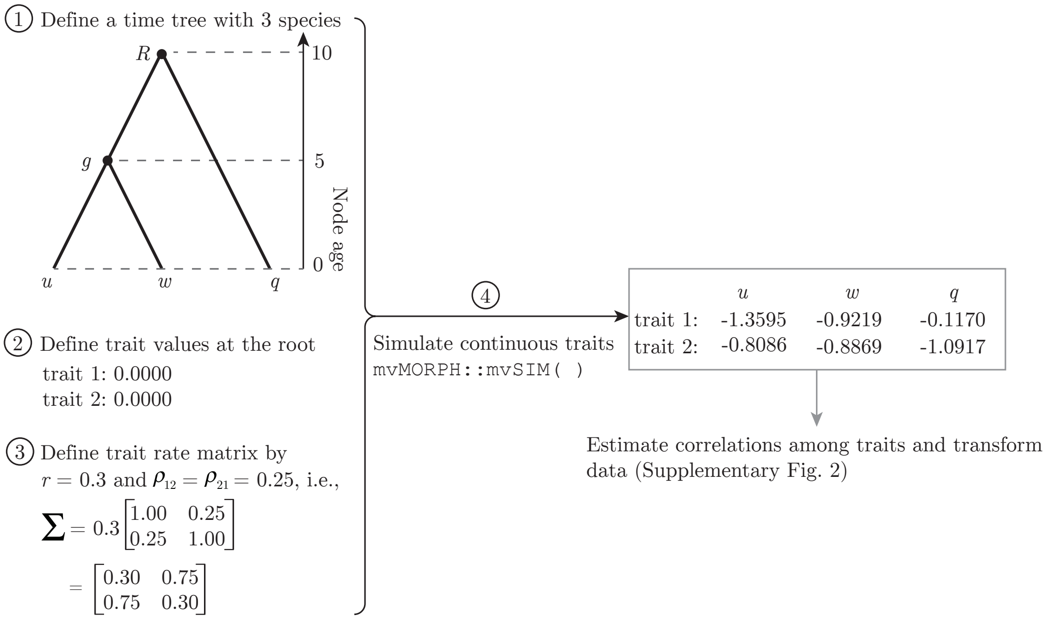


Supplementary Figure 1. Simulation of two continuous traits on a three-taxon tree (for worked example).

After simulation, we calculate $\boldsymbol{\rho}^*$ as described in the main text (note that for convenience sake, we estimate $\hat{\boldsymbol{\rho}}$ from the data observed at the tips), and transform the data as outlined above (Supplementary Fig. 2).

#### 5.2 Evaluating the probability density under the phylogenetic BM model

With both untransformed ($\mathbf{M}$) and transformed ($\mathbf{Z}$) data at hand, we now move on to calculating their probability densities under the phylogenetic BM model. Because the two methods we illustrate here (PIC and Mitov and Stadler, 2019's) are dynamic-programming algorithms on phylogenetic trees, we split the worked example into a "first prune" and "second prune" stages. The first prune performs calculations on data from species $u$ and $w$ (Boxes 1.1 to 1.3), while the second prune uses quantities calculated for the parent of $u$ and $w$, $g$, and species $q$ (Boxes 2.1 to 2.3).

Note that in the following sections, we calculate the restricted likelihood using PIC ("dropping the estimation of $[y_0]$"), and $\boldsymbol{\Sigma} = r\boldsymbol{\rho}^* = \begin{bmatrix} 0.3000 & -0.2237 \\ -0.2237 & 0.3000 \end{bmatrix}$ (Supplementary Fig. 2).

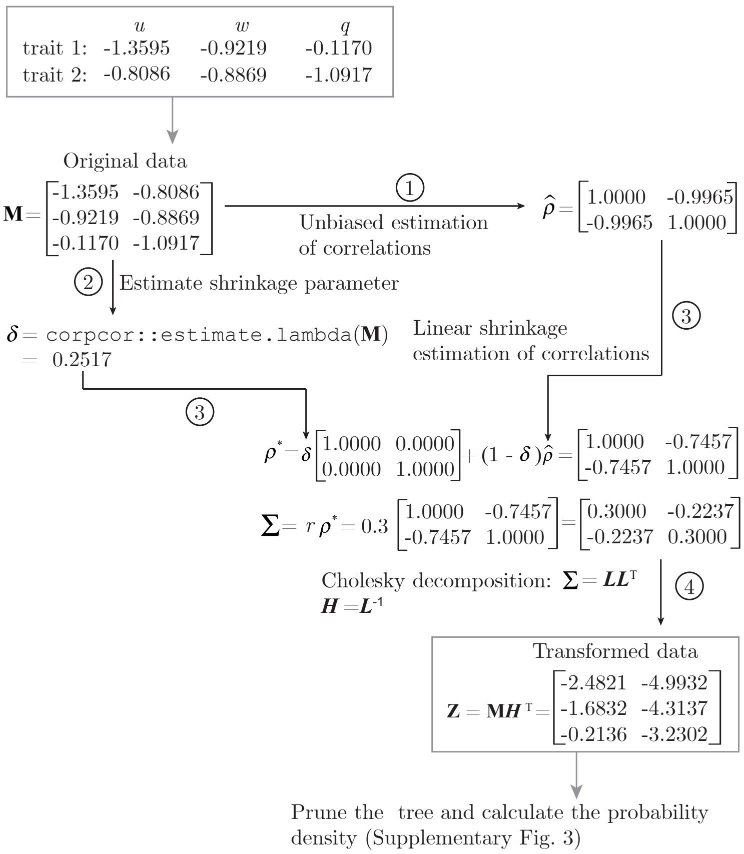


Supplementary Figure 2. Estimation of $\boldsymbol{\rho}^*$ and data transformation.

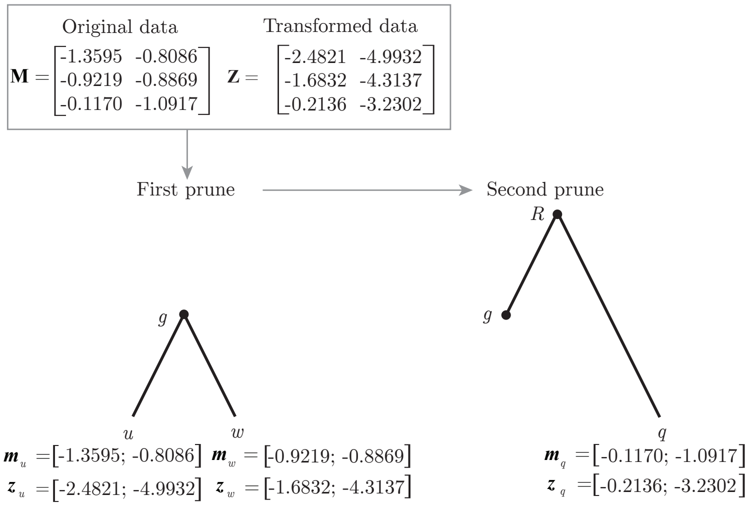


Supplementary Figure 3. On a three-taxon tree, dynamic-programming algorithms take two pruning stages in order to calculate the probability density of continuous trait data.

*First prune stage* This stage is documented in Boxes 1.1 to 1.3.

**Box 1.1: PIC with original data**

① Compute the trait-value contrasts for species $u$ and $w$ (Eq. (3a)):

$$\boldsymbol{x}_g = \boldsymbol{m}_u - \boldsymbol{m}_w = \begin{bmatrix} -1.3595 \\ -0.8086 \end{bmatrix} - \begin{bmatrix} -0.9219 \\ -0.8869 \end{bmatrix} = \begin{bmatrix} -0.4376 \\ 0.0783 \end{bmatrix}$$

② Compute the log-probability density of node $g$'s contrasts (Eq. (1)):

$$\begin{aligned} \ln f_g(\boldsymbol{x}_g|\Phi, \boldsymbol{b}_m, c_m, \boldsymbol{\Sigma}) &= -\frac{k}{2}\ln[2\pi] - 0.5\ln|(t_u+t_w)\boldsymbol{\Sigma}| - \frac{1}{2}\boldsymbol{x}_g{}^{\mathrm{T}}((t_u+t_w)\boldsymbol{\Sigma})^{-1}\boldsymbol{x}_g \\ &= -\frac{k}{2}\ln[2\pi(t_u+t_w)] - 0.5\ln|\boldsymbol{\Sigma}| - \frac{1}{2(t_u+t_w)}\boldsymbol{x}_g{}^{\mathrm{T}}(\boldsymbol{\Sigma})^{-1}\boldsymbol{x}_g \\ &= -\frac{2}{2}\ln[2\pi(5+5)] - 0.5\ln(0.03997) \\ &\quad - \frac{1}{2(5+5)}\begin{bmatrix} -0.4376 \\ 0.0783 \end{bmatrix}^{\mathrm{T}} \begin{bmatrix} 7.5074 & 5.5979 \\ 5.5979 & 7.5074 \end{bmatrix} \begin{bmatrix} -0.4376 \\ 0.0783 \end{bmatrix} \\ &= -2.5855 \end{aligned}$$

**Box 1.2: PIC with transformed data**

① Compute the trait-value contrasts for species $u$ and $w$ (Eq. (3a), but with transformed data):

$$\boldsymbol{x}'_g = \boldsymbol{z}_u - \boldsymbol{z}_w = \begin{bmatrix} -2.4821 \\ -4.9932 \end{bmatrix} - \begin{bmatrix} -1.6832 \\ -4.3137 \end{bmatrix} = \begin{bmatrix} -0.7989 \\ -0.6795 \end{bmatrix}$$

② Compute the log-probability density of node $g$'s contrasts (Eq. (1), but with transformed contrasts):

$$\begin{aligned} \ln f_g(\boldsymbol{x}'_g|\Phi, \boldsymbol{b}_m, c_m, \boldsymbol{\Sigma}) &= -\frac{k}{2}\ln[2\pi] - 0.5\ln|(t_u+t_w)\boldsymbol{\Sigma}| - \frac{1}{2(t_u+t_w))}\boldsymbol{x}'_g{}^{\mathrm{T}}\boldsymbol{x}'_g \\ &= -\frac{k}{2}\ln[2\pi(t_u+t_w)] - 0.5\ln|\boldsymbol{\Sigma}| - \frac{1}{2(t_u+t_w)}\boldsymbol{x}'_g{}^{\mathrm{T}}\boldsymbol{x}'_g \\ &= -\frac{2}{2}\ln[2\pi(5+5)] - 0.5\ln(0.03997) \\ &\quad - \frac{1}{2\times(5+5)}\begin{bmatrix} -0.7989 \\ -0.6795 \end{bmatrix}^{\mathrm{T}} \begin{bmatrix} -0.7989 \\ -0.6795 \end{bmatrix} \\ &= -2.5855 \end{aligned}$$

##### Box 1.3: Mitov and Stadler (2019)'s method with transformed data

① Compute constants $A$, $C$, $E$ and $f$ for species $u$ and $w$ (Eq. (23)):

$$A_u = -\tfrac{1}{2t_u} = -0.1 \quad C_u = -\tfrac{1}{2t_u} = -0.1 \quad E_u = \tfrac{1}{t_u} = 0.2$$
$$f_u = -\tfrac{2}{2}\ln(2\pi t_u) - 0.5\ln|\mathbf{\Sigma}| = -1.8374$$

$$A_w = -\tfrac{1}{2t_w} = -0.1 \quad C_w = -\tfrac{1}{2t_w} = -0.1 \quad E_w = \tfrac{1}{t_w} = 0.2$$
$$f_w = -\tfrac{2}{2}\ln(2\pi t_w) - 0.5\ln|\mathbf{\Sigma}| = -1.8374$$

---

② Compute parameters $\alpha$, $\boldsymbol{\beta}$ and $\gamma$ for node $g$ (Eq. (24)):

$$\alpha_g = C_u + C_w = -0.2$$

$$\boldsymbol{\beta_g} = E_u\boldsymbol{z}_u + E_w\boldsymbol{z}_w = 0.2\begin{bmatrix} -2.4821 \\ -4.9932 \end{bmatrix} + 0.2\begin{bmatrix} -1.6832 \\ -4.3137 \end{bmatrix} = \begin{bmatrix} -0.8331 \\ -1.8614 \end{bmatrix}$$

$$\begin{aligned}\gamma_g &= \left[A_u\boldsymbol{z}_u{}^{\mathrm{T}}\boldsymbol{z}_u + f_u\right] + \left[A_w\boldsymbol{z}_w{}^{\mathrm{T}}\boldsymbol{z}_w + f_w\right] \\ &= -0.1 \times \begin{bmatrix} -2.4821 \\ -4.9932 \end{bmatrix}^{\mathrm{T}}\begin{bmatrix} -2.4821 \\ -4.9932 \end{bmatrix} - 1.8374 \\ &- 0.1 \times \begin{bmatrix} -1.6832 \\ -4.3137 \end{bmatrix}^{\mathrm{T}}\begin{bmatrix} -1.6832 \\ -4.3137 \end{bmatrix} - 1.8374 \\ &= -8.9282\end{aligned}$$

*Second prune stage* This stage is documented in Boxes 2.1 to 2.3.

**Box 2.1: PIC with original data**

① Compute the expected trait values at node $g$ (Eq. (3c)):

$$\boldsymbol{m}'_g = \frac{t_u \boldsymbol{m}_w + t_w \boldsymbol{m}_u}{t_u + t_w} = \frac{5\begin{bmatrix} -1.3595 \\ -0.8086 \end{bmatrix} + 5\begin{bmatrix} -0.9219 \\ -0.8869 \end{bmatrix}}{5+5} = \begin{bmatrix} -1.1407 \\ -0.8478 \end{bmatrix}$$

② Compute the trait-value variance at node $g$ (Eq. (3d)):

$$t'_g = t_g + \frac{t_u t_w}{t_u + t_w} = 5 + \frac{5\times 5}{5+5} = 7.5$$

③ Compute the trait-value contrasts for species $v$ and internal node $g$ (Eq. (3a)):

$$\boldsymbol{x}_R = \boldsymbol{m}'_g - \boldsymbol{m}_q = \begin{bmatrix} -1.1407 \\ -0.8478 \end{bmatrix} - \begin{bmatrix} -0.1170 \\ -1.0917 \end{bmatrix} = \begin{bmatrix} -1.0238 \\ 0.2439 \end{bmatrix}$$

④ Compute the log-probability density of root node $R$'s contrasts (Eq. (1)):

$$\begin{aligned}
\ln f_R(\boldsymbol{x}_R|\Phi, \boldsymbol{b}_m, c_m, \boldsymbol{\Sigma}) &= -\frac{k}{2}\ln[2\pi] - 0.5\ln\left|(t_q + t'_g)\boldsymbol{\Sigma}\right| - \frac{1}{2}\boldsymbol{x}_R{}^{\mathrm{T}}((t_q + t'_g)\boldsymbol{\Sigma})^{-1}\boldsymbol{x}_R \\
&= -\frac{k}{2}\ln\left[2\pi(t_q + t'_g)\right] - 0.5\ln|\boldsymbol{\Sigma}| - \frac{1}{2(t_q + t'_g)}\boldsymbol{x}_R{}^{\mathrm{T}}(\boldsymbol{\Sigma})^{-1}\boldsymbol{x}_R \\
&= -\frac{2}{2}\ln[2\pi(10 + 7.5)] - 0.5\ln(0.03996) \\
&\quad - \frac{1}{2(10+7.5)}\begin{bmatrix} -1.0238 \\ 0.2439 \end{bmatrix}^{\mathrm{T}}\begin{bmatrix} 7.5074 & 5.5979 \\ 5.5979 & 7.5074 \end{bmatrix}\begin{bmatrix} -1.0238 \\ 0.2439 \end{bmatrix} \\
&= -3.2478
\end{aligned}$$

**Box 2.2: PIC with transformed data**

① Compute the expected trait values at node $g$ (Eq. (3c), but with transformed data), to be used later:

$$\boldsymbol{z}'_g = \tfrac{t_w \boldsymbol{z}_u + t_u \boldsymbol{z}_w}{t_u + t_w} = \frac{5\begin{bmatrix} -2.4821 \\ -4.9932 \end{bmatrix} + 5\begin{bmatrix} -1.6832 \\ -4.3137 \end{bmatrix}}{5+5} = \begin{bmatrix} -2.0827 \\ -4.6534 \end{bmatrix}$$

② Compute the trait-value variance at node $g$ (Eq. (3d)):
$t'_g = t_g + \frac{t_u t_w}{t_u + t_w} = 5 + \frac{5 \times 5}{5+5} = 7.5$

③ Compute the trait-value contrasts for species $v$ and internal root node $R$ (Eq. (3a), but with transformed data):

$$\boldsymbol{x}'_R = \boldsymbol{z}'_g - \boldsymbol{z}_q = \begin{bmatrix} -2.0826 \\ -4.6534 \end{bmatrix} - \begin{bmatrix} -0.2136 \\ -3.2302 \end{bmatrix} = \begin{bmatrix} -1.8691 \\ -1.4233 \end{bmatrix}$$

④ Compute the log-probability density of root node $R$'s contrasts (Eq. (1), but with transformed contrasts):

$$\begin{aligned} \ln f_R(\boldsymbol{x}'_R | \Phi, \boldsymbol{b}_m, c_m, \boldsymbol{\Sigma}) &= \frac{k}{2} \ln\left[2\pi\right] - 0.5 \ln\left|(t_v + t'_g)\boldsymbol{\Sigma}\right| - \frac{1}{2(t_q + t_g{}')} \boldsymbol{x}'_R{}^{\mathrm{T}} \boldsymbol{x}'_R \\ &= \frac{k}{2} \ln\left[2\pi(t_q + t'_g)\right] - 0.5 \ln\left|\boldsymbol{\Sigma}\right| - \frac{1}{2(t_q + t'_g)} \boldsymbol{x}'_R{}^{\mathrm{T}} \boldsymbol{x}'_R \\ &= -\frac{2}{2} \ln\left[2\pi(10 + 7.5)\right] - 0.5 \ln(0.03996) \\ &\quad - \frac{1}{2 \times (10 + 7.5)} \begin{bmatrix} -1.8691 \\ -1.4233 \end{bmatrix}^{\mathrm{T}} \begin{bmatrix} -1.8691 \\ -1.4233 \end{bmatrix} \\ &= -3.2478 \end{aligned}$$

**Box 2.3: Mitov and Stadler (2019)'s method with transformed data**

① Compute constants $A$, $C$, $E$ and $f$ for internal node $g$ and species $q$ (Eq. (23)):

$$A_q = -\tfrac{1}{2t_q} = -0.05 \quad C_q = -\tfrac{1}{2t_q} = -0.05 \quad E_q = \tfrac{1}{t_q} = 0.1$$
$$f_q = -\tfrac{2}{2}\ln(2\pi t_q) - 0.5\ln|\boldsymbol{\Sigma}| = -2.5305$$

$$A_g = -\tfrac{1}{2t_g} = -0.1 \quad C_g = -\tfrac{1}{2t_g} = -0.1 \quad E_g = \tfrac{1}{t_g} = 0.2$$
$$f_g = -\tfrac{2}{2}\ln(2\pi t_g) - 0.5\ln|\boldsymbol{\Sigma}| = -1.8374$$

② Compute parameters $\alpha$, $\boldsymbol{\beta}$ and $\gamma$ for root node $R$ (Eqs. 24 and 25):

$$\alpha_R = C_q + \left[C_g - \frac{1}{4(A_g+\alpha_g)}{E_g}^2\right] = -0.05 + \left[-0.1 - \frac{1}{4(-0.1-0.2)} \times 0.2^2\right]$$
$$= -0.1167$$

$$\boldsymbol{\beta}_R = E_q\boldsymbol{z}_q + \left[-\frac{1}{2(A_g+\alpha_g)}E_g\boldsymbol{\beta}_g\right] = 0.1\begin{bmatrix}-0.2136\\-3.2302\end{bmatrix} - \frac{0.2}{2(-0.1-0.2)}\begin{bmatrix}-0.8331\\-1.8614\end{bmatrix}$$
$$= \begin{bmatrix}-0.2990\\-0.9435\end{bmatrix}$$

$$\gamma_R = \left[A_q{\boldsymbol{z}_q}^{\mathrm{T}}\boldsymbol{z}_q + f_q\right] + \Bigg\{f_g + r_g + \frac{k}{2}\ln(2\pi) - \frac{1}{2}\ln\left[(-2)(A_g+\alpha_g)^k\left|(\boldsymbol{\Sigma})^{-1}\right|\right]$$
$$-\frac{1}{4(A_g+\alpha_g)}{\boldsymbol{\beta}_g}^{\mathrm{T}}\boldsymbol{\beta}_g\Bigg\}$$
$$= -0.05\begin{bmatrix}-0.2136\\-3.2302\end{bmatrix}^{\mathrm{T}}\begin{bmatrix}-0.2136\\-3.2302\end{bmatrix} - 2.5305 - 1.8374 - 8.9282$$
$$+\frac{2}{2}\ln(2\pi) - \frac{1}{2}\ln[(-2)(-0.1-0.2)^2(25.0248)]$$
$$-\frac{1}{4(-0.1-0.2)}\begin{bmatrix}-0.8331\\-1.8614\end{bmatrix}^{\mathrm{T}}\begin{bmatrix}-0.8331\\-1.8614\end{bmatrix}$$
$$= -9.6157$$

*Return the probability density* We finish our worked example by returning the probability density under the phylogenetic BM model, as documented in Boxes 3.1 to 3.3. Note that our implementation of Mitov and Stadler (2019)'s method returns the "full" likelihood by default. In order to match the PIC restricted likelihood calculations (as implemented in

`MCMCTree`, and as returned in Boxes 3.1 and 3.2), we must (i) use the maximum-likelihood estimate of $\boldsymbol{y_0}$, $\widehat{\boldsymbol{y_0}}$, and (ii) subtract the variance term associated with estimating $\boldsymbol{y_0}$, as shown in step ② in Box 3.3.

**Box 3.1: PIC with original data**

$$\ln f(\mathbf{M}|\Phi, \boldsymbol{b}_m, c_m, \boldsymbol{\Sigma}) = \ln f_g(\boldsymbol{x}_g) + \ln f_R(\boldsymbol{x}_R) = -2.5855 - 3.2478 = -5.8334$$

**Box 3.2: PIC with transformed data**

$$\ln f(\mathbf{Z}|\Phi, \boldsymbol{b}_m, c_m, \boldsymbol{\Sigma}) = \ln f_g(\boldsymbol{x}'_g) + \ln f_R(\boldsymbol{x}'_R) = -2.5855 - 3.2478 = -5.8334$$

**Box 3.3: FLC with transformed data**

① Compute the expected trait values at root node $R$ (Eq. (3c), but with transformed data):

$$\boldsymbol{z}'_R = \frac{t_q \boldsymbol{z}'_g + t'_g \boldsymbol{z}_q}{t_q + t'_g} = \frac{10\begin{bmatrix} -2.0826 \\ -4.6534 \end{bmatrix} + 7.5\begin{bmatrix} -0.2136 \\ -3.2302 \end{bmatrix}}{10+7.5} = \begin{bmatrix} -1.2816 \\ -4.0435 \end{bmatrix}$$

② Compute trait-value variance at root node $R$ (Eq. (28)):

$$t'_R = \frac{t'_g t_q}{t'_g + t_q} = \frac{10 \times 7.5}{10+7.5} = 4.2857$$

③ Compute full log-probability density, using $\widehat{\boldsymbol{y_0}} = \boldsymbol{z}'_R$ (Eq.(26)):

$$\begin{aligned}
\ln f(\mathbf{Z}|\Phi, \boldsymbol{b}_m, c_m, \boldsymbol{\Sigma}, \boldsymbol{z}'_R) &= \alpha_R \boldsymbol{z}'^{\mathrm{T}}_R \boldsymbol{z}'_R + \boldsymbol{z}'^{\mathrm{T}}_R \boldsymbol{\beta}_R + \gamma_R \\
&= -0.3888 \times \begin{bmatrix} -1.2816 \\ -4.0435 \end{bmatrix}^{\mathrm{T}} \begin{bmatrix} -1.2816 \\ -4.0435 \end{bmatrix} \\
&+ \begin{bmatrix} -1.2816 \\ -4.0435 \end{bmatrix}^{\mathrm{T}} \begin{bmatrix} -0.2990 \\ -0.9435 \end{bmatrix} - 9.6157 \\
&= -7.5166
\end{aligned}$$

④ Compute restricted probability density:

$$\begin{aligned}
\ln f(\mathbf{Z}|\Phi, \boldsymbol{b}_m, c_m, \boldsymbol{\Sigma}, \boldsymbol{z}'_R) &= \alpha_R \boldsymbol{z}'^{\mathrm{T}}_R \boldsymbol{z}'_R + \boldsymbol{z}'^{\mathrm{T}}_R \boldsymbol{\beta}_R + \gamma_R - \left(-\frac{k}{2}\ln(2\pi) - 0.5\ln|t'_R \boldsymbol{\Sigma}|\right) \\
&= -7.5166 - \left(-\frac{k}{2}\ln(2\pi t'_R) - 0.5\ln|\boldsymbol{\Sigma}|\right) \\
&= -7.5166 - \left(-\frac{2}{2}\ln(2\pi \times 4.2857) - 0.5\ln(0.03996)\right) \\
&= -5.8334
\end{aligned}$$

### 6. Model validation and benchmarking

#### 6.1 Graphical representation of Bayesian networks

The models we use to simulate data and carry out inference belong to the class of models known as Bayesian networks, which are probabilistic graphical models that can be represented by a directed acyclic graph (DAG). In phylogenetics, this graphical representation of Bayesian networks was introduced by Höhna et al. (2014).

Unlike the more streamlined graphical representation used in Höhna et al. (2014) (and in subsequent work by those authors and their collaborators), in the present study we chose to use a more explicit and general graphical representation known as a factor graph (Frey, 2002). The main difference is that factors graphs have function nodes defining the relationships between variables in the model. In our case (Supplementary Figs. 4 and 18), these function nodes represent:

1. Sampling distributions (denoted by filled squares), e.g., an “FBD” node representing a fossilized birth-death process we use to sample a phylogenetic tree;
2. Deterministic functions (denoted filled diamonds), e.g., a strict clock that deterministically scales a time tree into a tree measured in expected character changes over time. This latter output node is then (redundantly) represented by a red diamond.

We attempted to keep our notation as semantically consistent as possible, as well as consistent with the written formulation of the integrative model (Eq. (2.6) in the main text). For example, we call all global clock rates “c”, and use subscripts to differentiate between the continuous morphological (subscript $m$), discrete morphological (subscript $d$) and molecular (subscript $s$) rates. One downside of doing so is that subscripts indicating data types may be confused with subscripts indicating elements in a container. We thus note that, unless explicitly specified within a plate (a rounded rectangle representing a container in the factor graph ) – e.g., $\forall i \in \{1, 2, ..., n\}$ – subscripts denote either categorical classes such as data types (e.g., $d$, $m$, $s$), or a random variable a set is associated to (e.g., $\theta_\Phi$ denotes the set of parameters related to the FBD process, which generates $\Phi$).

In our factor graphs we do not represent the structure of phylogenetic trees explicitly (see Höhna et al., 2014, for a discussion of this possibility), by plating (see above) their terminal and internal nodes. This means that despite certain nodes being depicted as conditionally independent given the tree ($\mathbf{M}_{ij}$ and $\mathbf{D}_{ij}$, or $\boldsymbol{b}_m$ and $\boldsymbol{b}_d$; Supplementary Fig. 18), they will in fact be constrained in terms of the values they may take. If a given species represented by index $i$ is scored for both discrete and continuous morphology, for example, $\mathbf{M}$ and $\mathbf{D}$ will be constrained to have specific values at $i$. These are complexities hidden away by the simpler notation we chose; we do believe, however, that this simpler presentation makes the main structure of the model clearer.

#### 6.2 Simulation study I

We performed a coverage validation study to verify the correctness of our implementation of Mitov and Stadler (2019)'s model. A model is correct and said to be well-calibrated if 95% HPD intervals on its parameters cover (i.e., include) simulated “true” values approximately 95% of the time (Dawid, 1982). Alternatively, one can say a correct model has appropriate coverage. Simulation and inference must be done under the same model, which includes prior distributions.

The model configuration we validated is presented in its graphical form in Supplementary Fig. 4, with all parameters and prior distributions described in Table S1 and Table S2. Briefly, we assumed the fossilized birth-death (FBD; Gavryushkina et al., 2014a) tree prior for the species tree, and a strict morphological clock model. We discarded simulated

Table S1. Fossilized birth-death (FBD) parameter priors. LN: Log-normal.

| Description | Parameter | Prior distribution |
|---|---|---|
| Net diversification rate | $d = \lambda - \mu$ | LN(-0.65, 0.1) |
| Turnover rate | $v = \frac{\mu}{\lambda}$ | Beta(10, 40) |
| Sampling proportion | $s = \frac{\psi}{\mu+\psi}$ | Beta(20, 40) |

Table S2. Phylogenetic BM parameter priors ($i, j \in \{1, 2, 3, 4\}$ and $i \neq j$). LN: Log-normal; U: Uniform; N: Normal. See section 3 for the definition of $o_{ij}$ and $p_i$.

| Description | Parameter | Prior distribution |
|---|---|---|
| Off-diagonal elements associated with correlations between trait $i$ and $j$ | $o_{ij}$ | U(-1, 1) |
| Diagonal elements associated with $i$ trait | $p_i$ | LN(1.0, 0.3) |
| Root value of $i$-th trait | $y_0^i$ | N(0.0, 2.0) |

trees with (i) fewer than 10 taxa because data sets of this size are not informative, and (ii) with more than 200 taxa, for the sake of analysis runtime.

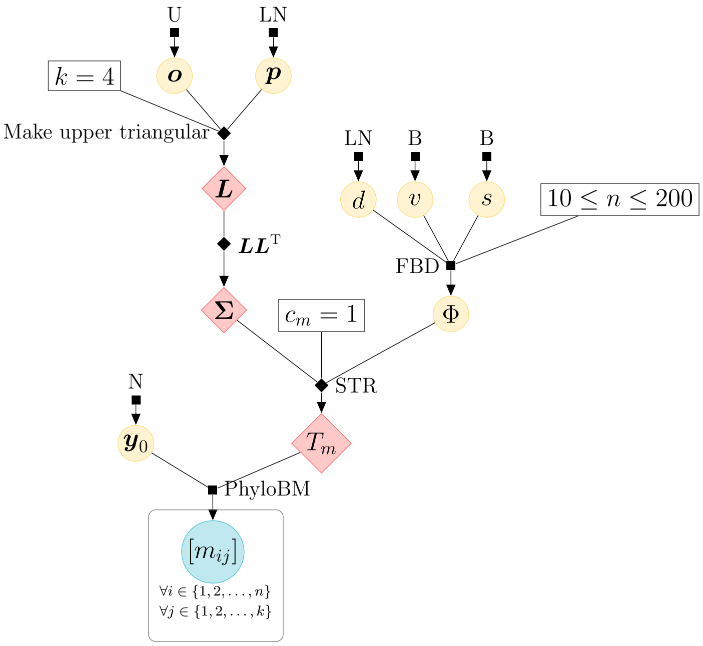


Supplementary Figure 4. Full probabilistic graphical model used in simulation study I (the coverage validation study). Filled black squares and diamonds represent sampling distributions and deterministic functions, respectively. Yellow and blue filled circles denote random variables that do not and that do have data associated to them, respectively. $T_m$ is the morphological tree measured in expected change per character after the incorporation of branch rates by a strict (STR) molecular morphological clock. $\Phi$ represents the time-scaled species tree (both topology and branch lengths). $\mathbf{M} = [m_{ij}]$; the remaining parameters are described in the text. Sampling distributions include U (uniform), N (normal), LN (log-normal), B (beta), FBD (fossilized birth-death), PhyloBM (phylogenetic Brownian motion). For more details, please see subsection “Graphical representation of Bayesian networks” above.

One hundred independent data sets were simulated using a series of programs listed

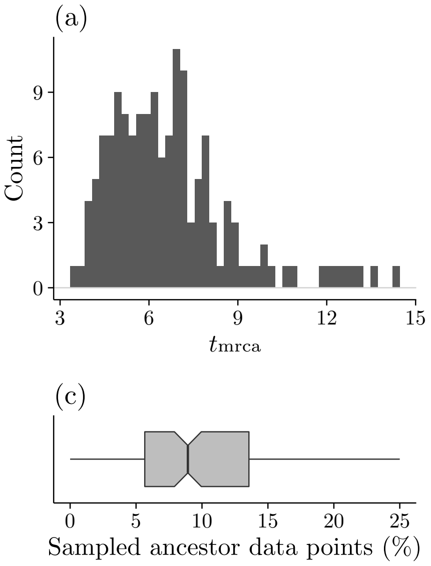


Supplementary Figure 5. Summary of tree statistics. (a) Histogram of species tree heights (most recent tip to root). (b) Histogram of origin time. (c) Percentage of observed taxa that are sampled ancestors. (d) Number of tips.

in Table S15. Those programs were written independently from BEAST 2. We simulated four continuous characters for each species. Inference was carried out with 10 million-generation chains yielding 2,000 samples (we discarded 10% of the samples as burn-in). MCMC chains convergence was diagnosed with mean effective sample sizes (ESSs) greater than 200 for all parameters (Table S3).

As expected of a correct model implementation, we observed good coverage for all parameters (Table S4). Plots of true values against mean posterior estimates (Supplementary Figs. 6 to 8) indicate most parameters could be recovered under our simulated scenarios. Parameters such as the net diversification rate and some of the ancestral character values were harder to identify, but this result has no bearing on model correctness because it is not related to coverage.

Table S3. Effective sample sizes for each parameter in coverage validation experiment. "S.A." stands for sampled ancestor.

| Parameter | Mean | Parameter | Mean |
|---|---|---|---|
| $r_1$ | 1595 | $\rho_{12},\ \rho_{21}$ | 1710 |
| $r_2$ | 1554 | $\rho_{13},\ \rho_{31}$ | 1713 |
| $r_3$ | 1570 | $\rho_{14},\ \rho_{41}$ | 1695 |
| $r_4$ | 1618 | $\rho_{23},\ \rho_{32}$ | 1695 |
| $y_0^1$ | 1718 | $\rho_{24},\ \rho_{42}$ | 1696 |
| $y_0^2$ | 1715 | $\rho_{34},\ \rho_{43}$ | 1673 |
| $y_0^3$ | 1686 | $\lambda - \mu$ | 1675 |
| $y_0^4$ | 1700 | $\frac{\psi}{\mu+\psi}$ | 1724 |
| $\frac{\mu}{\lambda}$ | 1723 | $t_{\mathrm{mrca}}$ | 1035 |
| $\mathcal{O}$ | 1660 | S.A. count | 1694 |

Table S4. Parameter 95%-HPD coverage from coverage validation experiment

| Parameter | Coverage (%) | Parameter | Coverage (%) |
|---|---|---|---|
| $r_1$ | 96 | $\rho_{12},\ \rho_{21}$ | 92 |
| $r_2$ | 95 | $\rho_{13},\ \rho_{31}$ | 94 |
| $r_3$ | 93 | $\rho_{14},\ \rho_{41}$ | 95 |
| $r_4$ | 95 | $\rho_{23},\ \rho_{32}$ | 96 |
| $y_0^1$ | 97 | $\rho_{24},\ \rho_{42}$ | 96 |
| $y_0^2$ | 94 | $\rho_{34},\ \rho_{43}$ | 98 |
| $y_0^3$ | 95 | $\lambda - \mu$ | 94 |
| $y_0^4$ | 96 | $\frac{\psi}{\mu+\psi}$ | 97 |
| $\frac{\mu}{\lambda}$ | 90 | $t_{\text{mrca}}$ | 95 |
| $\mathcal{O}$ | 92 | S.A. count | 98 |

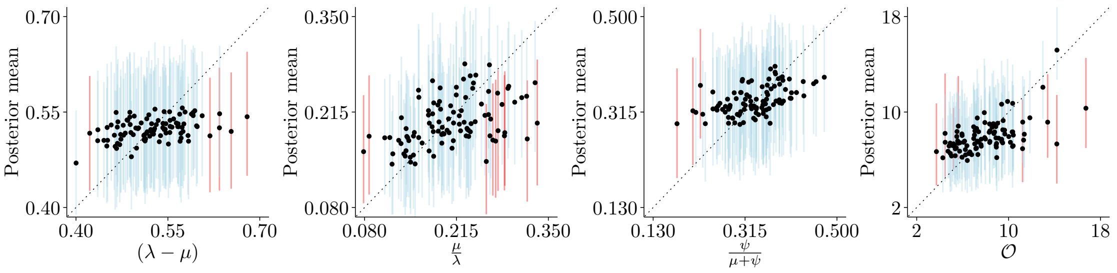


Supplementary Figure 6. Posterior means against simulated ("true") parameter values from simulation study I (100 data sets). Light blue and red vertical lines represent 95%-HPDs that do and do not include the true parameter value, respectively. Parameters are described in the main text.

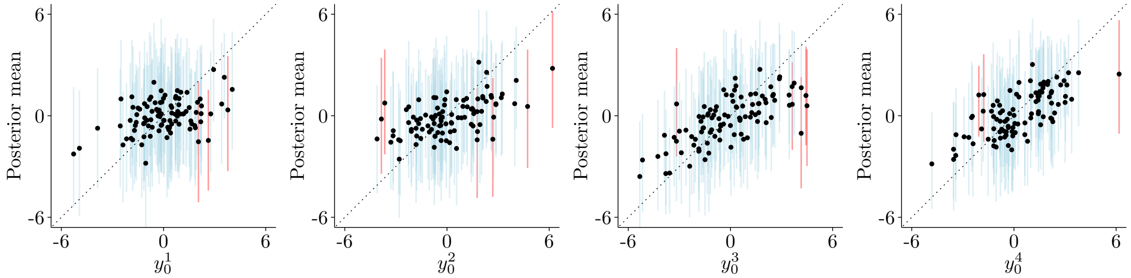


Supplementary Figure 7. Posterior means against simulated ("true") parameter values from simulation study I (100 data sets). Light blue and red vertical lines represent 95%-HPDs that do and do not include the true parameter value, respectively. Parameters are described in the main text.

#### 6.3 Simulation study II

For panels (a-c) in figure 2 in the main text, we drew an arbitrary tree from an FBD process. This tree (Supplementary Fig. 9a) had 24 taxa, out of which two were sampled (direct) ancestors of living taxa and one was a fossil leaf.

For panels (d-i), we started with a second FBD tree we refer to as "source" tree 1 (ST1; Supplementary Fig. 9b), chosen for its symmetry about the root, in terms of extant and fossil taxon count. ST1 had 42 taxa on both sides of the root, with 18 fossils on one side, and 16 on the other. By starting from ST1, we produced a derived tree ("derived tree 1", DT1) by pruning all the fossils on the right side of the root; this derived tree had 18

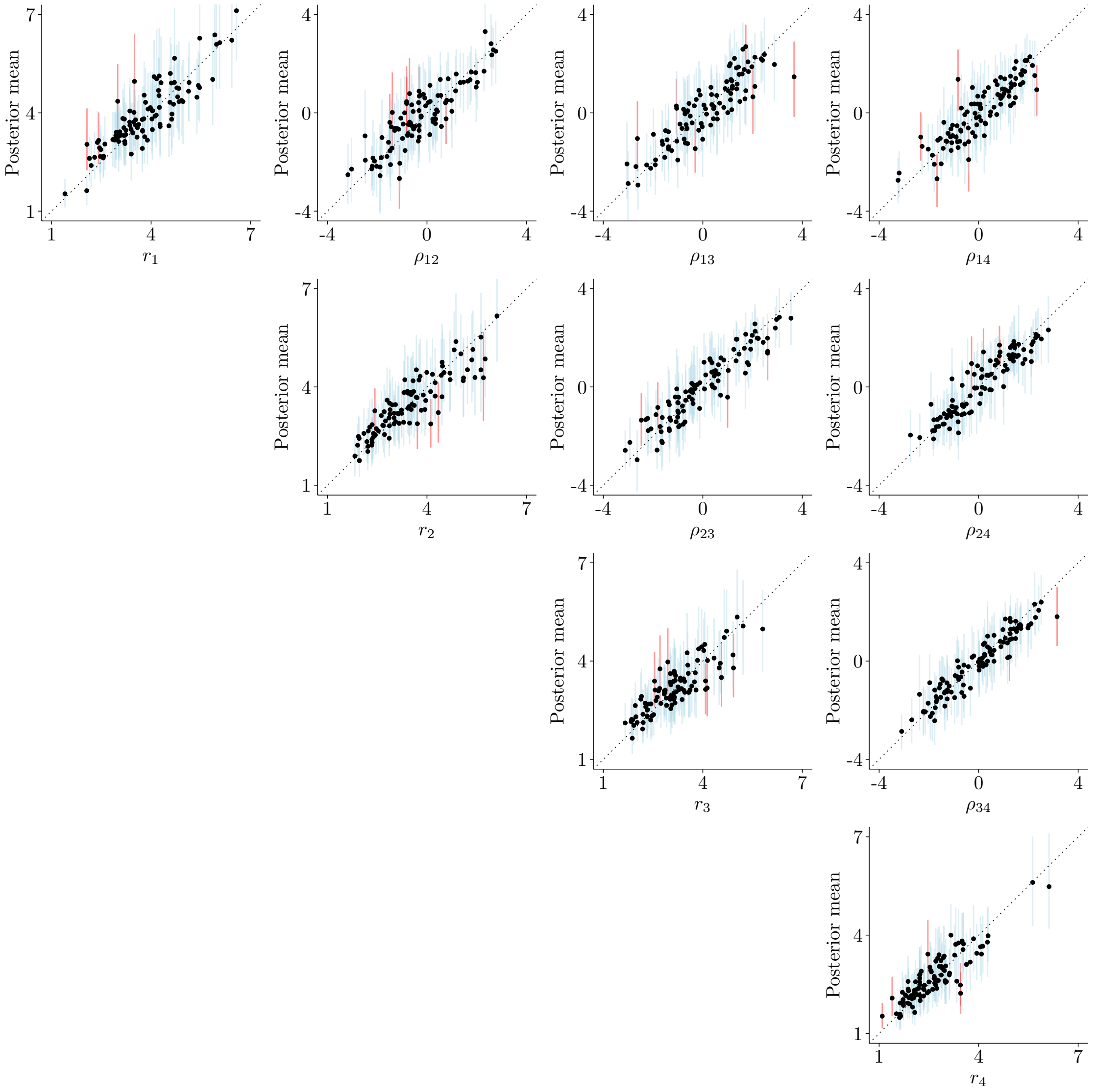


Supplementary Figure 8. Posterior means against simulated ("true") parameter values from simulation study I (100 data sets). Light blue and red vertical lines represent 95%-HPDs that do and do not include the true parameter value, respectively. Parameters are described in the main text.

fossils and was used for panels (d-f) in figure 2 from the main text. A second derived tree (DT2) was obtained by randomly selecting 9 fossils on both sides of ST1 (to a total of 18 fossils), so that the resulting DT2 (also used for panel (d) in Fig. 2) would be comparable to DT1. Lastly, we derived a third and fourth trees (DT3 and DT4), by removing the 17 oldest or 17 youngest fossils, respectively. DT3 and DT4 underlaid the comparison made in panels (g-i) in figure 2 from the main text.

We used the aforementioned trees to simulate 1, 2, 5, 10 or 50 continuous characters, replicated 20 times each. When simulating continuous characters, the state at the root, $\boldsymbol{y_0}$, was set to 0.0 for all characters; pairwise character correlations in $\boldsymbol{\rho}$ were all set to be 0.0 or 0.9. Evolutionary rates, shared by all characters and branches in all trees, were simulated from a log-normal prior with log-mean 0.0 and log-standard deviation of 0.1.

Fig. 2 in the main text shows the results of carrying out inference on the data sets simulated as described above. We estimated evolutionary rates, divergence times and fossil placement. Characters were always assumed independent during inference (i.e., the off-diagonal elements of $\boldsymbol{\rho}$ were fixed at 0.0). Consequently, in one set of analyses the model was correctly specified (when simulated characters indeed evolved independently; Fig. 2a-c, gray curve), and in the other set the model was misspecified (when simulated characters evolved in highly correlated fashion; Fig. 2a-c, black curve). The character value at the root was set to the truth, and so was the topology of the extant taxa subtree.

For each number of characters and each replicate we ran one MCMC chain with 10 million steps, logging every 5,000 states and using a burn-in of 10%. Chain convergence was assessed by observing an ESS > 200.

#### 6.4 Simulation study III

For panels (a-c) in figure 3 in the main text, we randomly drew another "source" tree (ST2; Supplementary Fig. 9c) from an FBD process. ST2 had 30 living and 30 fossil species. From ST2 we derived four different trees (accompanying this manuscript as attached supplementary image files), by randomly pruning 0, 10, 20 and 30 fossils. These four trees were then used in 20 replicated simulations (a total of 80 simulated data sets) of ten correlated characters (pairwise correlations, the off-diagonal elements of $\boldsymbol{\rho}$, were all set to 0.9). Evolutionary rates were simulated from a log-normal distribution with log-mean of 0.0 and log-standard deviation of 0.1, and shared by all characters and tree branches. The character(s) value at the root were set to 0.0. Figure 3a-c in the main text summarize the accuracy and precision of node ages estimates from the simulated data sets. The model was correctly specified in inference; we assumed pairwise character correlations to be known, and co-estimated evolutionary rates and fossil placement.

Panels (d-f) show the inference results from two sets of simulations, the first of two and the second of ten correlated continuous characters (pairwise correlations, the off-diagonal elements of $\boldsymbol{\rho}$, were all set to 0.9). When simulating continuous characters, we used the tree shown in supplementary figure 9b (ST1); evolutionary rates were shared by all characters and tree branches and drawn from a log-normal distribution with log-mean of 0.0 and log-standard deviation of 0.1. The main text describes how we simulated continuous characters so as to have different levels of model fit to data (the x-axis in panels [d-f]). Like with panels (a-c), the model was correctly specified; we assumed pairwise character correlations to be known, and estimated evolutionary rates, divergence times and fossil placement.

Along the same lines, panels (g-i) in figure 3 show the results from inferring divergence times using ten characters simulated 20 independent times on four different trees. These four trees (not shown) were derived from a third "source" tree with 50 extant and 16 fossil taxa (ST3; Supplementary Fig. 9d), drawn from an FBD process. Each tree was produced by removing increasingly larger numbers of extant taxa – 5, 15, 25, 35 – from ST3. A key difference in this experiment was that the ten characters were simulated using the upper-triangular decomposition of $\boldsymbol{\Sigma}$ described above, with the off-diagonal elements of $\boldsymbol{L}$ being drawn from a normal distribution with a mean and a standard deviation of 0.5, truncated between -1.0 and 1.0.

So in addition to estimating divergence times and fossil placement, figure 3g-i show results from also accounting for character correlations by either (i) estimating $\boldsymbol{L}$ with MCMC ($\boldsymbol{r}$ and $\boldsymbol{\rho}$ can be obtained by post-processing the MCMC chain), (ii) assuming characters evolved independently, or (iii) estimating $\boldsymbol{\rho}^*$ as described in the main text. In order to do the latter, we first obtained $\boldsymbol{\rho}$ by post-processing $\boldsymbol{\Sigma}$ and $\boldsymbol{L}$; next we simulated 100 intraspecific data points using a multivariate normal distribution with mean $\mathbf{0.0}$ and variance-covariance matrix $\boldsymbol{\rho}$. These intraspecific data points were then used during inference for obtaining $\boldsymbol{\rho}^*$.

For all panels in figure 3 in the main text, the extant subtree topology was again fixed to the truth (see previous section), and so was the character(s) value(s) at the root. Each replicate in panels (a-c) and (d-i) was analysed with one MCMC chain with 5 and 10 million steps, respectively. For all MCMC chains, we logged every 5,000 generations and used a 10% burn-in. Chain convergence was assessed by observing an ESS > 200.

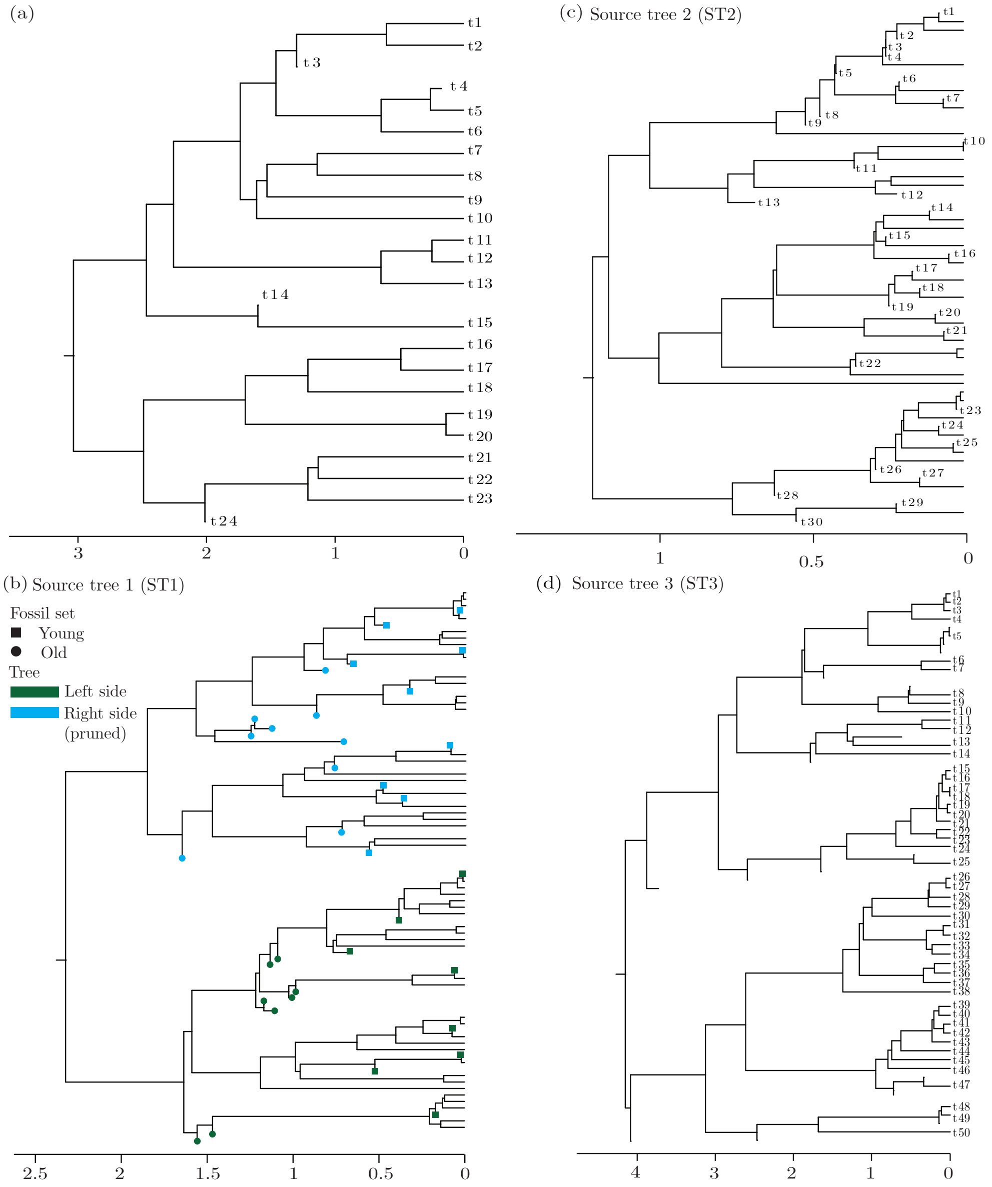


Supplementary Figure 9. Phylogenetic trees used in simulation studies II and III. The trees are random samples of a fossilised birth-death model. Trees in (a) and (b) are used in the analyses in Fig. 2 in main manuscript. Trees in (b), (c), and (d) are used in the analyses in Fig. 3 in manuscript.

#### 6.5 Simulation study IV

We benchmarked our phylogenetic BM implementation against MCMCTree's under (i) increasing number of species while holding the number of characters constant (at four; Supplementary Fig. 10), and (ii) increasing number of characters while holding the number of species constant (at 30; Supplementary Fig. 11). In both (i) and (ii), five replicate analyses were carried out for each number of taxa and number of characters, respectively. The input data sets were the same across replicates. Table S5 describes the model used to generate the input data sets.

Table S5. Parameter prior distributions for model used in benchmarking analysis. U(·) and N(·) denote uniform and normal distributions, respectively.

| Parameter | Prior distribution |
|---|---|
| $\Phi$ | Yule($\lambda = 0.2$) |
| $r$ | Gamma(20.0, 10.0) |
| $\rho_{ij}$ | U(-1.0, 1.0) |
| $y_0^i$ | N(0.0, 2.0) |

We attempted to match BEAST 2 and MCMCTree analyses to the best of our ability, which includes MCMC chain lengths and operators. For both programs, MCMC chains were run for 2.5 million generations, sampling every 1,000 and discarding the first 10% as burn-in. We used BEAST 2's `ScaleOperator` for $t_{\text{mrca}}$, `RealRandomWalkOperator` for the shared evolutionary rate, and `Uniform` operator for the internal node times. The species tree topology was kept fixed. All operators had their Bactrian version activated (Yang and Rodríguez, 2013).

We evaluated performance by observing the running time (in seconds) of the entire MCMC run as logged by each program, the ESS of the evolutionary rate shared by all traits ($r$), and the average ESS of among-character covariance parameters (which we refer to as $\bar{\boldsymbol{\rho}}$). At the moment of writing, we could not find a way to estimate among-character covariance parameters with MCMCTree (this is why this program does not have a $\bar{\boldsymbol{\rho}}$ curve in Supplementary Figs. 10 and 11).

BEAST 2's implementation outperformed MCMCTree's in all cases, both in running time and in parameter ESSs (Supplementary Figs. 10 and 11). We note that our comparisons pit not only the implementations of the likelihood classes against each other, but also those of prior distributions, operators, caching mechanisms, and logging functions.

#### 6.6 Birth-death sequential sampling (BDSS) tree model: implementation and validation

The BDSS tree model was originally introduced and implemented by Stadler and Yang (2013), and used by Álvarez–Carretero et al. (2019). We implemented and validated this model in BEAST 2* for two reasons: we wanted to be able to compare our results from those obtained with other tools (Álvarez–Carretero et al., 2019), but also make this tree model available for future analyses as an alternative to the FBD model.

Like the FBD, the birth-death-sequential-sampling (BDSS) process starts with a single lineage that can then give birth to another lineage (interpreted as speciation in our case), die (interpreted as extinction), or be sampled sequentially (sequentially here means 'not at the present moment'). Birth, death and sequential sampling events all follow Poisson processes characterized by rates $\lambda$, $\mu$ and $\psi$, respectively. In inference, the last (most recent) time is normally assumed to be the present, at which point each extant lineage is sampled

*Available on `https://github.com/fkmendes/bdtree`.

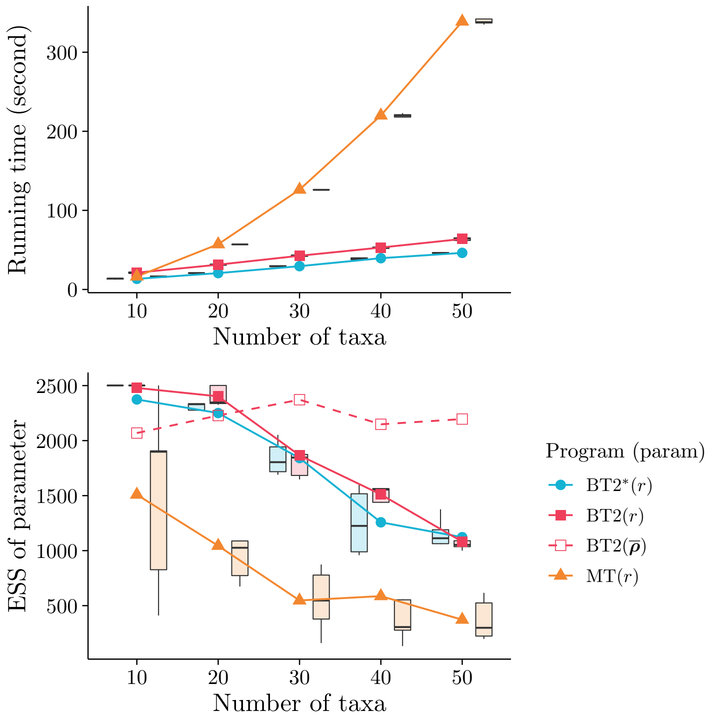


Supplementary Figure 10. Running time (top) and parameter ESSs (bottom) as a function of taxon number. BT2 and BT2* stand for BEAST 2 sampling $\boldsymbol{\rho}$ and using $\boldsymbol{\rho}^*$, respectively; MT stands for MCMCTree. $\bar{\boldsymbol{\rho}}$ denotes an average over of all among-character covariances.

with probability $\rho$ (we keep the notation consistent with that of Stadler and Yang, 2013, but note that this parameter is not to be confused with the character variance-covariance matrix of the phylogenetic BM model).

Importantly, the BDSS model differs from the FBD in that it assumes that the $N$ sampled lineages are labeled $1, \ldots, N$ from left to right (i.e., the tree is oriented). Their corresponding sampling times are denoted by $\boldsymbol{z} = \{z_1, z_2, \ldots, z_{N-1}\}$, which are also labeled from left to right. The splitting times at the internal nodes are represented by $\boldsymbol{x} = \{x_1, x_2, \ldots, x_N\}$, with the age of the root being $t_{\text{mrca}} = \max(\boldsymbol{x})$.

The BDSS model is used here as a prior distribution – within a larger hierarchical model – for the (unobservable) divergence times $\boldsymbol{x}$ of a phylogeny. In isolation, this model can be seen as treating $\boldsymbol{x}$ as "data", and is characterized by parameters $\boldsymbol{z}$, $k$, and $t_{\text{mrca}}$. This model gives us probability density:

$$f[\boldsymbol{x}|z, k, t_{\text{mrca}}] = \prod_{i=1, i\neq k}^{N-1} \frac{c_1(1-c_2)e^{-c_1 x_i}}{g(x_i)^2 \left\{ \frac{1}{g(t\text{mrca})} - \frac{1}{g(z_i^*)} \right\}}, \quad (29)$$

where $k$ is number of samples in the left subtree descending from the root, $z_i$ represents the $i$-th tip sampling time, and $x_i$ represents the splitting time of the $i$-th internal node. The remaining variables are defined as:

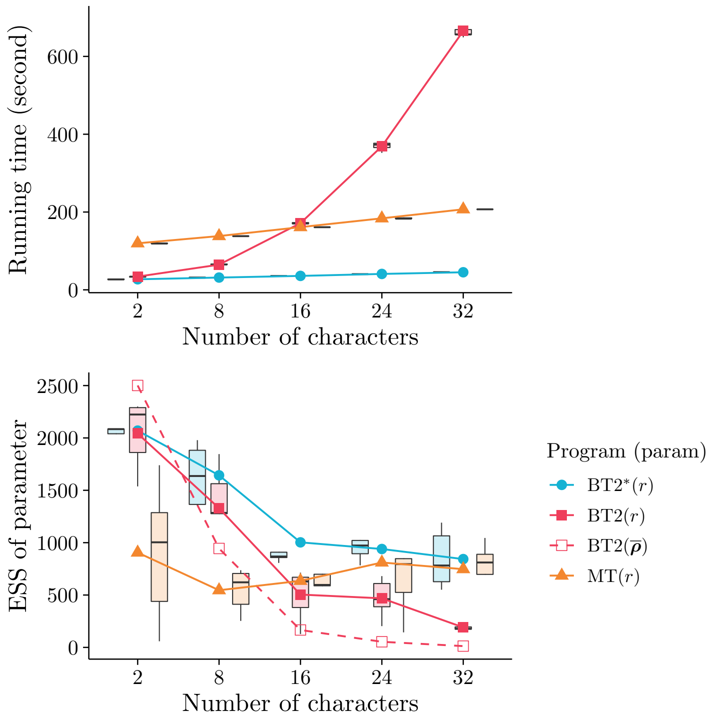


Supplementary Figure 11. Running time (top) and parameter ESSs (bottom) as a function of character number. BT2 and BT2* stand for BEAST 2 sampling $\boldsymbol{\rho}$ and using $\boldsymbol{\rho}^*$, respectively; MT stands for MCMCTree. $\bar{\boldsymbol{\rho}}$ denotes an average over of all among-character covariances.

$$\begin{aligned} z_i^* &= \max\{z_i, z_{i+1}\} \\ c_1 &= \sqrt{(\lambda - \mu - \psi)^2 + 4\lambda\psi} \\ c_2 &= -\frac{\lambda - \mu - 2\lambda\rho - \psi}{c_1} \\ g(t) &= e^{-c_1 t}(1 - c_2) + (1 + c_2), \end{aligned} \tag{30}$$

In order to match the model used by Álvarez–Carretero et al. (2019), we had to further implement soft-bounded node time priors (Yang and Rannala, 2006). The probability density of a node time $t$ assigned a uniform prior with lower and upper bounds $t_L$ and $t_U$ (keeping the original notation in Yang and Rannala, 2006), respectively, is:

$$f(t) = \begin{cases} 0.025 \times \frac{\theta_1}{t_L}\left(\frac{t}{t_L}\right)^{\theta_1 - 1} & \text{, if } 0 < t \leqslant t_L \\ \frac{0.95}{t_U - t_L} & \text{, if } t_L < t \leqslant t_U \\ 0.025 \times \theta_2 \exp\{-\theta_2(t - t_U)\} & \text{, if } t > t_U, \end{cases} \tag{31}$$

with interval $[t_L, t_U)$ consisting of the 95% HPD interval, $[0, t_L)$ having a power decay, and $t > t_U$ having an exponential decay. We set $\theta_1 = \frac{0.95 t_L}{0.025(t_U - t_L)}$ and $\theta_2 = \frac{0.95}{0.025(t_U - t_L)}$. Note that $t$ in Eq. (31) is the same as $x$ in Eq. (29).

*BDSS direct simulator* The first step in validating our BDSS implementation was to implement a direct simulator for this model. To the best of our knowledge, a BDSS direct simulator does not exist in any software platform. We must thus derive the necessary equations for carrying out the well known "inverse transform sampling" method. In a nutshell, this method requires that:

- we obtain the cumulative distribution function (*cdf*), $F_X(x)$, from the model (defining the probability density function $f_X(x)$) we are sampling a random value $x$ for variable $X$ from;
- we obtain the inverse of the *cdf*, $F'_X(x)$.

Given a known inverse *cdf* $F'_X(x)$, one can plug into it random values $x$ drawn from a standard uniform distribution – the resulting output will be equivalent to random samples from the model at hand. In our case, we note that there is one added complication: because of how the BDSS model treats (oriented) trees, the variable we are sampling (splitting times) can only take values within specific bounds (see $z_i^*$ and $p_{z_i^*}$ below).

We start by defining:

$$\begin{aligned} a_1 &= c_1(1-c_2) \\ a_2 &= \frac{1}{g(t_{\text{mrca}})} - \frac{1}{g(z_i^*)}, \end{aligned} \tag{32}$$

the probability density function of the splitting time of the $i$-th node, $X_i$, can be written as:

$$f_{X_i}(x_i) = \frac{a_1 e^{-c_1 x_i}}{a_2 g(x_i)^2}, \tag{33}$$

and the corresponding cumulative distribution function (Tanja Stadler, personal communication, 2020) as:

$$\begin{aligned} F_{X_i}(x_i) &= \frac{1}{a_2 g(x_i)} \\ &= \frac{1}{a_2[e^{-c_1 x_i}(1-c_2)+(1+c_2)]}. \end{aligned} \tag{34}$$

Indeed, we arrive at Eq. (33) by deriving Eq. (34) with respect to $x_i$:

$$\begin{aligned} \frac{\mathrm{d}}{\mathrm{d}x} F_{X_i}(x_i) &= -\frac{1}{(a_2)^2[g(x_i)]^2}(-c_1)a_2(1-c_2)e^{-c_1 x_i} \\ &= \frac{c_1(1-c_2)e^{-c_1 x_i}}{a_2[g(x_i)]^2} \\ &= \frac{a_1 e^{-c_1 x}}{a_2 g(x_i)^2} \\ &= f_{X_i}(x_i). \end{aligned} \tag{35}$$

When sampling the splitting time ($x_i$) of node $i$ from which left ($l$) and right ($r$) subtrees descend, one must respect $x_i > \max(x_{z_1}, x_{z_2}) = z_i^*$, where $z_1$ and $z_2$ are the rightmost child of $l$ and the leftmost child of $r$, respectively. This is because the BDSS model conditions not only on $t_{\text{mrca}}$ and the number of taxa, but also on tree orientation. We now use Eq. (34) to define:

$$\begin{aligned} p_{z_i^*} &= \int_{z_i^*}^{x_i} f_{X_i}(x_i)dx \\ &= F_{X_i}(x_i) - F_{X_i}(z_i^*) \\ &= \frac{1}{a_2\left[e^{-c_1 x_i}(1-c_2)+(1+c_2)\right]} - \frac{1}{a_2\left[e^{-c_1 z_i^*}(1-c_2)+(1+c_2)\right]} \end{aligned} \tag{36}$$

At this point we can derive the quantile function $F'_{X_i}(p_{z_i^*}) = x_i$, the inverse of *cdf* $F_{X_i}(x_i)$, by solving $p_{z_i^*}$ for $x_i$:

$$\begin{aligned} \frac{1}{a_2\left[e^{-c_1 x_i}(1-c_2)+(1+c_2)\right]} - c_3 &= p_{z_i^*} \\ \frac{1}{a_2\left[e^{-c_1 x_i}(1-c_2)+(1+c_2)\right]} &= p_{z_i^*} + c_3 \\ e^{-c_1 x_i}(1-c_2)+(1+c_2) &= \frac{1}{a_2(p_{z_i^*}+c_3)} \\ e^{-c_1 x_i}(1-c_2) &= \frac{1}{a_2(p_{z_i^*}+c_3)} - (1+c_2) \\ e^{-c_1 x_i} &= \frac{\frac{1}{a_2(p_{z_i^*}+c_3)} - (1+c_2)}{1-c_2} \\ e^{-c_1 x_i} &= \frac{1}{a_2(p_{z_i^*}+c_3)1-c_2} - \frac{1+c_2}{1-c_2} \\ -c_1 x_i &= \ln\left[\frac{1}{a_2(p_{z_i^*}+c_3)1-c_2} - \frac{1+c_2}{1-c_2}\right] \\ x_i &= \frac{\ln\left[\frac{1}{a_2(p_{z_i^*}+c_3)(1-c_2)} - \frac{1+c_2}{1-c_2}\right]}{(-c_1)} = F'_{X_i}(p_{z_i^*}), \end{aligned} \tag{37}$$

where:

$$c_3 = \frac{1}{a_2[e^{c_1 z_i^*}(1-c2)+(1+c2)]} \tag{38}$$

Once we have defined $F'_{X_i}(p_{z_i^*})$, direct simulation under the BDSS model follows Algorithm 1, in Section 7 below.

*Direct simulation* vs. *MCMC* We compared trees directly simulated as described above to trees sampled with MCMC using the probability density function of the BDSS model. The two models (four- and five-taxon trees) we simulated under are detailed in Table S6. We observed a close match between node times in directly simulated trees and those sampled with MCMC, under both models (Supplementary Fig. 12). This result suggests both the simulator and the probability density function have been correctly implemented.

Table S6. Model used in validation of BDSS simulator. $z_5$ is only used in the five-taxon phylogeny.

| | | | | |
|---|---|---|---|---|
| BDSS parameters | | | | |
| $\lambda = 0.5$ | $\mu = 0.3$ | $\psi = 0.001$ | $\rho = 0.1$ | |
| Node times | | | | |
| $z_1 = 13.135$ | $z_2 = 0.0$ | $z_3 = 0.0$ | $z_4 = 25.615$ | $(z_5 = 6.65)$ |
| $t_{\mathrm{mrca}} = 36$ | | | | |

*BDSS model in BEAST 2* vs. *MCMCTree* We further compared our implementation of the BDSS model against the implementation in MCMCTree by conducting an analysis analogous to a coverage validation study. Here, we simulated trees from our BDSS implementation using MCMC (100 equally spaced trees, from a total of 2,000), and then inferred parameters using MCMCTree. The initial tree at the start of the MCMC chain was random, apart from tip ages, which came from the data set in Álvarez–Carretero et al. (2019). We simulated four independent continuous characters for each individual tree. During inference, BDSS parameters ($\lambda$, $\mu$, $\psi$, $\rho$ and tip times; Table S7) were fixed to the truth, and we estimated morphological evolutionary rate, the root character value, the tree topology and the internal node ages. MCMC chains were run for 10 million generations, sampling every 5,000, to a total of 1,900 samples after discarding the first 10% as burn-in.

Appropriate coverage was obtained for all estimated parameters: (i) root age (Supplementary Fig. 13a), (ii) continuous character evolutionary rate (Supplementary Fig. 13b), and (iii) percentage of nodes (per tree in the posterior distribution) present in the true tree. These results suggest our BDSS implementation matches MCMCTree's.

Table S7. Model used in the BDSS comparison analysis, between BEAST 2 and MCMCTrees.

| | | | | |
|---|---|---|---|---|
| Node times | | | | |
| $z_1 = 13.135$ | $z_5 = 25.615$ | $z_9 = 0.0$ | $z_{13} = 0.0$ | $z_{17} = 6.65$ |
| $z_2 = 0.0285$ | $z_6 = 14.785$ | $z_{10} = 0.0$ | $z_{14} = 0.0$ | $z_{18} = 0.0285$ |
| $z_3 = 11.95$ | $z_7 = 28.55$ | $z_{11} = 0.0$ | $z_{15} = 0.0$ | $z_{19} = 0.0$ |
| $z_4 = 35.55$ | $z_8 = 0.0$ | $z_{12} = 0.0$ | $z_{16} = 0.0$ | $t_{\mathrm{mrca}} \sim \mathrm{U}(37.3, 66)$ |
| BDSS parameters | | | | |
| $\lambda = 1.0$ | $\mu = 1.0$ | $\psi = 0.001$ | $\rho = 0.0$ | |
| BM parameters | | | | |
| $r \sim \mathrm{Gamma}(20, 10)$ | $\boldsymbol{y_0} \sim \mathrm{N}(0, 2)$ | | | |

#### 6.7 Linear shrinkage estimator of $\boldsymbol{\rho}$

*6.7.1 The shrinkage (tuning) parameter $\delta$.* Given a parameter $\theta$, its estimator $\hat{\theta}$ is efficient if it has a low mean square error (MSE):

$$\mathrm{MSE}(\hat{\theta}) = \mathrm{E}[(\hat{\theta} - \theta)^2] = \mathrm{Bias}^2(\hat{\theta}) + \mathrm{Var}(\hat{\theta}) = (\mathrm{E}[\hat{\theta}] - \theta)^2 + \mathrm{Var}(\hat{\theta}) \tag{39}$$

where $\mathrm{Bias}(\hat{\theta}, \theta) = \mathrm{E}[\hat{\theta}] - \theta$ and $\mathrm{Var}(\hat{\theta}) = \mathrm{E}[(\hat{\theta} - \mathrm{E}[\hat{\theta}])^2]$. This is sometimes referred to as

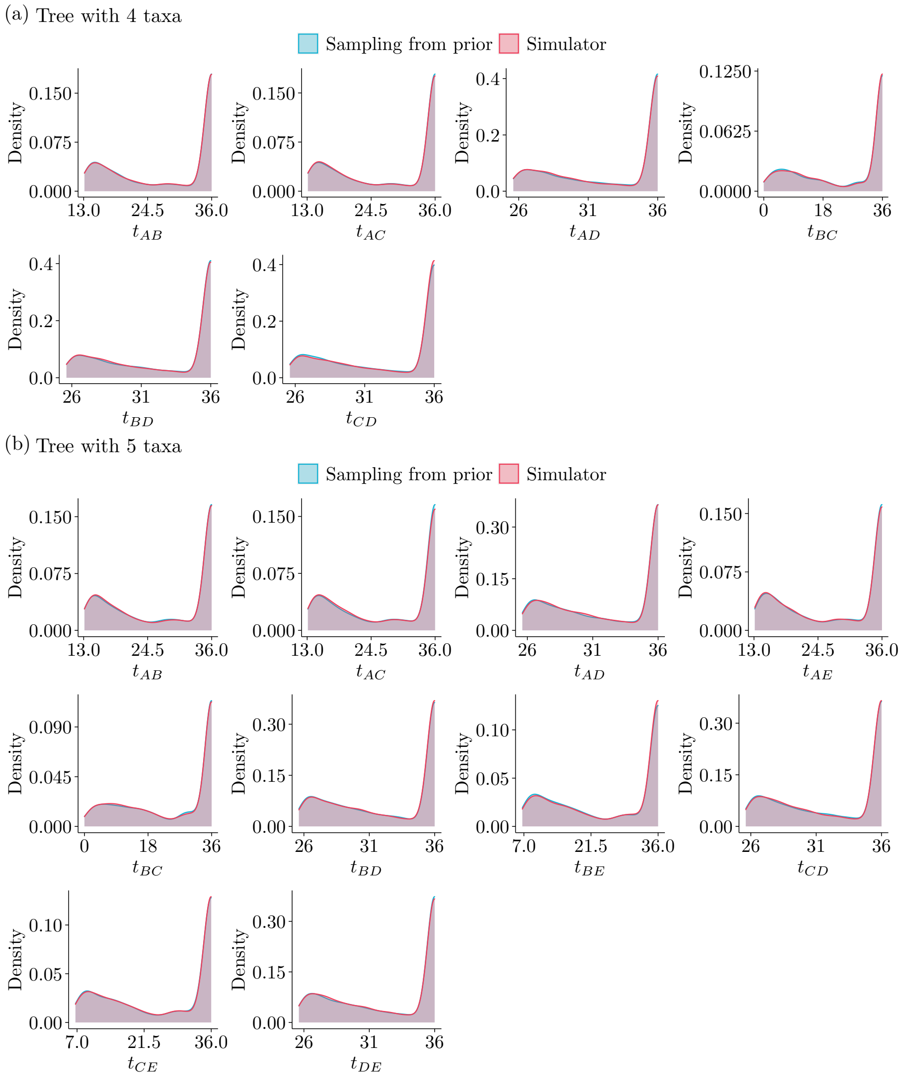


Supplementary Figure 12. Comparison of divergence times between trees sampled with MCMC (blue) and simulated directly (red). (a) In a tree with 4 taxa, there are 6 possible internal nodes with two leaf children. (b) In a tree with 5 taxa, there are 10 possible internal nodes with two leaf children.

bias-variance decomposition. The MSE of an estimator can be thus minimized by minimizing its squared bias and its variance.

In our empirical analyses described in the main text, we employ an "archetypal ridge estimator", $\boldsymbol{\rho}^*$, to approximate the among-character covariance matrix $\rho$. In the context of maximum-likelihood phylogenetics, this estimator was introduced by Clavel et al. (2019). Later, it was used in a Bayesian framework by Álvarez–Carretero et al. (2019) for divergence time estimation. Using what we described above, and if we adopt the notation used in the main text, Ledoit and Wolf (2003) and Schäfer and Strimmer (2005) defined:

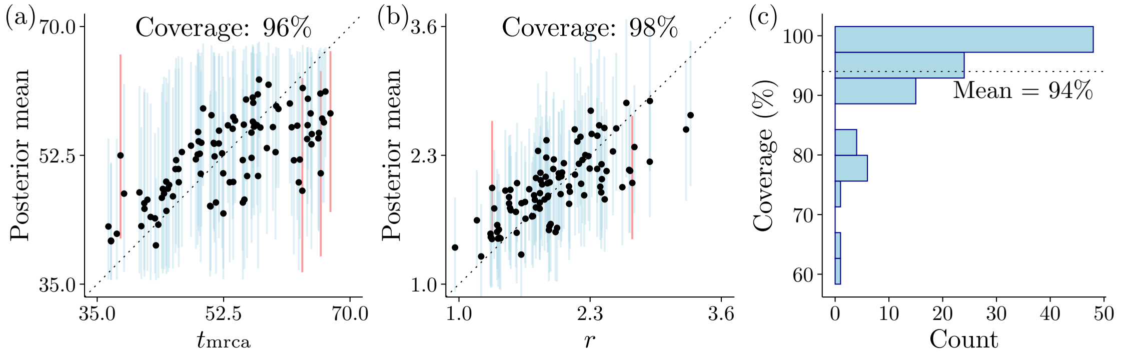


Supplementary Figure 13. Validation of BDSS likelihood implementation. (a) True root age against corresponding posterior means. (b) True evolutionary rate against corresponding posterior means. In (a) and (b) light blue and red vertical lines represent 95% HPDs that do and do not include the true parameter value, respectively. (c) Number of estimated trees where certain percent of true node times are covered in the 95% HPDs of the posterior distributions. On average, 94% of the true nodes times are covered in the 95% HPDs of the posterior distributions.

$$
\begin{aligned}
\mathrm{MSE}(\boldsymbol{\rho}^*) &= \mathrm{Bias}^2(\boldsymbol{\rho}^*) + \mathrm{Var}(\boldsymbol{\rho}^*) \\
&= \sum \left(\mathbf{E}[\boldsymbol{\rho}^*] - \boldsymbol{\rho}\right)^2 + \mathrm{Var}(\boldsymbol{\rho}^*) \\
&= \left(\mathbf{E}[\delta \boldsymbol{I} + (1-\delta)\widehat{\boldsymbol{\rho}}] - \boldsymbol{\rho}\right)^2 + \mathrm{Var}(\delta \boldsymbol{I} + (1-\delta)\widehat{\boldsymbol{\rho}}).
\end{aligned} \tag{40}
$$

At this point, we can see that how good an estimator of $\boldsymbol{\rho}^*$ is depends on what shrinkage parameter $\delta$ we use. In Bayesian analysis, we could estimate $\delta$ if we have a reasonable prior distribution for it (not the case here). Alternatively, Schäfer and Strimmer (2005) used the fact that $\widehat{\boldsymbol{\rho}}$ is an unbiased estimator, and proposed we used the $\delta$ that minimizes $\mathrm{MSE}(\boldsymbol{\rho}^*)$:

$$
\delta^* = \frac{\sum \mathrm{Var}[\widehat{\boldsymbol{\rho}}] - \mathrm{Cov}[\widehat{\boldsymbol{\rho}}, \mathbf{I}]}{\sum \mathbf{E}[(\mathbf{I} - \widehat{\boldsymbol{\rho}})^2]}. \tag{41}
$$

This is the $\delta$ we used when employing the archetypal ridge estimator.

*6.7.2 Assessing the effects of using $\boldsymbol{\rho}^*$ in inference.* Our implementation of the linear shrinkage estimator of $\boldsymbol{\rho}$, $\boldsymbol{\rho}^*$, was assessed both in terms of its behavior (as compared to the implementation in MCMCTree), and of its ability to recover correct parameter values. We simulated 100 different ten-taxon trees under the BDSS model, all with six extant and four extinct species. Fossil and root ages were arbitrarily chosen and shared by all 100 trees: $z_2 = 1.0, z_4 = 2.0, z_5 = 1.5, z_9 = 4.8$ and $t_{\mathrm{mrca}} = 10.0$, respectively. BDSS parameters were set as $\lambda = \psi = 1.0$, $\mu = 0.5$, $\rho = 0.1$. Phylogenetic BM parameter priors are listed in Table S8. We simulated four correlated continuous characters sharing the same evolutionary rate $r$, under a strict clock model.

Each of the 100 data sets was then analyzed with BEAST 2 and MCMCTree while matching the model used in simulation. The linear shrinkage estimate of $\boldsymbol{\rho}$, $\boldsymbol{\rho}^*$, was calculated from the continuous character values at all tips. (Note that the goal of our analysis is not to investigate the impact of modeling intraspecific variance under the phylogenetic BM model, as already thoroughly done by Álvarez–Carretero et al., 2019; instead, we want to

Table S8. Phylogenetic BM parameter priors used when assessing linear shrinkage estimation of $\boldsymbol{\rho}$ ($i, j \in \{1, 2, 3, 4\}$; $i \neq j$). U: Uniform; N: Normal.

| Parameter | Prior distribution |
|---|---|
| $r$ | Gamma(20.0, 10.0) |
| $\rho_{ij}$ | U(-1.0, 1.0) |
| $y_0^i$ | N(0.0, 2.0) |

investigate if we have correctly implemented the machinery that estimates $\boldsymbol{\rho}$.) Divergence times were estimated, but not the tree topology. MCMC chain lengths were set to 5 million, with samples taken every 5,000 generations and a 10% burn-in. Mean ESSs are reported in Table S9.

Table S9. Mean effective sample sizes of $r$.

| | Mean ESS |
|---|---|
| BEAST 2 sampling $\boldsymbol{\rho}$ | 885 |
| BEAST 2 with $\boldsymbol{\rho}^*$ | 881 |
| MCMCTree $\boldsymbol{\rho}^*$ | 862 |

BEAST 2 and MCMCTree yielded highly correlated $r$ posterior means (Supplementary Fig. 14), which strongly suggests the two implementations match. We ran a "control" analysis with BEAST 2 in which $\boldsymbol{\rho}$ was sampled during MCMC (i.e., we did not use the linear shrinkage estimate; Supplementary Fig. 15a). As expected, this "control" analysis produced appropriate coverage for $r$, close to 95%. On the other hand, using $\boldsymbol{\rho}^*$ reduced coverage similarly for both software platforms (Supplementary Fig. 15b,c). The moderate reduction in coverage (to approximately 84%) was caused by either under- or overestimation of $r$.

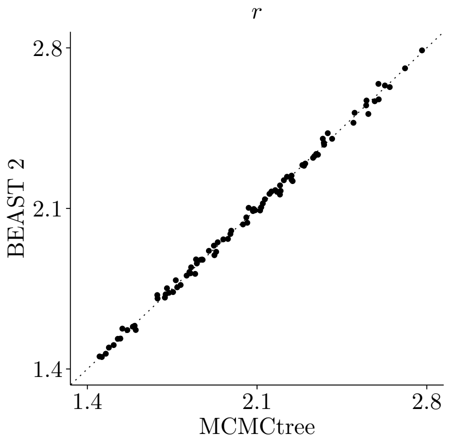


Supplementary Figure 14. Posterior mean estimates of $r$ (from 100 simulated data sets) obtained with BEAST 2 and MCMCTree, when using $\boldsymbol{\rho}^*$. In this model, four characters were used, all sharing the same $r$.

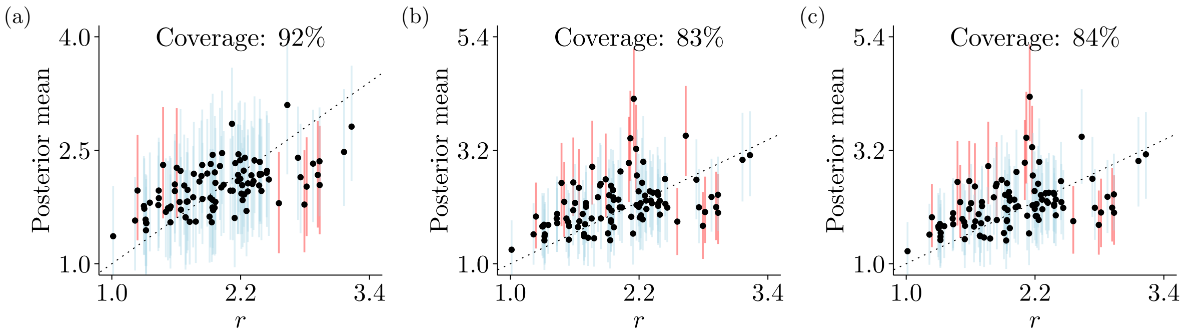


Supplementary Figure 15. Posterior distributions (black dots denoting posterior means, blue lines the 95%-HPD intervals) about $r$ against true (simulated) values (from 100 simulated data sets). In this model, four characters were used, all sharing the same $r$. (a) BEAST 2 sampling $\boldsymbol{\rho}$. (b) BEAST 2 using $\boldsymbol{\rho}^*$. (c) MCMCTree using $\boldsymbol{\rho}^*$.

#### 6.8 Validating with empirical data set

We carried out the same fixed-tree analyses in BEAST 2 and MCMCTree while matching the model used in Álvarez–Carretero et al. (2019), the first with only morphological data, and the second with both molecular and morphological data. 95%-HPDs on node times (Supplementary Fig. 16) and on most parameters (Supplementary Fig. S10 and Table S10) closely matched. The only parameters whose posterior distributions only moderately or weakly overlapped were the gamma distribution shape parameters for the molecular substitution rates. Parameter descriptions can be found in Table S14.

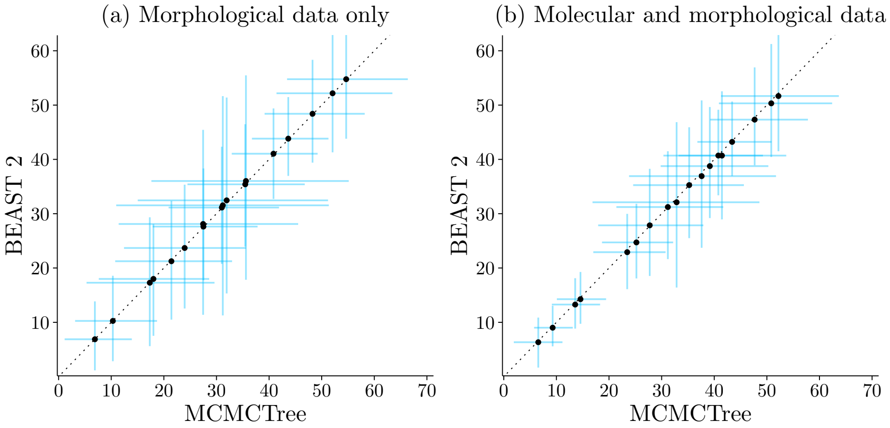


Supplementary Figure 16. Carnivora divergence time 95%-HPDs (blue lines) and posterior means (black dots) from MCMCTree and BEAST 2 analyses of (a) morphological data alone, and (b) both molecular and morphological data.

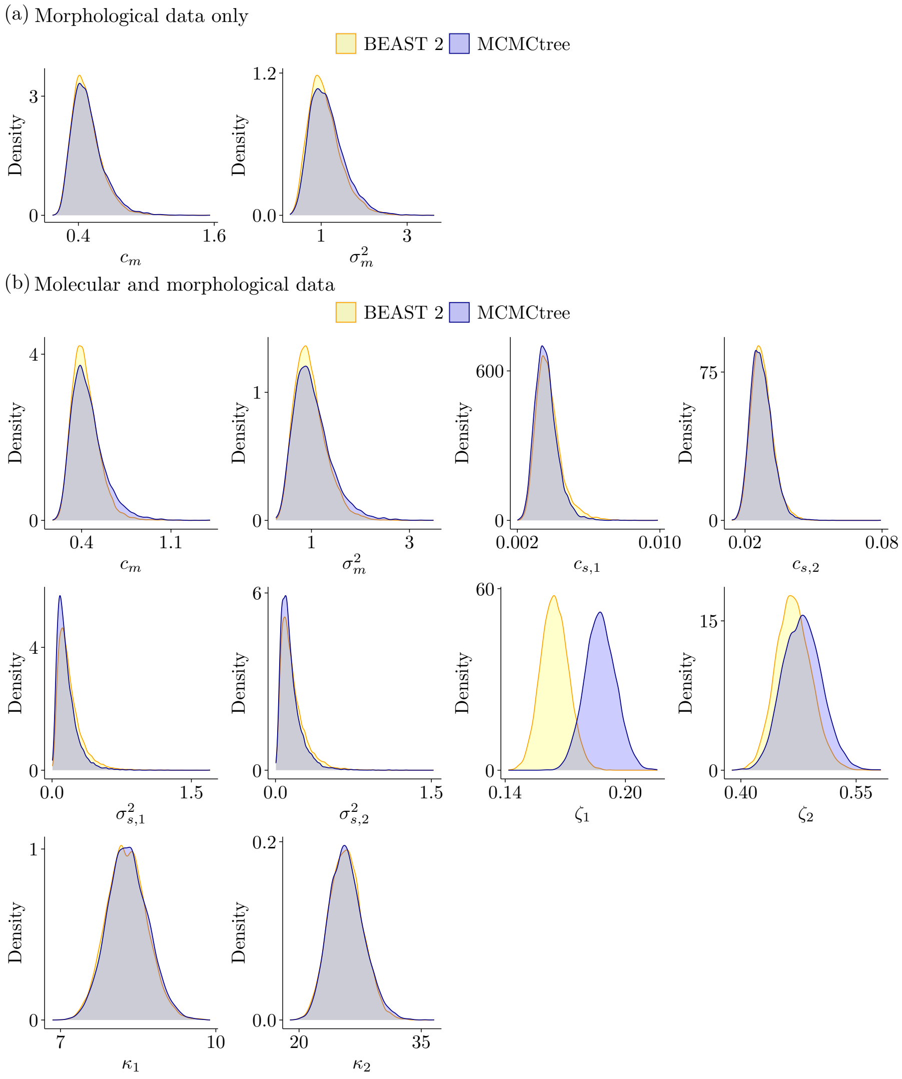


Supplementary Figure 17. Parameter posterior distributions from BEAST 2 (yellow) and MCMCTree (blue) analyses of (a) morphological data alone, and (b) both molecular and morphological data.

### 7. Carnivora empirical analyses

#### 7.1 Carnivora data set

Table S11 summarizes the data used from the 27 species analysed in the present work. Of these 27 species, 11 are extinct and 16 extant species. We used fossil midpoint ages in all our analyses. Molecular sequences of 12 mitochondrial genes were downloaded from NCBI (accession numbers are shown in Table S12), adding up to a total of 10,980. Continuous morphological data consisted of 29 cranium landmarks obtained from Álvarez–Carretero et al., 2019, with each three-dimensional landmark decomposed into three continuous characters, to a total of 87 characters. Discrete morphological characters (a total of 227) were retrieved from (Barrett et al., 2021), and included cranial, basicranial and dental morphological features. Of these characters, 124 were represented by three states, and the remaining by four states.

Table S10. Posterior distribution summaries in Carnivora fixed-tree analyses.

| | BEAST 2 | | MCMCTree | |
|---|---|---|---|---|
| Parameter | 95%-HPD | Mean | 95%-HPD | Mean |
| | **Morphological data only** | | | |
| $c_m$ | [0.2587, 0.7433] | 0.4759 | [0.2585, 0.7708] | 0.4856 |
| $\sigma_m^2$ | [0.4322, 1.851] | 1.0895 | [0.5019, 1.9874] | 1.1447 |
| | **Molecular and morphological data** | | | |
| $c_m$ | [0.2492, 0.6499] | 0.4362 | [0.2395, 0.7152] | 0.4518 |
| $c_s^1$ | [0.0025, 0.0052] | 0.0038 | [0.0025, 0.0049] | 0.0036 |
| $c_s^2$ | [0.0187, 0.038] | 0.0277 | [0.0189, 0.0368] | 0.0273 |
| $\sigma_m^2$ | [0.4198, 1.6088] | 0.9777 | [0.4034, 1.7879] | 1.027 |
| $\sigma_{s1}^2$ | [0.0175, 0.4637] | 0.1955 | [0.0129, 0.3697] | 0.1573 |
| $\sigma_{s2}^2$ | [0.0174, 0.4206] | 0.169 | [0.0195, 0.355] | 0.1478 |
| $\kappa_1$ | [7.5536, 9.063] | 8.282 | [7.5416, 9.0633] | 8.3035 |
| $\kappa_2$ | [21.8494, 29.6859] | 25.7317 | [21.825, 29.9932] | 25.8418 |
| $\zeta_1$ | [0.1517, 0.1778] | 0.1648 | [0.1721, 0.2022] | 0.1875 |
| $\zeta_2$ | [0.4249, 0.5137] | 0.4691 | [0.4322, 0.5295] | 0.4801 |

Table S11. List of species, data types and fossil ages used in Carnivora empirical analyses.

| No. | Taxon ( † indicates extinct taxa) | Molecular | Continuous | Discrete | Specimen midpoint age |
|---|---|---|---|---|---|
| 1 | *Aelurodon ferox* † | | ✓ | | 13.135 |
| 2 | *Ailurus fulgens* | ✓ | ✓ | ✓ | 0.0 |
| 3 | *Canis dirus* † | | ✓ | | 0.0285 |
| 4 | *Canis latrans* | ✓ | | ✓ | 0.0 |
| 5 | *Canis lupus* | ✓ | ✓ | | 0.0 |
| 6 | *Cerdocyon thous* | ✓ | ✓ | | 0.0 |
| 7 | *Cuon alpinus* | ✓ | ✓ | | 0.0 |
| 8 | *Echinosorex gymnura* | ✓ | ✓ | | 0.0 |
| 9 | *Enhydrocyon pahinsintewakpa* † | | ✓ | | 28.55 |
| 10 | *Epicyon haydeni* † | | ✓ | | 11.95 |
| 11 | *Erinaceus europaeus* | ✓ | | | 0.0 |
| 12 | *Hesperocyon gregarius* † | | ✓ | ✓ | 35.55 |
| 13 | *Hyaena hyaena* | ✓ | | ✓ | 0.0 |
| 14 | *Hyaenictitherium wongii* † | | ✓ | | 6.65 |
| 15 | *Nandinia binotata* | ✓ | ✓ | ✓ | 0.0 |
| 16 | *Otarocyon macdonaldi* † | | | ✓ | 33.9 |
| 17 | *Otocyon megalotis* | ✓ | ✓ | | 0.0 |
| 18 | *Panthera leo* | ✓ | | ✓ | 0.0 |
| 19 | *Paradoxurus hermaphroditus* | ✓ | ✓ | | 0.0 |
| 20 | *Paraenhydrocyon josephi* † | | ✓ | | 25.615 |
| 21 | *Puma concolor* | ✓ | | ✓ | 0.0 |
| 22 | *Smilodon fatalis* † | | ✓ | ✓ | 0.0285 |
| 23 | *Smilodon populator* † | ✓ | | | 0.39 |
| 24 | *Speothos venaticus* | ✓ | ✓ | | 0.0 |
| 25 | *Tomarctus hippophaga* † | | ✓ | | 14.785 |
| 26 | *Ursus americanus* | ✓ | ✓ | ✓ | 0.0 |
| 27 | *Vulpes vulpes* | ✓ | ✓ | | 0.0 |

Table S12. Mitochondrial genes (NCBI accession numbers) used in Carnivora empirical analyses.

| Species | ATP6 | ATP8 | COX1 | COX2 | COX3 | CYTB | NADH1 | NADH2 | NADH3 | NADH4L | NADH4 | NADH5 |
|---|---|---|---|---|---|---|---|---|---|---|---|---|
| Ailurus fulgens | NC_011124<br>.1:7952-8632 | NC_011124<br>.1:7791-7994 | NC_011124<br>.1:5350-6894 | NC_011124<br>.1:7034-7717 | NC_011124<br>.1:8632-9415 | NC_011124<br>.1:14171-15310 | NC_011124<br>.1:2755-3711 | NC_011124<br>.1:3920-4963 | NC_011124<br>.1:9485-9830 | NC_011124<br>.1:9901-10197 | NC_011124<br>.1:10191-11568 | NC_011124<br>.1:11766-13586 |
| Canis latrans | NC_008093<br>.1:7965-8645 | NC_008093<br>.1:7804-8007 | NC_008093<br>.1:5350-6894 | NC_008093<br>.1:7035-7718 | NC_008093<br>.1:8645-9428 | NC_008093<br>.1:14185-15324 | NC_008093<br>.1:2748-3704 | NC_008093<br>.1:3915-4958 | NC_008093<br>.1:9497-9843 | NC_008093<br>.1:9913-10209 | NC_008093<br>.1:10203-11580 | NC_008093<br>.1:11780-13600 |
| Canis lycaon | JF342907<br>.3:7965-8645 | JF342907<br>.3:7804-8007 | JF342907<br>.3:5350-6894 | JF342907.3<br>:7035-7718 | JF342907<br>.3:8645-9428 | 14778 | JF342907<br>.3:2748-3703 | JF342907<br>.3:3915-4956 | JF342907<br>.3:9497-9842 | JF342907<br>.3:9913-10209 | JF342907<br>.3:10203-11580 | JF342907<br>.3:11780-13600 |
| Cerdocyon thous | NA | NA | KT236271<br>.1 | NA | NA | KU253530<br>.1 | NA | NA | NA | NA | NA | NA |
| Cuon alpinus | NC_013445<br>.1:7963-8643 | NC_013445<br>.1:7802-8005 | NC_013445<br>.1:5348-6892 | NC_013445<br>.1:7033-7716 | NC_013445<br>.1:8643-9426 | NC_013445<br>.1:14185-15324 | NC_013445<br>.1:2751-3706 | NC_013445<br>.1:3891-4958 | NC_013445<br>.1:9495-9840 | NC_013445<br>.1:9913-10209 | NC_013445<br>.1:10203-11580 | NC_013445<br>.1:11780-13600 |
| Echinosorex gymnura | NC_002808<br>.1:7960-8640 | NC_002808<br>.1:7799-7990 | NC_002808<br>.1:5351-6895 | NC_002808<br>.1:7041-7724 | NC_002808<br>.1:8640-9424 | NC_002808<br>.1:14202-15342 | NC_002808<br>.1:2747-3703 | NC_002808<br>.1:3911-4954 | NC_002808<br>.1:9492-9838 | NC_002808<br>.1:9910-10206 | NC_002808<br>.1:10200-11580 | NC_002808<br>.1:11782-13602 |
| Erinaceus europaeus | NC_002080.2:7947-8627 | NC_002080.2:7786-7992 | NC_002080.2:5346-6890 | NC_002080.2:7034-7717 | NC_002080.2:8627-9410 | NC_002080.2:14182-15321 | NC_002080.2:2745-3698 | NC_002080.2:3909-4952 | NC_002080.2:9485-9829 | NC_002080.2:9901-10197 | NC_002080.2:10191-11568 | NC_002080.2:11769-13580 |
| Hyaena hyaena | NC_020669<br>.1:8734-9414 | NC_020669<br>.1:8573-8776 | NC_020669<br>.1:6130-7674 | NA | NC_020669<br>.1:9414-10197 | NC_020669<br>.1:14952-16091 | NC_020669<br>.1:3531-4486 | NA | NC_020669<br>.1:10267-10613 | NC_020669<br>.1:10681-10977 | NC_020669<br>.1:10971-12348 | NC_020669<br>.1:12548-14368 |
| Nandinia binotata | NC_024567<br>.1:7955-8635 | NC_024567<br>.1:7794-7997 | NC_024567<br>.1:5353-6897 | NC_024567<br>.1:7038-7721 | NC_024567<br>.1:8635-9418 | NC_024567<br>.1:14174-15313 | NC_024567<br>.1:2752-3707 | NC_024567<br>.1:3918-4959 | NC_024567<br>.1:9488-9834 | NC_024567<br>.1:9904-10200 | NC_024567<br>.1:10194-11571 | NC_024567<br>.1:c14101-13574 |
| Otocyon megalotis | NA | NA | AF028198<br>.1 | AF028222<br>.1 | NA | AF028150<br>.1 | NA | NA | NA | NA | NA | NA |
| Panthera leo | NC_028302<br>.1:8636-9316 | NC_028302<br>.1:8475-8678 | NC_028302<br>.1:6033-7577 | NC_028302<br>.1:7719-8402 | NC_028302<br>.1:9316-10099 | NC_028302<br>.1:14855-15994 | NC_028302<br>.1:3433-4388 | NC_028302<br>.1:4599-5640 | NC_028302<br>.1:10169-10515 | NC_028302<br>.1:10585-10881 | NC_028302<br>.1:10875-12252 | NC_028302<br>.1:12451-14271 |
| Paradoxurus hermaphroditus | NA | NA | KC894743<br>.1 | NA | NA | AB075975<br>.1 | NA | AY170056<br>.1 | NA | NA | NA | AB075977<br>.1 |
| Puma concolor | NC_016470<br>.1:8866-9546 | NC_016470<br>.1:8705-8908 | NC_016470<br>.1:6263-7807 | NC_016470<br>.1:7949-8632 | NC_016470<br>.1:9546-10329 | NC_016470<br>.1:15085-16224 | NC_016470<br>.1:3660-4615 | NC_016470<br>.1:4826-5867 | NC_016470<br>.1:10399-10745 | NC_016470<br>.1:10815-11111 | NC_016470<br>.1:11105-12482 | NC_016470<br>.1:12681-14501 |
| Smilodon populator | MF871700<br>.1:7966-8646 | DQ097167<br>.1 | MF871700<br>.1:5361-6759 | MF871700<br>.1:7048-7731 | MF871700<br>.1:8646-9429 | DQ097174<br>.1 | MF871700<br>.1:2761-3716 | MF871700<br>.1:3927-4968 | MF871700<br>.1:9499-9845 | MF871700<br>.1:10042-10211 | MF871700<br>.1:10205-11582 | DQ097169<br>.1 |
| Speothos venaticus | KT448285<br>.1:7921-8601 | KT448285<br>.1:7760-7963 | KT448285<br>.1:5306-6850 | KT448285<br>.1:6991-7674 | KT448285<br>.1:8601-9384 | KT448285<br>.1:14141-15280 | KT448285<br>.1:2702-3658 | KT448285<br>.1:3869-4912 | KT448285<br>.1:9453-9798 | KT448285<br>.1:9869-10165 | KT448285<br>.1:10159-11536 | KT448285<br>.1:11742-13556 |
| Ursus americanus | NC_003426<br>.1:8893-9573 | NC_003426<br>.1:8732-8935 | NC_003426<br>.1:6291-7835 | NC_003426<br>.1:7975-8658 | NC_003426<br>.1:9573-10356 | NC_003426<br>.1:15113-16252 | NC_003426<br>.1:3700-4656 | NC_003426<br>.1:4864-5907 | NC_003426<br>.1:10426-10772 | NC_003426<br>.1:10842-11138 | NC_003426<br>.1:11132-12509 | NC_003426<br>.1:12708-14536 |
| Vulpes vulpes | NC_008434<br>.1:7965-8645 | NC_008434<br>.1:7804-8007 | NC_008434<br>.1:5350-6894 | NC_008434<br>.1:7035-7718 | NC_008434<br>.1:8645-9428 | NC_008434<br>.1:14185-15324 | NC_008434<br>.1:2749-3705 | NC_008434<br>.1:3916-4959 | NC_008434<br>.1:9497-9843 | NC_008434<br>.1:9913-10209 | NC_008434<br>.1:10203-11581 | NC_008434<br>.1:11780-13600 |

#### 7.2 Integrative model used in Carnivora TED

The general structure of our integrative model can be represented by a probabilistic graphical model (Fig. 18). With the exception of the discrete morphology model component and the partial or absent topological constraints (in some of our analyses), we matched the model in Álvarez–Carretero et al. (2019), which includes molecular and continuous morphological clock models, and all hyperprior distributions.

We used the same partitioning scheme for the molecular data – two partitions comprised by 7,320 sites from first and second codon positions, and 3,660 sites from third codon positions, respectively – as well as the same substitution model (HKY+Γ with five discrete categories; Hasegawa et al., 1985; Yang, 1994). Equilibrium nucleotide frequencies were set to their empirical values.

Continuous morphological evolution was modeled with a phylogenetic BM model, with all 87 superimposed, standardized characters (see above) sharing the same relative evolutionary rate ($\boldsymbol{r} = \mathbf{1}$; Fig. 18). We estimated the landmark root values ($\boldsymbol{y_0}$) using maximum-likelihood, obtaining $\widehat{\boldsymbol{y_0}}$ as a byproduct of pruning as done in Álvarez–Carretero et al., 2019. We did so first because this approach allows for direct comparison with results from that study, and second because it is not immediately obvious how to choose a prior for $\boldsymbol{y_0}$. Using $\widehat{\boldsymbol{y_0}}$ is analogous to employing empirical nucleotide frequencies as the equilibrium distribution under molecular substitution models. We note that both assuming the same evolutionary rates across characters and calculating $\widehat{\boldsymbol{y_0}}$ are not requirements of our implementation; $\boldsymbol{r} = \mathbf{1}$ can be relaxed, and $\boldsymbol{y_0}$ can be sampled during MCMC. Finally, uncorrelated log-normal relaxed clock models were used for both molecular and morphological evolutionary rates.

Discrete morphological evolution was modeled with a Lewis' Mkv+Γ model (Lewis, 2001), with three discrete categories, the best partitioning found by Barrett et al. (2021).

#### 7.3 Analyses estimating both divergence times and topology

Table S14 details all priors used in our empirical this set of analyses, many of which are the same as those used in Álvarez–Carretero et al. (2019) and Barrett et al. (2021) (we point the reader to those reference for more detail on prior choice).

Birth and death rates from the BDSS and FBD tree models were assigned priors centered at the values used in Álvarez–Carretero et al. (2019), with variance of 0.5. We specified a log-normal prior distribution for the sampling rate, $\psi$, the 95%-HPD of which covers the [0.00805, 0.0417] interval. This is a wide prior that includes values 9/300 to 9/600 (our data set includes 9 fossils and the Carnivora order is thought to have between 300 to 600 species; Kruuk, 2001; Duszynski et al., 2018). We used a beta distribution for $\rho$ (the probability of sampling extant species) as this parameter is bounded by 0.0 and 1.0, with approximately 90% of the probability density being assigned to values $< 0.2$.

#### 7.4 MCMC and model comparison

Inference in all six empirical model configurations was carried out with MCMC as implemented in BEAST 2. For each of the six analyses, at least three independent MCMC chains were run in parallel, were inspected for convergence, and were then combined (four, five and six chains were run for BDSS-1, FBD-1 and FBD-3), each with 100 million states being logged at every 2,000. Chain convergence was evaluated by visual inspection of chain traces and by verifying that effective sample sizes exceeded 200.

Model comparison was conducted by computing posterior odds between pairs of model configurations, out of the six possibilities described in the main text. The posterior odd between two models $\mathcal{M}_0$ and $\mathcal{M}_1$ is given by:

Table S13. List of initial values of estimated parameters used in Carnivora empirical analyses.

| Analysis | FBD-1 | | | FBD-2 | | | FBD-3 | | | BDSS-1 | | | BDSS-2 | | | BDSS-3 | | |
|---|---|---|---|---|---|---|---|---|---|---|---|---|---|---|---|---|---|---|
| Parameter | Chain 1 | Chain 2 | Chain 3 | Chain 1 | Chain 2 | Chain 3 | Chain 1 | Chain 2 | Chain 3 | Chain 1 | Chain 2 | Chain 3 | Chain 1 | Chain 2 | Chain 3 | Chain 1 | Chain 2 | Chain 3 |
| $\lambda$ | 1.62 | 1.29 | 0.66 | 2.04 | 1.81 | 0.75 | 1.04 | 1.05 | 1.22 | 1.27 | 1.75 | 1.15 | 1.88 | 0.64 | 0.74 | 0.96 | 1.13 | 2.11 |
| $\mu$ | 0.61 | 0.58 | 0.58 | 0.58 | 1.20 | 0.64 | 0.44 | 1.04 | 1.02 | 0.75 | 1.37 | 0.41 | 1.41 | 0.41 | 0.47 | 0.91 | 0.70 | 1.21 |
| $\psi$ | 0.01 | 0.02 | 0.03 | 0.02 | 0.02 | 0.01 | 0.02 | 0.02 | 0.02 | 0.03 | 0.01 | 0.01 | 0.01 | 0.02 | 0.04 | 0.02 | 0.06 | 0.03 |
| $\rho$ | 0.01 | 0.02 | 0.10 | 0.01 | 0.02 | 0.06 | 0.06 | 0.03 | 0.03 | 0.15 | 0.09 | 0.03 | 0.16 | 0.24 | 0.05 | 0.02 | 0.11 | 0.06 |
| $\kappa_1$ | 1.73 | 0.14 | 1.79 | 2.83 | 1.27 | 2.38 | 1.84 | 0.35 | 2.36 | 1.67 | 0.34 | 0.88 | 1.82 | 0.24 | 0.36 | 0.13 | 0.73 | 0.03 |
| $\sigma_{s1}$ | 1.16 | 1.07 | 0.83 | 1.36 | 1.10 | 0.74 | 0.66 | 0.55 | 0.49 | 0.57 | 0.94 | 0.65 | 0.98 | 0.96 | 0.69 | 0.83 | 1.02 | 1.56 |
| $\boldsymbol{b}_s^1$ | 0.38 | 1.41 | 0.11 | 0.82 | 0.41 | 0.46 | 1.18 | 0.56 | 1.24 | 0.59 | 0.53 | 0.93 | 0.74 | 0.97 | 0.82 | 2.22 | 4.24 | 0.28 |
| $c_s^1$ | 0.21 | 0.29 | 0.18 | 0.11 | 0.20 | 0.19 | 0.06 | 0.16 | 0.29 | 0.08 | 0.06 | 0.10 | 0.06 | 0.10 | 0.03 | 0.27 | 0.15 | 0.43 |
| $\zeta_1$ | 0.04 | 1.51 | 0.13 | 2.99 | 0.49 | 0.31 | 0.24 | 1.54 | 1.94 | 1.18 | 1.98 | 1.29 | 0.27 | 0.77 | 1.61 | 0.98 | 1.42 | 0.11 |
| $\kappa_2$ | 0.38 | 2.38 | 0.02 | 2.26 | 2.29 | 1.45 | 0.55 | 1.93 | 2.98 | 0.56 | 0.75 | 0.14 | 0.10 | 0.34 | 0.19 | 3.21 | 0.85 | 0.76 |
| $\sigma_{s2}$ | 0.59 | 1.19 | 0.87 | 0.68 | 1.47 | 0.86 | 0.70 | 1.10 | 0.81 | 1.09 | 0.97 | 1.13 | 1.15 | 1.31 | 0.67 | 1.09 | 0.83 | 1.00 |
| $\boldsymbol{b}_s^2$ | 0.79 | 1.87 | 0.98 | 0.70 | 0.17 | 0.71 | 0.94 | 0.61 | 0.63 | 0.25 | 1.94 | 0.46 | 1.63 | 0.37 | 1.40 | 2.09 | 0.68 | 1.66 |
| $c_s^2$ | 0.19 | 0.33 | 0.14 | 0.26 | 0.36 | 0.16 | 0.61 | 0.04 | 0.06 | 0.09 | 0.10 | 0.18 | 0.21 | 0.13 | 0.28 | 0.21 | 0.08 | 0.31 |
| $\zeta_2$ | 1.66 | 0.46 | 4.98 | 2.06 | 1.04 | 0.20 | 0.91 | 0.75 | 0.12 | 0.35 | 0.26 | 0.30 | 0.07 | 2.89 | 1.16 | 0.01 | 1.16 | 0.16 |
| $\sigma_m$ | 0.93 | 0.50 | 0.59 | 0.27 | 0.90 | 1.07 | 0.63 | 0.69 | 1.25 | 1.07 | 1.19 | 0.38 | 0.89 | 1.31 | 1.05 | 0.40 | 1.05 | 0.95 |
| $\boldsymbol{b}_m$ | 0.64 | 0.55 | 0.47 | 1.28 | 0.97 | 0.67 | 0.19 | 0.70 | 0.16 | 0.44 | 0.29 | 1.17 | 0.46 | 1.44 | 0.58 | 1.27 | 3.58 | 0.91 |
| $c_m$ | 0.12 | 0.21 | 0.25 | 0.43 | 0.46 | 0.11 | 0.23 | 0.16 | 0.05 | 0.26 | 0.01 | 0.61 | 0.33 | 0.12 | 0.05 | 0.07 | 0.08 | 0.03 |
| $\zeta_d^2$ | 1.05 | 0.45 | 0.40 | 0.58 | 0.86 | 0.91 | 0.02 | 0.42 | 0.31 | 0.18 | 1.27 | 1.73 | 0.76 | 1.08 | 0.70 | 0.97 | 2.71 | 1.31 |
| $\zeta_d^3$ | 0.65 | 1.70 | 0.72 | 0.09 | 1.74 | 0.39 | 0.25 | 0.56 | 3.15 | 0.99 | 0.89 | 2.54 | 1.06 | 1.91 | 1.60 | 0.21 | 2.09 | 0.19 |
| $\zeta_d^4$ | 0.06 | 0.94 | 1.01 | 1.58 | 0.35 | 0.36 | 1.00 | 2.32 | 0.76 | 1.62 | 1.39 | 0.43 | 0.19 | 0.33 | 0.67 | 0.19 | 0.39 | 2.41 |
| $c_d$ | 1.45 | 0.51 | 1.49 | 0.78 | 0.33 | 0.54 | 0.57 | 1.26 | 0.76 | 1.35 | 1.03 | 1.41 | 1.50 | 0.61 | 1.16 | 1.63 | 0.55 | 0.72 |
| $\sigma_d$ | 0.03 | 0.06 | 0.56 | 0.02 | 0.27 | 0.01 | 0.05 | 0.12 | 0.03 | 6.10e-04 | 0.11 | 0.03 | 0.07 | 8.43e-05 | 0.03 | 1.66e-03 | 0.20 | 0.12 |
| $\boldsymbol{b}_d$ | 1.01 | 0.99 | 0.37 | 0.97 | 0.85 | 0.98 | 1.05 | 1.12 | 0.97 | 1.00 | 0.85 | 1.01 | 0.97 | 1.00 | 0.98 | 1.00 | 0.82 | 0.92 |

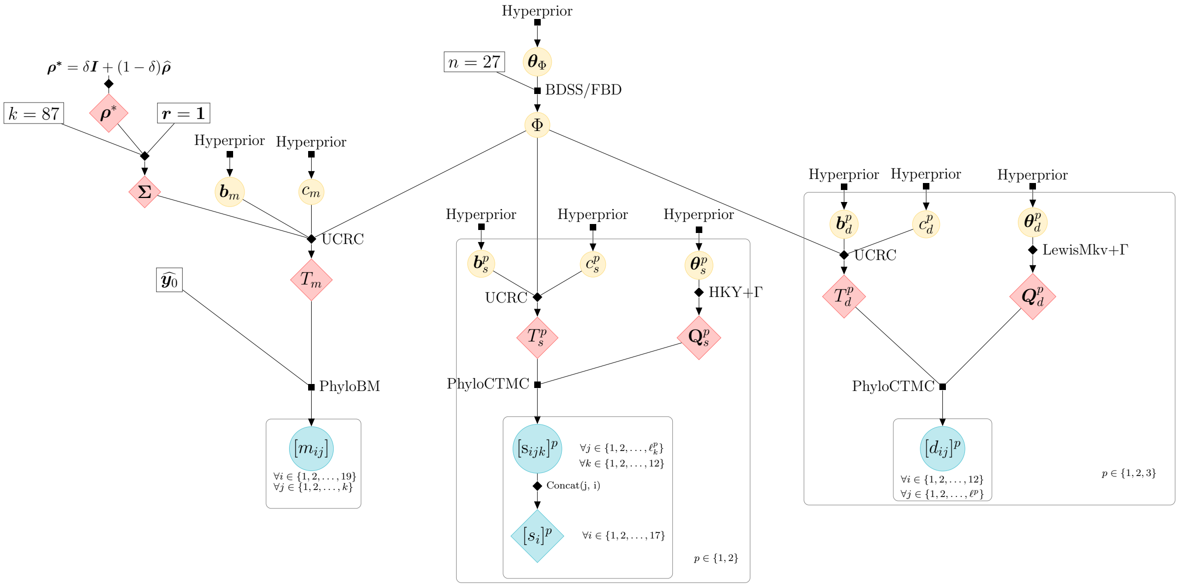


Supplementary Figure 18. .Probabilistic graphical model used in the analyses of the Carnivora data set, composed of 12 concatenated mitochondrial genes ($L = 12$) split into two partitions ($p \in \{1, 2\}$ for molecular data, $p \in \{1, 2, 3\}$ for discrete morphological data), and 87 continuous characters ($k = 87$) from 19 Carnivora species ($n = 19$). Filled squares denote sampling distributions, diamonds denote deterministic functions (black) and their outputs (red), and circles denote random variables that do not (yellow) or that do have data associated to them (blue). Bold symbols represent vectors or matrices, otherwise they are scalars. $\mathbf{M} = [m_{ij}]$ and $\mathbf{D} = [d_{ij}]$. $l_k^p$ inside the plate for the molecular data denotes the length of the $k$-th locus in the $p$-th molecular partition. Other symbols are defined in the main text. 'PhyloBM' and 'PhyloCTMC' stand for phylogenetic Brownian motion and phylogenetic continuous-time Markov chain models, and 'UCRC' for uncorrelated relaxed clock models. The sampling distribution for the species tree ($\Phi$) was either the BDSS or FBD model (see Methods section and Table 1 in the main manuscript). $\mathrm{Q}_{\boldsymbol{s}}^{\boldsymbol{p}}$ and $\mathrm{Q}_{\boldsymbol{d}}^{\boldsymbol{p}}$ represent the instantaneous rate matrices under the substitution and discrete-morphology models, respectively, for a partition $p$. "Concat($\cdot$)" denotes the deterministic task of concatenating all sites from all loci within one of the two partitions. For the sake of clarity, $\boldsymbol{\theta} = \{\boldsymbol{\theta}_\Phi, \boldsymbol{\theta}_s^p\}$ and 'Hyperprior' encompass all parameters and priors not explicitly shown in the graphical model. For more details, please see subsection "Graphical representation of Bayesian networks" above.

$$\underbrace{\frac{f(\mathcal{M}_0|D)}{f(\mathcal{M}_1|D)}}_{\text{Posterior odds}} = \underbrace{\frac{f(D|\mathcal{M}_0)}{f(D|\mathcal{M}_1)}}_{\text{Bayes factor}} \times \underbrace{\frac{f(\mathcal{M}_0)}{f(\mathcal{M}_1)}}_{\text{Prior odds}} . \tag{42}$$

By assuming both models have the same prior probability (i.e., prior odds = 1), the posterior odds reduces to the ratio of the marginal likelihoods $\mathcal{Z}_0 = f(D|\mathcal{M}_0)$ and $\mathcal{Z}_1 = f(D|\mathcal{M}_1)$. The marginal likelihood of a model is the probability of observing the data under that model, and so it measures a model's goodness-of-fit. The ratio of two model's marginal likelihoods, known as the Bayes factor (BF), thus allows one to quantify which of the two has the best fit. A log-BF $> 0$, for example, indicates $\mathcal{M}_0$ fits the data better; when $> 2$, the log-BF suggests $\mathcal{M}_0$ is decisively supported over $\mathcal{M}_1$ (Kass and Raftery, 1995).

Many Bayesian methods exist for calculating marginal likelihoods, such as the harmonic mean approach (Newton and Raftery, 1994), thermodynamic integration (Lartillot and Philippe, 2006), steppingstone sampling (Xie et al., 2011), generalized steppingstone sampling (Fan et al., 2011), and nested sampling (NS; Russel et al., 2019). We chose NS

Table S14. Summary of prior distributions used in Carnivora data set analysis. Gamma$(\alpha, \beta)$ denotes a gamma distribution with shape parameter $\alpha$ and inverse scale parameter $\beta$; LN($\mu$, $\sigma$, T) denotes a log-normal distribution with real-space mean $\mu$ and log-space standard deviation $\sigma$; LN($\mu$, $\sigma$, F) denotes a log-normal distribution with log-space mean $\mu$ and log-space standard deviation $\sigma$; Beta($\alpha$, $\beta$) denotes a beta distribution with shape parameters $\alpha$ and $\beta$'; $\text{U}_s(l, u)$ denotes a uniform distribution with soft lower bound $l$ and soft upper bound $u$; Exp($\alpha$) denotes an exponential distribution with mean $\alpha$. "s", "d" and "m" subscripts denote parameters from models using molecular sequences, discrete morphology and continuous morphology, respectively.

| Parameter | Description | Prior distribution |
|---|---|---|
| | **BDSS/FBD parameters** | |
| $\lambda$ | Birth rate | LN(1, 0.5, T) |
| $\mu$ | Death rate | LN(1, 0.5, T) |
| $\psi$ | Sampling rate | LN(-4, 0.5, F) |
| $\rho$ | Probability of sampling an extant species | Beta(1, 10) |
| $t_{\text{mrca}}$ | Root age | $\text{U}_s$(37.3, 66.0) |
| | **Substitution model parameters** | |
| $\kappa_1$, $\kappa_2$ | 2 × transition-transversion ratio | Gamma$(1, 1)$ |
| $\zeta_1$, $\zeta_2$ | Shape of discretized gamma (substitution) model | Gamma$(1, 1)$ |
| $\zeta_d^1$, $\zeta_d^2$, $\zeta_d^3$, $\zeta_d^4$ | Shape of discretized gamma (state change) model | Exp(1) |
| | **Clock model parameters** | |
| $\boldsymbol{b}_m$ | Branch-specific rate scaler | LN(1, $\sigma_m^2$) |
| $\boldsymbol{b}_d$ | Branch-specific rate scaler | LN(1, $\sigma_d^2$) |
| $\boldsymbol{b}_s^1$ | Branch-specific rate scaler (1st partition) | LN(1, $\sigma_{s1}^2$) |
| $\boldsymbol{b}_s^2$ | Branch-specific rate scaler (2nd partition) | LN(1, $\sigma_{s2}^2$) |
| $c_m$ | Global clock rate | Gamma$(2, 10)$ |
| $c_d$ | Global clock rate | LN$(1, 1.25)$, $c_d < 2$ |
| $c_s^1$, $c_s^2$ | Global clock rate | Gamma$(2, 10)$ |
| $\sigma_m^2$ | Variance of branch-specific rates | Gamma$(2, 2)$ |
| $\sigma_d^2$ | Variance of branch-specific rates | Gamma$(0.5396, 0.3819)$ |
| $\sigma_{s1}^2$, $\sigma_{s2}^2$ | Variance of branch-specific rates | Gamma$(2, 2)$ |

because it is readily available in BEAST 2, the platform in which we implemented our integrative model, but also because this technique overcomes many shortcomings of other methods. Namely, NS is robust to phylogenetic tree spaces with tree "islands", copes better with convex likelihood functions, requires simpler tuning, dispenses with the burn-in stage, and can have the uncertainty around its estimate calculated in a single run (Russel et al., 2019).

The gist of NS consists of approximating integral (see Fig. 1 in Russel et al., 2019):

$$\mathcal{Z} = \int_0^1 L(\xi) d\xi, \tag{43}$$

where $\xi$ denotes the prior mass, and $L(\xi)$ gives us the likelihood value above which a $\xi$ fraction of prior draws will be. NS allows us to obtain a set of points on the $L(\xi)$ curve that can be used to approximate the integral above via the standard quadrature method.

The procedure involves iteratively drawing these points (with replacement) from a pool of "active points" of fixed size $N$ (i.e., $N$ active points): $\{\boldsymbol{\theta_1}, ..., \boldsymbol{\theta_N}\}$. The first point is drawn from the prior, but then successive draws always consist of the point with the lowest likelihood in the pool. After a point $\boldsymbol{\theta}_i$ is taken, it is replaced with another $\boldsymbol{\theta}$ (drawn from the prior) that has a higher likelihood. This has the effect of progressively constraining the prior from which points are being drawn to areas of higher likelihood – something referred to as dynamic likelihood restriction.

When approximating $\mathcal{Z}$ with NS, the two main tuning parameters are the number of active points $N$ and the "subchain length", which gives us the number of MCMC steps between draws from the pool of active points. Larger $N$ values increase the precision (but are unrelated to the bias) of our $\mathcal{Z}$ estimates, while larger subchain lengths allow draws to be more independent. This latter condition is a requirement for the NS procedure. (The interested reader can find thorough formal treatments of NS in Russel et al., 2019 and Skilling et al., 2006.)

In order to calculate posterior odds for a pair of models (Eq. (42)), we followed the NS protocol described in (Russel et al., 2019). We ran an initial chain with one active point ($N = 1$) and a sub-chain length of 5,000 ($K = 5,000$), and based on the estimated standard deviation of the marginal likelihood, $\sigma_{\mathcal{Z}}$, we updated the number of active points to 55 ($N = 55$) so that $\sigma_{\mathcal{Z}} < 2$. ($\sigma_{\mathcal{Z}}$ is inversely proportional to the square root of $N$ Skilling et al., 2006.) The appropriate sub-chain length was determined by running two chains $\mathcal{A}$ and $\mathcal{B}$ (for each of the two models) with sub-chain lengths of 5,000 and 7,500. A length of 5,000 was deemed sufficient because $|\mathcal{Z}_A - \mathcal{Z}_B| < 2\sqrt{\sigma^2_{\mathcal{Z}_\mathcal{A}} + \sigma^2_{\mathcal{Z}_\mathcal{B}}}$. For each of the models being compared, we thus ran a chain with 55 particles and sub-chain length of 5,000.

##### 7.5 Summary trees from Carnivora analyses

Below, we provide the maximum-clade-credibility (MCC) summary trees from the analyses described in the main text (Supplementary Figs. 19). We also present the MCC summary trees from analyses using just continuous morphological data (Supplementary Figure 20).

##### 7.6 Multidimensional scaling (MDS)

MDS of the 29 cranium landmarks was done on distances calculated for each species $i$-$j$ pair as:

$$d(i,j) = \sqrt{\sum_{k=1}^{29} \left( \sqrt{(x_i^k - x_j^k)^2 + (y_i^k - y_j^k)^2 + (z_i^k - z_j^k)^2} \right)} \tag{44}$$

where $i, j \in \{1, 2, ..., 19\}$ ($i \neq j$), and $x_i^k$, $y_i^k$ and $z_i^k$ represent the three coordinates of the $k$-th landmark for species $i$.

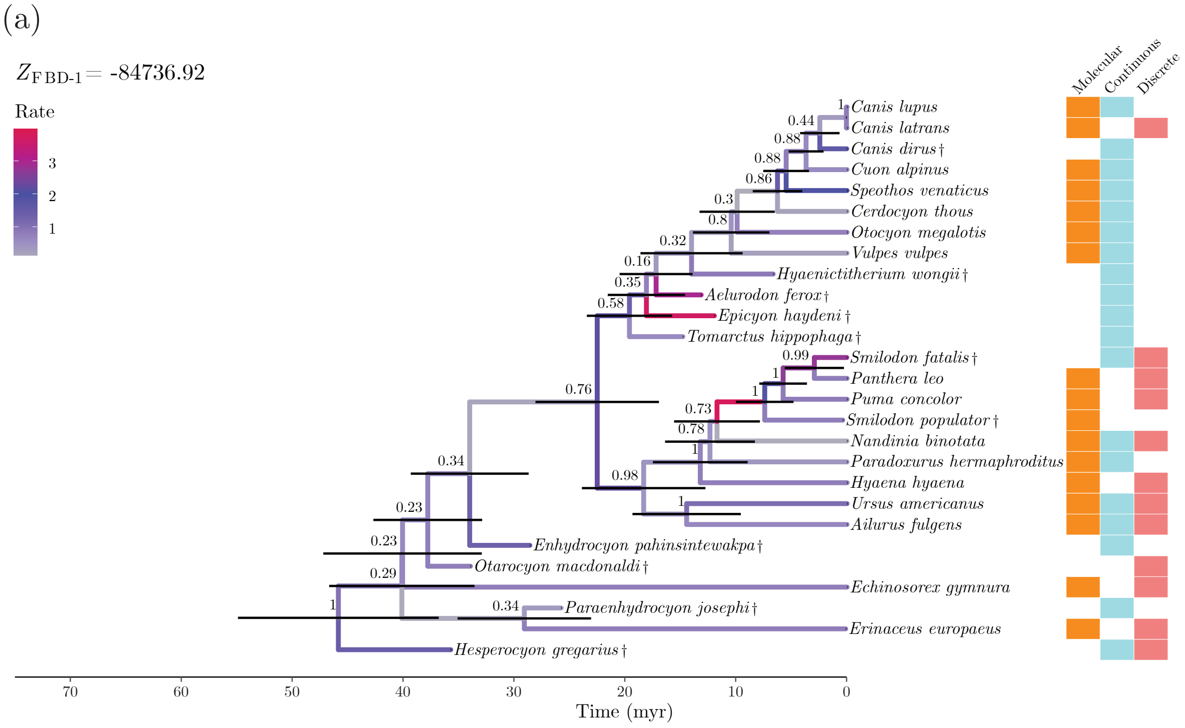


Supplementary Figure 19. Maximum-clade-credibility (MCC) summary trees of Carnivora species. Horizontal bars at internal nodes show the 95% credible intervals for their times. Numbers by internal nodes indicate each clade's posterior probability. (a) Tree estimated in analysis 1.

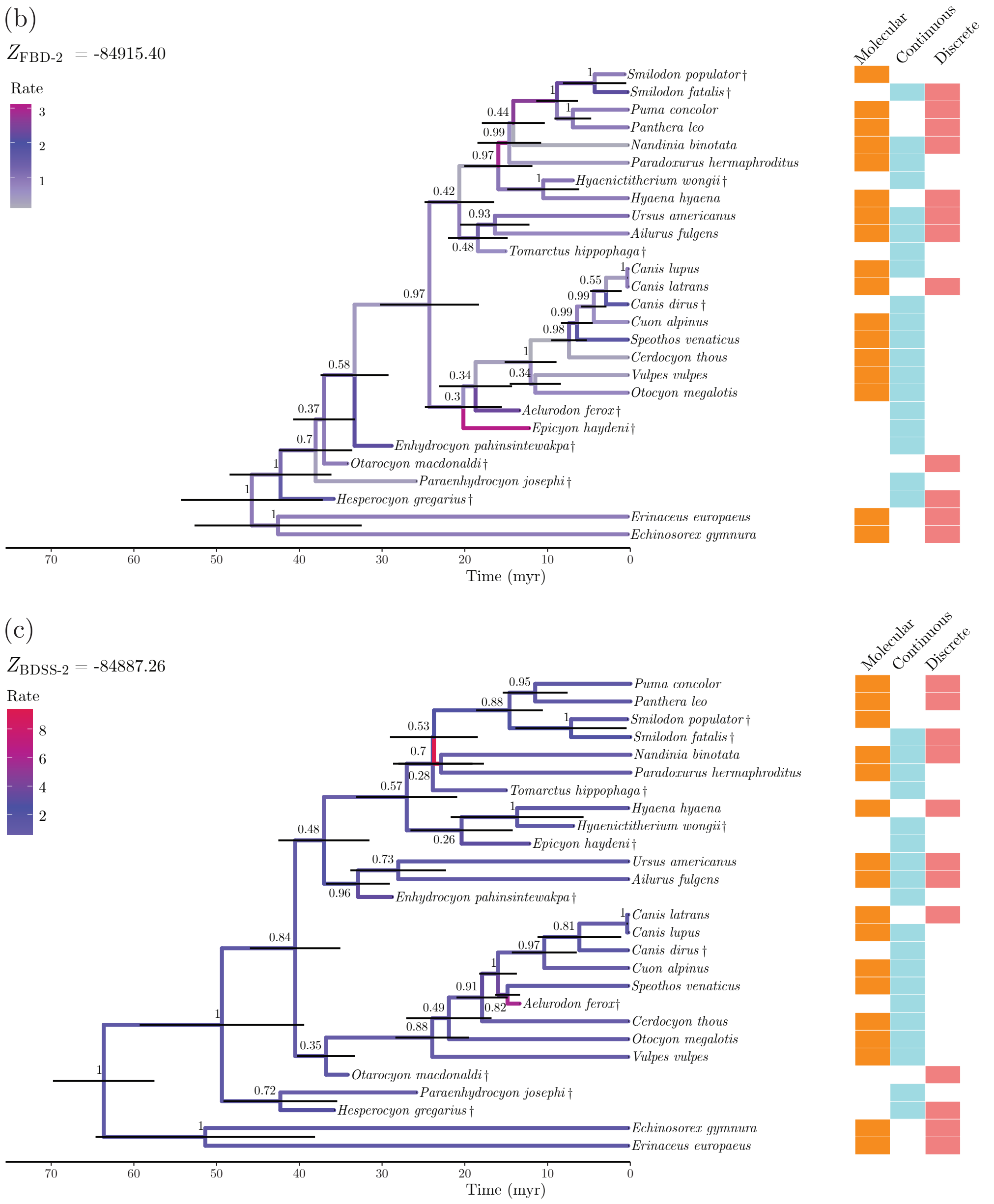


Supplementary Figure 19. Maximum-clade-credibility (MCC) summary trees of Carnivora species (continued). (b) Tree estimated in analysis 2. (c) Tree estimated in analysis 5.

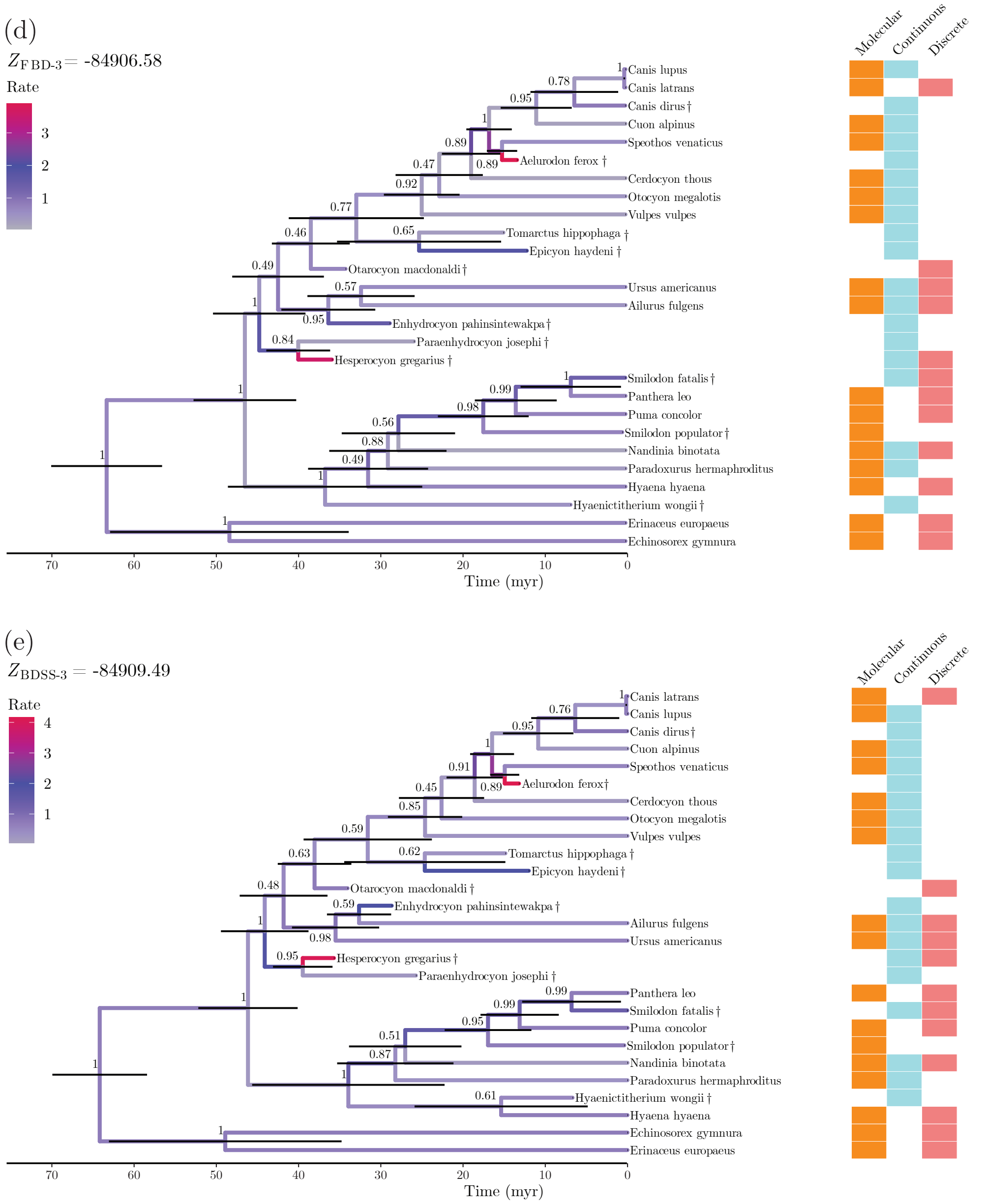


Supplementary Figure 19. Maximum-clade-credibility (MCC) summary trees of Carnivora species (continued). (d) Tree estimated in analysis 3. (e) Tree estimated in analysis 6.

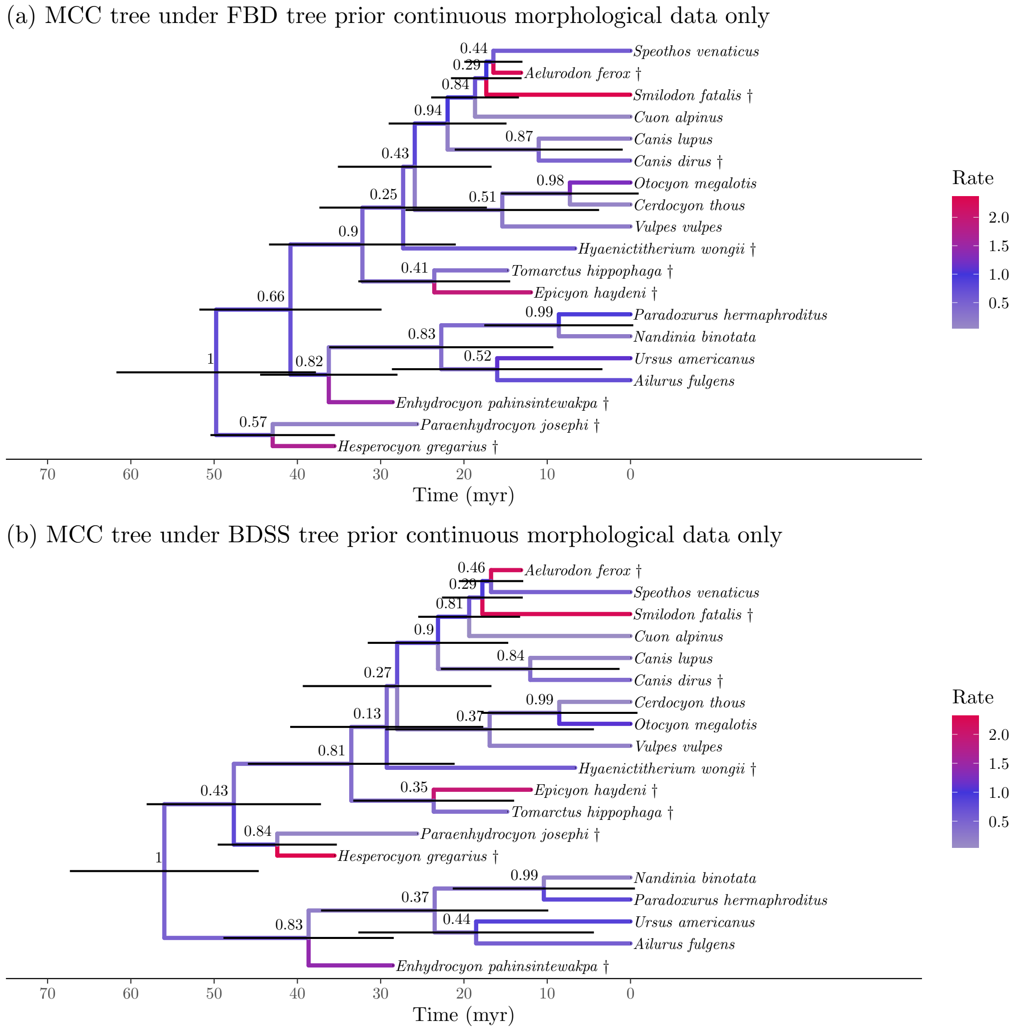


Supplementary Figure 20. Maximum-clade-credibility (MCC) summary trees of Carnivora species. Horizontal bars at internal nodes show the 95% credible intervals for their times. Numbers by internal nodes indicate each clade's posterior probability. (a) Tree estimated with only continuous morphological data, following analysis FBD-1's configuration. (b) Tree estimated with only continuous morphological data, following analysis BDSS-1's configuration.

#### 8. Algorithms

---

**Algorithm 1:** BDSS model direct simulator

---

**Input:**

A set of species $\boldsymbol{S} = \{\mathit{1}, \mathit{2}, \cdots, n\}$;

A set of sampled time corresponding to each species $\boldsymbol{z} = \{z_1, z_2, \cdots, z_n\}$;

Root time (tree height) $t_{\mathrm{mrca}}$;

Model parameters $\boldsymbol{\theta} = \{\lambda, \mu, \psi, \rho\}$

**Output:** A time tree $\Phi = \{\mathcal{T}, \boldsymbol{x}\}$, where $\mathcal{T}$ denotes the tree topology and $\boldsymbol{x}$ the set of internal node times.

**Simulation process:**

$\mathcal{T} \leftarrow$ `ConstructRandomTree(`$\boldsymbol{S}$`)` `// Defined in Algorithm 2`;

$\boldsymbol{x} \leftarrow$ `ProposeNodeTimes(`$\mathcal{T}$, $\boldsymbol{\theta}$, $\boldsymbol{z}$`)` `// Defined in Algorithm 3`;

$\Phi \leftarrow$ `ReconstructTree(`$\mathcal{T}$, $\boldsymbol{x}$`)` `// Defined in Algorithm 4`;

---

**Algorithm 2:** Subroutine `ConstructRandomTree`

---

**Input:** A set of species $\boldsymbol{S}$

$\boldsymbol{C} \leftarrow \boldsymbol{S}$ `// Construct a set of nodes` $\boldsymbol{C}$`, each node from a species in` $\boldsymbol{S}$

**Output:** A tree topology $\mathcal{T}$, rooted at node $(2n-1)$

**begin**

    **for** $i = n + 1$ **to** $2n - 1$ **do**

        $\{a, b\} \leftarrow$ randomly pick two nodes from $\boldsymbol{C}$

        Remove $a$ and $b$ from $\boldsymbol{C}$

        `// Make node` $i$ `as parent of` $a$ `and` $b$

        Parent of $a \leftarrow i$

        Parent of $b \leftarrow i$

        Add node $i$ to $\boldsymbol{C}$

    **end**

**end**

---

**Algorithm 3:** Subroutine `ProposeNodeTimes`

```
Input:
Tree topology 𝒯;
Model parameters θ;
Observed tip times z;
Output: A time tree Φ = {𝒯, x};
begin
    for internal node i ∈ 𝒯 do
        l ← left subtree descending from i;
        z_l ← time of rightmost tip in l;
        r ← right subtree descending from i;
        z_r ← time of leftmost tip in r;
        z_i^* ← max(z_l, z_r);
        Randomly draw p_{z_i^*} from U(0,1);
        x_i ← F'_{X_i}(p_{z_i^*}) // From Eq. (37) (x_i is the age of i);
        if i is root then
            t_mrca ← x_i;
        end
    end
end
```

**Algorithm 4:** Subroutine `ReconstructTree`

```
Input:
Tree topology 𝒯;
A set of internal node times x;
A set of sampled times of tips z;
Output: A time tree Φ = {𝒯, t};
begin
    I ← index of nodes from left to right;
    t ← {z, x};
    NodeReheight(I, t, 1, 2n − 1) // Defined in Algorithm 5;
end
```

**Algorithm 5:** Subroutine `NodeReheight`

**Input:**

A set of node index in the order of from left to right (in-order traversal) $\boldsymbol{I}$;

A set of node times including tips and internal nodes $\boldsymbol{t}$;

Starting index to visit the node index $i$;

Ending index to finish the visit $j$

**begin**

$nm \leftarrow$ node with maximum height among nodes from $\boldsymbol{I}[i]$ to $\boldsymbol{I}[j]$;

$nl \leftarrow$ node with maximum height among nodes from $\boldsymbol{I}[i]$ to $\boldsymbol{I}[nm-1]$;

$nr \leftarrow$ node with maximum height among nodes from $\boldsymbol{I}[nm+1]$ to $\boldsymbol{I}[j]$;

`// Make node` $nm$ `as the parent of` $nl$ `and` $nr$;

parent of $nl \leftarrow nm$;

parent of $nr \leftarrow nm$;

`NodeReheight(`$\boldsymbol{I}$, $\boldsymbol{t}$, $i$, $nm$`)`;

`NodeReheight(`$\boldsymbol{I}$, $\boldsymbol{t}$, $nm$, $j$`)`;

**end**

#### 9. Software used in validation analysis

Table S15 lists the programs and packages we used in our validation and empirical analyses.

Table S15. Programs and packages used in validation and empirical analyses.

| Package | Type | Version | Use | Reference |
|---|---|---|---|---|
| `FossilSim` | R package | 2.1.1 | FBD tree simulation | Barido-Sotani et al. (2019) |
| `PCMBase` | R package | 1.2.10 | Continuous traits simulation | |
| `phangorn` | R package | 2.5.3 | Molecular sequence simulation; calculation of tree distances | Schliep (2011) |
| `phytools` | R package | 0.6-60 | Phylogenetic tree parsing and visualisation | Revell (2012) |
| `Tracer` | Standalone | 1.7.1 | MCMC diagnostics | Rambaut et al. (2018) |
| `sampled-ancestors` | BEAST 2 package | 2.0.2 | FBD model implementation | Gavryushkina et al. (2014b) |
| `BEASTLabs` | BEAST 2 package | 1.8.0 | BEAST utilities (dependency of other packages) | N/A |
| `MCMCTree` | PAML package | 4.9j | Estimation for continuous traits | Yang and Rannala (2006) |
